# Candidate transmission survival genome of *Mycobacterium tuberculosis*

**DOI:** 10.1101/2025.01.30.635747

**Authors:** Saurabh Mishra, Prabhat Ranjan Singh, Xiaoyi Hu, Landys Lopez-Quezada, Adrian Jinich, Robin Jahn, Luc Geurts, Naijian Shen, Michael A. DeJesus, Travis Hartman, Kyu Rhee, Matthew Zimmerman, Veronique Dartois, Richard M. Jones, Xiuju Jiang, Ricardo Almada-Monter, Lydia Bourouiba, Carl Nathan

**Author notes:** Co-senior authors to whom correspondence should be addressed at **Email:** or. co-first authors (Saurabh Mishra, Prabhat Ranjan Singh, Xiaoyi Hu). Current address: National Institute of Pharmaceutical Education and Research (NIPER), Mohali, India (Saurabh Mishra). **Classification:** Biological sciences; Microbiology.

## Abstract

*Mycobacterium tuberculosis* (Mtb), a leading cause of death from infection, completes its life cycle entirely in humans except for transmission through the air. To begin to understand how Mtb survives aerosolization, we mimicked liquid and atmospheric conditions experienced by Mtb before and after exhalation using a model aerosol fluid (MAF) based on the water-soluble, lipidic and cellular constituents of necrotic tuberculosis lesions. MAF induced drug tolerance in Mtb, remodeled its transcriptome and protected Mtb from dying in microdroplets desiccating in air. Yet survival was not passive: Mtb appeared to rely on hundreds of genes to survive conditions associated with transmission. Essential genes subserving proteostasis offered most protection. A large number of conventionally nonessential genes appeared to contribute as well, including genes encoding proteins that resemble anti-desiccants. The candidate transmission survival genome of Mtb may offer opportunities to reduce transmission of tuberculosis.

**Significance Statement:** *Mycobacterium tuberculosis* (Mtb) travels from the lungs of one person through the air to the lungs of another and survives multiple stresses en route, including changes in temperature and in concentrations of oxygen, carbon dioxide, hydrogen ions, salts and organic solutes. Here we present a genetically tractable model of transmission to begin the identification of the transmission survival genome of Mtb. We devised a fluid that mimics TB lesions, found that it protects Mtb from transmission-related stresses, associated this with the structure of the droplets as they dry and their ability to retain water, and used it to query the potential contribution of each of Mtb’s genes to Mtb’s survival in models of three sequential stages of transmission.

## Introduction

Mtb is the cause of a multi-century bacterial pandemic that only occasionally yields to viral pandemics as the leading cause of human death from infection. While many pathogens of humans complete their life cycle in a reservoir other than the incidental human host, humans are Mtb’s reservoir as well as its victims. In people who are not immunosuppressed, the case fatality rate for tuberculosis (TB) is about 70% for those who have microscopically detectable Mtb in their sputum and receive no treatment^1^. For a pathogen that so readily kills its reservoir, transmission is an evolutionary bottleneck^2^. However, whether any Mtb genes facilitate transmission is unknown. The transmission biology of Mtb^3^ has been studied epidemiologically within populations (e.g.,^4–7^), mechanistically by exposing experimental animals to patients’ exhalations (e.g., ^8^) and inferentially in studies that capture bacilli exhaled by people with TB^9–13^. However, to our knowledge, there has been no pre-clinical model for studying the transmission biology of Mtb under physiologically relevant conditions at a genome-wide level.

The efficacy of TB treatment is monitored by tests on sputum. However, the typical viscosity of sputum (400-700 mPa⋅s)^14^ limits its potential for fragmentation into large numbers of infectious aerosols that are small enough, typically postulated to be < ∼ 5 μm in longest dimension^15–22^, to stay suspended in laminar intrabronchial airflow, escape entrapment on the bronchial ciliary escalator and pass through the narrowest bronchioles to the pulmonary alveoli of a new host. Fluids of lower viscosity are likely to generate more respirable aerosols^20, 23^. While diverse fragmentation processes may generate bacteria-laden respiratory particles^22, 24^, the presence of necrotic pulmonary cavities correlates with infectivity^25^. Cavity contents called “caseum” for their cheese-like viscosity routinely spill into bronchi^26^, where they may be diluted by low-viscosity bronchial secretions. The rheologic properties of the resulting mixture may favor respirable microdroplet formation upon the shearing and bursting fluid fragmentation expected to occur during human tidal breathing, speech, song and violent exhalations, such as coughs and sneezes^24^.

Most preclinical studies of mycobacterial biology have involved laboratory media with markedly different compositions, buffers and osmolarities than human body fluids, often under an atmosphere—room air—that is hyperoxic (O_2_ ∼21%) compared to most human tissues (O_2_ ∼ 5%)_27_ and lacks carbon dioxide. CO_2_ is the equilibrant that maintains the physiologic pH of the human bicarbonate buffer system, a key carbon source for Mtb’s metabolism^28^ and a sensitizer to oxidant injury^29^. Moreover, transmission involves not one set of environmental conditions but sequential passage through a series of environments, in each of which survival of the bacilli may require shared or distinct mechanisms of adaptation.

Here we present the introductory study in a series of efforts of increasing technical difficulty to identify the transmission survival genome of Mtb. Constrained by the equipment available for working at genome scale under biologic safety level 3 conditions, the present paper deals with 2-μL sessile droplets of Mtb. Given that the lessons learned may apply only in part to smaller droplets that are airborne, a forthcoming study based on newly designed devices will involve microdroplets suspended in air. A third study based on other new devices will test individual Mtb genes for their role in infecting mice via aerosol particles of defined size and number.

The present study models three stages of TB transmission in vitro (**Fig. 1A**): (1) residence of Mtb in a hypoxic, necrotic cavity closed off from the airways; (2) erosion of such a lesion into the airways, allowing for a higher level of oxygenation and mixture with bronchial secretions; and (3) expulsion into air in rapidly desiccating microdroplets. Incubation in the model aerosol fluid (MAF) described below mimicked residence in caseum in inducing phenotypic tolerance to several TB drugs^30^. Two novel findings emerged. First, incubation in MAF substantially preserved Mtb’s viability under conditions that model inter-host transit in hyperoxic droplets whose evaporation cools and alkalinizes them as their solutes concentrate or precipitate. Second, Mtb appears to survive under these conditions by calling on large numbers of essential genes and on many genes deemed “nonessential” because their knockout, knockdown or disruption by transposon insertion had not impaired Mtb’s survival in studies under conventional conditions in vitro or in mice.

**Figure 1.**
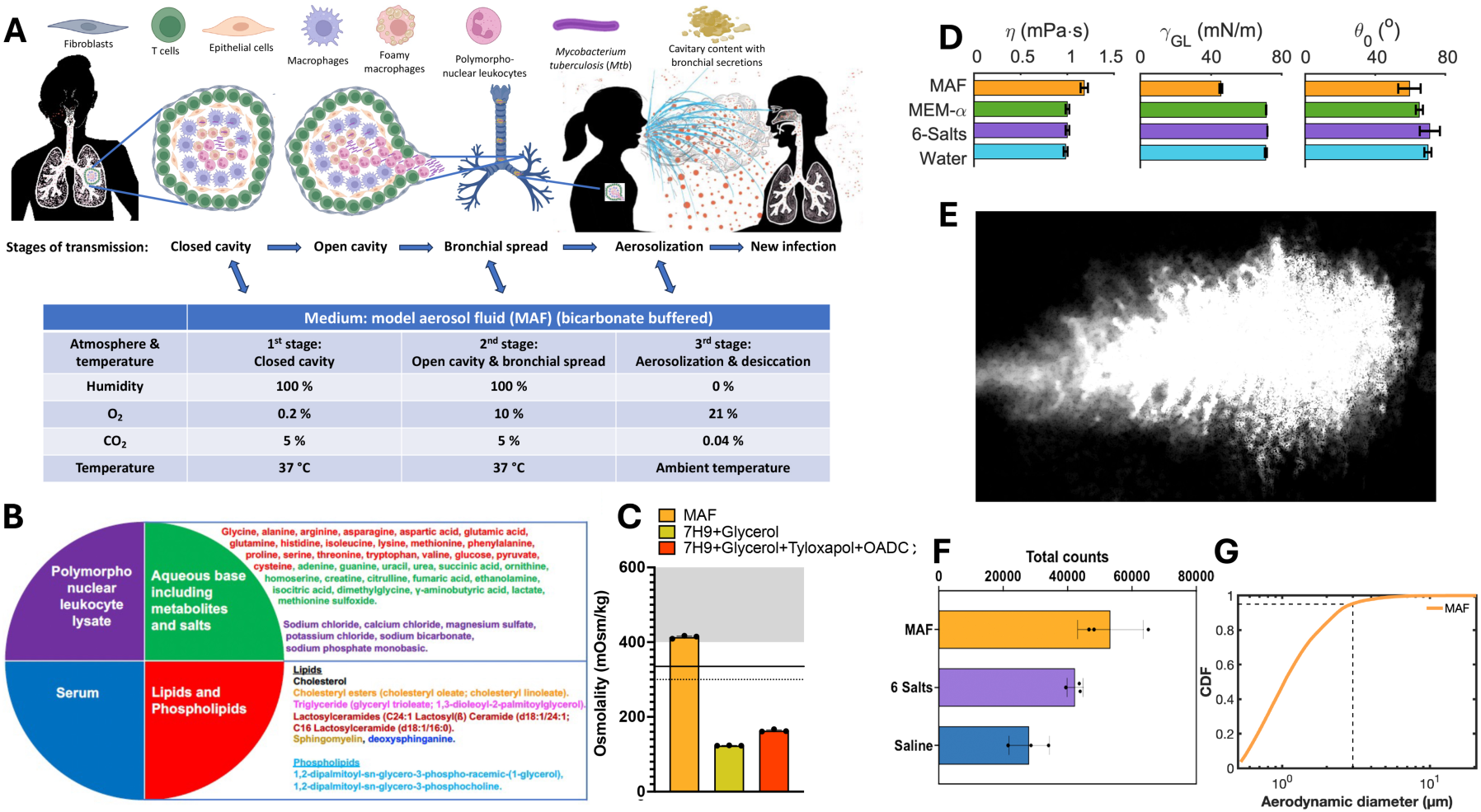
Fluids and atmospheres modeling three stages of transmission. **(A)** Schematic. Part of the cartoon is adapted from Manna and Bourouiba in ^31^**. (B)** Components of MAF. Organic metabolites present both in caseum and in MEM-*α* are in red. Those found in caseum but lacking in MEM-*α* are in green. Inorganic salts present in MEM-*α*are highlighted in purple, along with the organic compound bicarbonate, given its substantial contribution to the osmolality of MEM-*α*. Lipids are colored by class. Concentrations of each component are given in tables S2*A*-S2*F*. **(C) Osmolality** of MAF, 7H9 supplemented with glycerol and the latter further supplemented with tyloxapol, oleic acid, albumin, dextrose and catalase at the standard concentrations listed in tables S2a-S2f. Means ± SEM for 3 measurements. Dotted line indicates the osmolality of MEM-*α*, which mimics human extracellular fluid. Solid line indicates values reported for mouse spleen and shaded area highlights values reported for BCG-infected mouse tissue (see text). **(D)** Dynamic viscosity *η* and surface tension *γ_GL_* of MAF, MEM-*α* and its 6 major salts, and contact angle *θ*_0_ of those fluids on a tissue culture-treated polystyrene surface were measured and compared to MilliQ deionized water. Density is reported in Fig. S1*D*. Means ± SD for at least 3 measurements; individual values not shown where n > 10. **(E)** Cough cloud generated by the exhalation mimic system *ExhaleSimulator*^44^ with a trajectory here shown to span 3 meters from the source. **(F)** Total droplet counts within size range 0.5-20 µm, measured at 1 m from the source in **(E)**. **(G)** Cumulative droplet size distribution of MAF particles counted in **(F)**. Dashed lines indicate that *∼*94% were ≤ 3 µm in diameter.

## Results

### Conditions modeled

There may be diverse mechanisms of generation of infectious Mtb-laden respiratory microdroplets^24, 31^. Here we focused on one mechanism (**Fig. 1A**), beginning with Mtb’s residence in the extracellular milieu of a necrotic lesion in the lung that does not communicate with an airway. In rabbits, such lesions had measured O_2_ concentrations of 1.61 ±   0.37 mm Hg (∼0.2%)^32^. When a necrotic lesion erodes into an airway, its aeration status may vary. Transthoracic needle aspiration revealed that the O_2_ concentration in human TB cavities averaged 17.8% for cavities called “open” (n=7) and 13.6% in cavities called “partially closed” (n = 8)^33^, but concentrations are likely lower in cavities that are too small for transthoracic aspiration and have smaller openings to an airway. We chose to model the open cavity O_2_ concentration at 10%. For both closed and open cavities, we modeled relative humidity of 100% and the physiologic CO_2_ concentration of 5%. To test thousands of individual microdroplets under biosafety level 3 (BSL3) conditions, we were limited to robotic equipment that dispensed 2 μL volumes (diameter ≈1.56 mm) onto polystyrene surfaces, which were maintained in air (O_2_ 21%, CO_2_ 0.04%) at ≈22 °C. To emulate the rapidity of evaporation from smaller, aerosolized microdroplets, we applied a relative humidity of ≲ 5%. As they evaporate, the droplets cool (Fig. S1*A*). As predicted, MAF alkalinizes in air (Fig. S1*B*) as it loses CO_2_ and its source, bicarbonate.

Having designated atmospheres for each of the three stages (Fig. 1A), we turned to the composition of the “model aerosol fluid” (MAF) in which we suspended Mtb (Fig. 1B). Rabbit caseum contained 3-6 μg/mg cholesterol, *∼*2 μg/mg cholesteryl esters and 0.3-9 μg/mg triglycerides^34, 35^. Marmoset caseum contained *∼*10 μg/mg cholesterol, *∼*5 μg/mg cholesteryl esters and 20 μg/mg triglycerides^34^. In human caseum, the predominant triacylglycerides (TAG) were 52:1, 52:2, 52:3, 54:1, 54:2, 54:3 and 54:4^34^. We measured a mean of 70 ng/mL deoxysphinganine in caseum from 4 pulmonary TB lesions in 3 rabbits and 3 TB lesions from 1 marmoset (Fig. S1*C*). We detected and quantified 36 water-soluble metabolites in 8 caseous human TB lesions provided by L. Via and C. Barry (NIH) (table S1*A*, S1*B*). We found 20 of these metabolites present in Minimum Essential Medium-α (MEM-α) with nucleosides (ThermoFisher 41061037) (table S1*B*). To MEM-α we added the remaining 16 metabolites to match their mean levels in human caseum (table S1b). We also added sphingomyelin, C24:1 lactosyl(ß)ceramide (d18:1/24:1) and C16 lactosylceramide (d18:1/16:0) based on semi-quantitative evaluation of published TLC results^36^. The phospholipids 1,2-dipalmitoyl-*sn*-glycero-3-phosphocholine (DPPC) and 1,2-dipalmitoyl-*sn*-glycero-3-phospho-racemic-(1-glycerol) (DPPG) were added to aid emulsification. Finally, we supplemented the aqueous base with 3% (vol/vol) human serum (a level typical of an inflammatory exudate) and sufficient polymorphonuclear leukocyte (PMN) lysate (6 µg DNA/10^6^ PMN) from healthy donors to provide 0.17 μg DNA/μL MAF (S2a), based on the following considerations. Rabbit caseum contained 0.17 μg DNA/mg caseum^35^. Mtb are rarely numerous enough in caseum to be the chief source of the DNA. Though dying macrophages contribute to caseum, PMN (mostly neutrophils, with some eosinophils) are the predominant host cell in sputum and bronchoalveolar lavage (BAL) fluid of TB patients and the cell type most often containing Mtb in cavities, BAL fluid and sputum^37–39^. Necrotic TB lesions are chronic, and PMN have a short half-life. This suggests that products of dead PMN can accumulate. Accordingly, we made the simplifying assumption that most of the DNA comes from PMN and serves as a marker for the other cellular remains of PMN, including their antimicrobial proteins^40^. The final concentrations of each component are listed in tables S2*B-H*, which match or approximate the foregoing measurements, with 2 major differences: we lowered the concentration of proteins by ∼10-fold and of cholesterol, cholesteryl esters, triglycerides and TAG by ∼100-fold to produce an emulsion with a dynamic viscosity in the range of 1 mPa⋅s, for two reasons. This was necessary to allow reproducible robotic dispensation of 2 μL droplets from a suspension in which Mtb was distributed homogeneously, and, as will be seen below, it allowed for efficient generation of aerosols of respirable size. We speculated that these reductions in concentration may mimic dilution of caseum by bronchial secretions. We prepared MAF as a sterile emulsion that remains stable for at least 48 hours. The osmolality of MAF (400 mOsm/kg) was higher than in 7H9 medium (120-160 mOsm/kg), closer to the osmolality of human serum (∼295 mOsm/kg) and between the values reported for mouse spleen (335 mOsm/kg)^41^ and mouse tissue infected with bacille Calmette-Guérin (BCG), the vaccine strain of *M. bovis* (400-600 mOsm)^42^ (Fig. 1C).

### Rheological properties of MAF

We measured rheological properties of MAF and 3 reference solutions (Fig. 1D, Fig. S1*D*, and table S3), the compositions of which are in tables S2*A*, *B*, and *G*). Considering that 93% of the volume of MAF consists of MEM-*α* and 16 additional metabolites, we compared MAF with MEM-*α*, the 6 inorganic salts in MEM-*α* that account for most of its osmolality (“6 salts”) and water. The density, *ρ_L_*, of the fluids is similar (fig. S1d[a]), consistent with their solute mass concentration of < 2%. The dynamic viscosity, *η*, of the inorganic salt solution is similar to that of water (Fig. 1D and Fig. S1*D*[b]). MEM-*α* and MAF have slightly higher dynamic viscosity, consistent with their content of amino acids and proteins (tables S2b,d,f). The surface tension, *γ_GL_*, of MAF was 40% lower than that of the other reference fluids, reflecting its higher content of surface active lipids and phospholipids.^43^

### Physical properties of MAF particles

Next we studied MAF’s propensity to fragment into microdroplets upon aerosolization by an ExhaleSimulator^44^ that generates an analog exhaled respiratory multiphase cloud characteristic of human violent exhalations^45–47^. Fig. 1E shows a representative time-lapse superposed trajectory of the cloud. We compared MAF to a solution of NaCl (“saline”) at the concentration found in MEM-*α* and to the 6 salts and sampled droplets along the trajectories of the cloud at various distances, focusing on particle sizes in the range of 0.5-20 μm. At ≈1 meter from the source, MAF fragmented into a higher number of particles than the other fluids (Fig. 1F), consistent with its lower surface tension (Fig. 1D), and generated a high proportion of droplets of ≤ 3 μm diameter (Fig. 1G), similar to the size distribution of most of the particles containing viable Mtb captured from subjects with TB as they coughed (≤ 0.65 – 3.3 μm)^9^.

### Phenotype of Mtb in MAF

Two distinctive features of Mtb’s residence in caseum are what appears to be a slowly-or non-replicating state and its phenotypic tolerance to antibiotics^30^. We next asked whether Mtb incubated in MAF shares these features. To simulate the closed cavity environment, we incubated Mtb in MAF under 0.2% O_2_, 5% CO_2_ and compared its growth and survival to that of Mtb incubated in MAF under the standard laboratory condition of hyperoxia (21% O_2_) while maintaining CO2 at 5% CO_2_ (Fig. S2*A*). Mtb did not replicate in MAF. Instead, it showed a relatively small time-dependent decline in survival that was greater in hypoxia (**Fig. 2A**) and became relatively tolerant to rifampicin, moxifloxacin, kanamycin and isoniazid. The acquisition of tolerance was gradual in onset and showed a different time-dependence for each antibiotic (Fig. 2B-F), demonstrating that non-replication was not a mechanistic explanation for the tolerance. The same trend was seen for Mtb in MAF under either 0.2% or 21% O_2_. Thus, the medium played a bigger role in inducing tolerance than the oxygen concentration.

**Figure 2.**
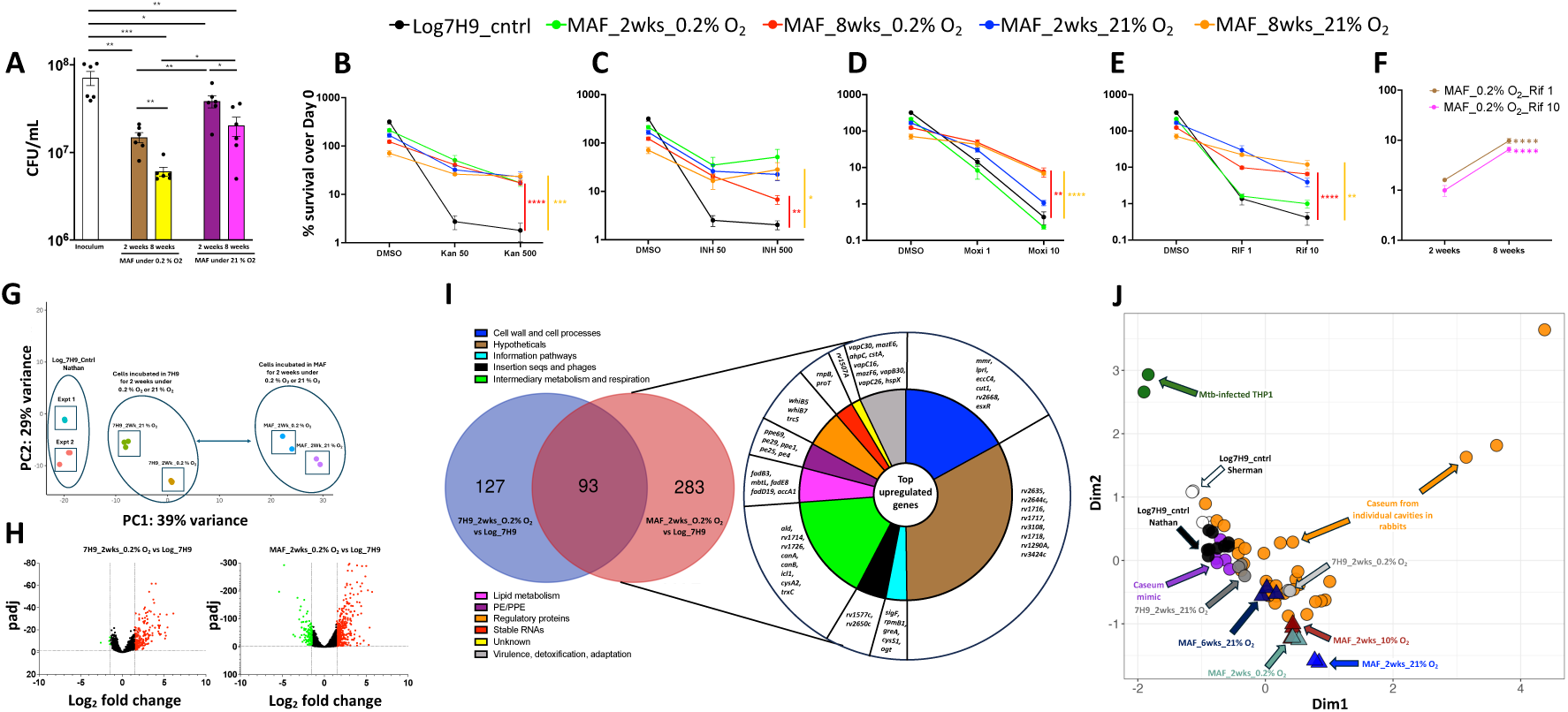
Effects of MAF on Mtb’s survival, drug tolerance and transcriptome in bulk culture. **(A)** Survival of Mtb preincubated in MAF under 0.2 % O_2_, 5 % CO_2_ or 21 % O_2_, 5 % CO_2_ for 2 or 8 weeks. **(B-F)** Survival of Mtb from log phase replicating cultures in 7H9 under 21 % O_2_, 5 % CO_2_ or following preincubation in MAF under the indicated levels of O_2_ and CO_2_ for 2 or 8 weeks and treated with the indicated antibiotics. Kan, kanamycin. INH, isoniazid. Moxi, moxifloxacin. RIF, rifampicin. DMSO, dimethylsulfoxide vehicle (1%). **(A-F)** Data are means ± SEM of two independent experiments. P values were determined by unpaired t test. *P ≤0.05; **P ≤0.01; ***P ≤0.001; ****P ≤0.0001. **(G)** Principal component analysis of gene expression profiles of Mtb from log phase replicating culture in 7H9 under or 21 % O_2_, 5 % CO_2_ or after incubation for 2 weeks in 7H9 or MAF under the indicated O_2_ and CO_2_ concentrations. **(H)** Volcano plot of differentially expressed genes from Mtb incubated under O.2 % O_2_, 5 % CO_2_ in 7H9 or MAF for 2 weeks versus a log phase replicating culture in 7H9 under 21 % O_2_, 5 % CO_2_. **(I)** Comparison of upregulated genes in Mtb incubated under O.2% O_2_, 5% CO_2_ in 7H9 vs MAF for 2 weeks. Selected top genes upregulated in MAF and belonging to different functional categories are listed. **(J)** Multi-dimensional scaling (MDS) of the Mtb H37Rv transcriptome in log phase replicating culture in 7H9 under 21 % O_2_, 5 % CO_2_ or after incubation in 7H9 or MAF under the indicated O_2_ and CO_2_ tensions for 2 or 6 weeks, compared with the following: 34 samples of Mtb-containing caseum from 26 pulmonary lesions in 5 rabbits infected with Mtb HN878 (duplicate samples were analyzed from 8 of the lesions) having CFU burdens ranging from 7.45 x 10^4^ to 1.10 x 10^8^ per gram; Mtb incubated in a caseum mimic prepared from a lysate of human macrophage-like THP1 cells made foamy by incubation with stearic acid; and Mtb-infected THP1 cells. The MDS was kindly generated in the Sherman lab (University of Washington) using the data for Mtb H37Rv in MAF and comparing it to their own results for Mtb HN878 in caseum, caseum mimic and THP1 cells. The data were batch corrected using ComBat-seq^90^ prior to dimensionality reduction.

Transcriptomic comparison of Mtb in MAF vs Mtb in 7H9 under 0.2% O_2_ or 21% O_2_ (Fig. S2*B*) further indicated that MAF does not induce metabolic dormancy but instead drives Mtb on a path of differentiation that differs from that of Mtb in 7H9. This was true regardless of the oxygen concentration, although Mtb did react somewhat differently to hypoxia than to hyperoxia (**Fig. 2G**). As compared to a replicating culture in 7H9, Mtb incubated in MAF for 2 weeks under 0.2% O_2_ upregulated more genes (376) than it downregulated (121), a ratio markedly different than for Mtb in 7H9 reacting to hypoxia (220 genes upregulated; 4 genes downregulated) (**Fig. 2H**). The pattern was similar for Mtb incubated in MAF under 21% O_2_ (Fig. S2*C*). Of the genes upregulated in hypoxic Mtb, 75% of those upregulated during incubation in MAF were different from those upregulated during incubation in 7H9 (Fig. 2I). Among the genes most upregulated in MAF-incubated Mtb during hypoxia were *canA* and *canB,* encoding carbonic anhydrases. These catalyze interconversion of CO_2_ and bicarbonate and may play a role in Mtb’s survival in bicarbonate-buffered MAF under 5% CO_2_. Another highly upregulated gene was *icl1,* encoding isocitrate lyase, whose upregulation has been observed in Mtb in pulmonary cavities^48^ (**Fig. 2I**).

Finally, we compared transcriptomes of Mtb H37Rv in MAF with the transcriptomes of Mtb HN878 in rabbit caseum (GEO accession number GSE273037), in a caseum mimic derived from lipid-fed THP1 human macrophage-like cells^48^ and in THP1 cells themselves. We generated additional RNAseq datasets to simulate progression of Mtb in MAF from the atmosphere of a closed cavity (0.2% O_2_, 5% CO_2_) to that approximating an open cavity (10% O_2_, 5% CO_2_). Mtb in different caseous lung lesions had heterogeneous transcriptomes. Mtb’s transcriptome in each of the test fluids mirrored the transcriptome of Mtb in some caseous lesions but not others (**Fig 2J**).

In sum, MAF is an emulsion that can be prepared at will and in quantity without access to tuberculous caseum and that can induce a physiologic state in Mtb similar to that of Mtb in the caseum derived from some but not all tuberculous lesions. A major difference between MAF and caseum is that the more dilute state of MAF allows for rheological properties (Fig. 1D) that are compatible with simulated human violent exhalation inducing its fragmentation into microdroplets of the size range (Figs. 1E-1G) of Mtb-bearing microdroplets recovered from people^9^.

### Survival of Mtb and BCG in MAF during desiccation

Next, we assessed to what extent Mtb survives during desiccation in MAF compared to survival in water, phosphate buffered saline (PBS), the standard 7H9 medium with glycerol as a supplemental carbon source, and the former with an additional standard supplement consisting of oleic acid, albumin, dextrose, catalase (OADC) and tyloxapol. We also tested survival in MEM-*α*or in a solution of the 6 salts in MEM-*α*that account for most of its osmolarity. We pelleted Mtb from log phase culture in 7H9 with glycerol, OADC and tyloxapol in 21% O_2_, 5% CO_2_, washed the cells and resuspended them in bulk culture in the foregoing media for incubation in 21% O_2_, 5% CO_2_ for 1 day. At that point, recovery of the input number of colony-forming units (CFU) was quantitative. In contrast, when the cells were then allowed to desiccate for 1 day in air in 2 μL droplets, there was a 4.5 log_10_ drop in CFU for Mtb in water, ∼2 log_10_ drop in PBS, ∼1.8 log_10_ drop in the solution of 6 salts, ∼1.0-∼1.6 log_10_ drop in the two 7H9-based media, ∼0.8 log_10_ drop in MEM-*α*, and ∼0.6 log_10_ drop in MAF (Fig. 3A). Among the 6 salts, NaCl contributed most to protection against desiccation, followed by NaHCO_3_, and the two of them together were as effective as all 6 (Fig S3*A*[I]). Addition of the water-soluble metabolites of MAF that are not present in MEM-*α* did not increase protection. However, the lipids, phospholipids and proteins of MAF augmented protection during periods of desiccation longer than 1 day (Fig. S3*A*[II]). Results were similar when Mtb was passed through the 1^st^ and 2^nd^ stages of the in vitro transmission model before desiccation (Fig. *S3B*). BCG likewise survived for 1 day in bulk culture in water, 6 salts, MEM-*α* and MAF. However, upon desiccation in microdroplets, BCG lost all detectable CFU in water and survived only marginally better when desiccating in the 6 salts solution. MEM-*α* and MAF afforded the best protection of desiccating BCG (Fig. 3A).

**Figure 3.**
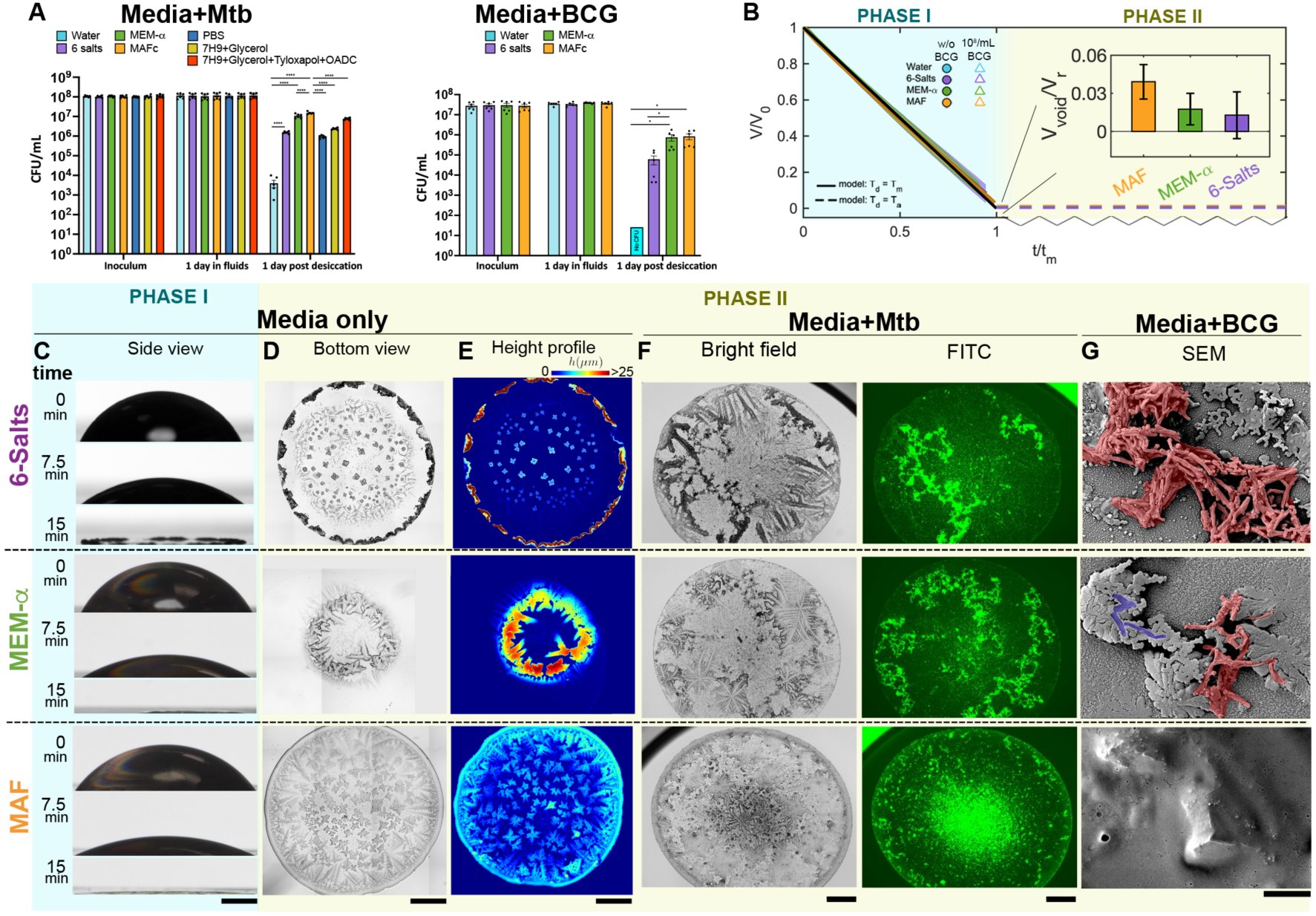
Properties of desiccating droplets. **(A)** Survival of Mtb and BCG taken from log phase culture in 7H9 and transferred to the indicated fluids for one day in bulk culture before desiccation for one day in 2 μL droplets. Means ± SEM of two independent experiments, each in triplicate. *P ≤0.05, ****P ≤0.0001 by unpaired t test. **(B)** Normalized volume, V/V_0_, as it changes from V_0_ = 2 μL in sessile droplets of MAF, MEM-*α*, 6 salts, deionized water, and MAF, with and without BCG, over normalized time, t/t_CCRL_, compared to theoretical predictions taking the droplet temperature, T_d_, to be either ambient temperature, T_a_, (dashed black line) or measured droplet surface temperature, T_m_ (solid black line); see Supplementary material and Fig. S3*C,D*. Phase I lasted about 15 minutes at relative humidity ≤ 5% and ambient temperature Ta =22 °C. Inset shows void fraction, *V_void_*(/*V_r_* = (*V_r_* – *V_S_*)/*V_r_*, where *V_r_* is the terminal residue volume and *V_S_* is the compact volume of dry solutes. Shown for 10 measurements per condition (mean ± SE). **(C)** Time-lapse photographs capturing side views of evaporating droplets in Phase I and the transition to Phase II, where the residues of the evaporated droplets formed. The edge of the droplets remained pinned except for MEM-*α*, which receded to the partially pinned edge towards one side. Scale bar = 500 μm. **(D)** Bottom views of the residue of droplets in Phase II. Scale bar = 500 μm. **(E)** Height profile of droplets in Phase II, also quantified in Fig. S3*E*, serves as basis, with Fig. S3*F*, to compute the terminal residue volume, *V_r_*, used in Phase II inset in **(B)** for each fluid. Scale bar = 500 μm. **(F)** Brightfield and fluorescence photomicrographs of 2-µL desiccated droplets of the indicated fluids containing Mtb-Mrx1roGFP2. Scale bars = 500 μm. **(G)** Scanning electron micrographs of the surfaces of desiccated droplets containing BCG. Crumpled BCG cells (false-colored red) were found on the surface of desiccated droplets of 6 salts and MEM-*α*. Some non-deformed BCG (false-colored blue) were also found on the surface of desiccated droplets of MEM-*α*. No BCG were found on the surface of desiccated droplets of MAF. Scale bar = 5 μm.

In sum, MAF has 3 important properties relevant to transmissibility of Mtb: induction of relative antibiotic tolerance, induction of relative desiccation tolerance, and efficient generation of aerosols of respirable size. Induction of antibiotic tolerance would not have begun to advantage Mtb’s transmission until about 75 years ago, and we have not compared simpler fluids for induction of antibiotic tolerance. However, MAF outperformed the simpler fluids we tested with respect to desiccation tolerance and aerosol generation.

### Physical environment of the desiccated droplets

To explore how MAF may protect Mtb from desiccation, we first compared the rate of evaporation of 2 μL droplets of MAF and the comparator fluids on the same tissue culture-compatible polystyrene surfaces used to study survival of desiccating Mtb, given that different timescales of evaporation could lead to distinct stresses on the organisms. Contact angles varied from 50° to 80°, with MAF being the most wetting, consistent with its surface-active components, but there was no significant difference in initial evaporation dynamics (Phase I) (Fig. 3B). From side-view imaging (Fig. 3C) we measured droplet volume and its rate of change, with and without BCG, which was compatible with a first-order diffusion-driven evaporation law^49^ when corrected to account for surface wetting, expected evaporative cooling, and surface heat-exchange (Fig. 3B, Figs. S3*C,D* and Supplemental Materials). However, the Phase I evaporation is macroscopic (milimeters) compared to the size of mycobacteria (microns). To characterize evaporation in the micro-environment of most relevance to the mycobacteria (Phase II), we turned next to smaller-scale imaging. Figs. 3C-F reveal fluid-specific structural distinctions among the residues of the desiccated sessile droplets of MAF, MEM-*α* and 6 salts. Quantitative profilometry showed distinct height profiles at the periphery (Fig. 3E and Fig. S3*E,F*) typical of “coffee-ring” elevated edges (Fig. 3D, E and Figs. S3*F-I*)^50^ that emerge via capillary driven flow, favoring solute transport to and deposition along the droplet’s boundary contact line edge. Residues from desiccated water droplets had little visible internal structure (Fig. S3*H, I*) as expected, while cuboidal and dendriform crystals emerged in residues of desiccating 6 salts and MEM-*α* droplets, respectively. In these fluids Mtb and BCG appeared to coat or be colocalized with some of the larger dendritic crystals (Fig 3F). In contrast, MAF residues were characterized by more amorphous crystals (Fig 3F). Moreover, in residues of MAF (Fig. 3F), BCG (Fig. S3*H*) and Mtb (Fig. S3*I*) clustered in the center. Both differences are consistent with the role that surfactants and proteins can play in altering crystal and residue formations^51–53^. Analysis of atomic composition confirmed that the crystals were mainly made of NaCl, while O was distributed throughout the residues of MAF and appeared enriched along the crystal borders for the other two fluids (fig. S3*J*).

We integrated the height profile of the desiccating droplets to measure the volume of the residues *V*_#_ in Phase II. We calculated the volume of solutes *V*_$_ using their mass and density and the associated residue “void” fraction as *V*_%&’(_/*V*_#_ = (*V*_#_ − *V*_$_)/*V*_#_. Residues of MAF held the highest “void” fraction, compatible with higher hydration (Fig. 3B).

Scanning electron microscopy (SEM) revealed that in droplet residues of water, 6 salts and MEM-*α*, BCG cells were found on the external surface (Fig. 3G and Figs. S3*K, L*). In contrast, bacteria were not seen on the surface of MAF residues (Fig. 3G and Fig. S3*M*). Instead, fluorescence microscopy revealed BCG (Fig. S3*H*) and Mtb (Fig. 3F and Fig. S3*I*) confined to the interior of MAF residues. BCG exposed on the surface of residues of MEM-*α*, 6 salts (Fig. 3G and Figs. S3*K, L*) and water (Fig. S3*K*) appeared shrunken and crumpled (artificially colored red in Fig. 3G), reminiscent of mechanical buckling, wrinkling, and dimple formation of emptied shells^54^ and consistent with the shrinking and wrinkling of bacterial membranes in hypertonic solutions^55, 56^. In MEM-*α*residues, some BCG cells encased in crystals (artificially colored purple) appeared to retain their outer structural integrity (Fig. 3G). In sum, physical properties of the media confer important heterogeneity and shape distinct local structures upon desiccation. In contrast to the other fluids, desiccating MAF forms a matrix that may protectively encapsulate mycobacteria while also maintaining their shape and apparent integrity.

### Candidate transmission survival genome of Mtb

To conduct genome-wide screens, we used a CRISPRi library that targets 4014 Mtb genes with an average of 20 sgRNAs for each gene^57^. We pre-depleted the library with anhydrotetracycline (Atc) for 5, 10 and 20 generations in 7H9 in 21% O_2_, 5% CO_2_ to achieve varying degrees of gene silencing. Each generational version of the library was then resuspended in MAF and sequentially subjected to one or more of the 3 stages of the in vitro transmission model: bulk culture for 1-2 weeks in a closed cavity environment (0.2% O_2_, 5% CO_2_), followed by bulk culture for 1-2 weeks in an open cavity environment (10% O_2_, 5% CO_2_), followed by desiccation in 2 µL droplets for 18-24 hours or 3 days in an aerosol environment (21% O_2_, 0.04% CO_2_). It is possible that Atc-dependent suppression of gene function may not have persisted through the longer periods of culture; if so, this might have led to some false-negative results. **Fig. 4A and Fig. S4*A-C*** list the 3 independent CRISPRi screens done for different stages. Libraries collected after each stage of the model with and without outgrowth in complete 7H9 medium in 21% O_2_, 5% CO_2_ were subjected to genomic DNA isolation and sgRNA sequencing. Post-sequencing analysis using MAGeCK^58^ identified significant depletions by comparing sgRNA abundances between conditions. We identified as a “significant difference” any decrease in sgRNA read-counts at a level corresponding to a log_2_ fold-change (FC) of ≤ -1.5 with a padj ≤ 0.05 (where “padj” is a p-value corrected for multiple hypothesis testing) when comparing a condition to any earlier condition. Thus, the number of possible comparisons increased with each successive stage: 12 for stage 1, 16 for stage 2 and 20 for stage 3 (Fig 4A) (during manual quality control, two sample sets belonging to stage 3 were discarded due to missing values or zero LFC values resulting in 18 comparisons for stage 3) as detailed in https://doi.org/10.7298/e4xs-y66359. We then tallied the number of significant differences (NOSDs) for each of the 4014 Mtb genes both for individual stages and for all 3 stages combined (**Fig. S4D)**. The larger the NOSDs for a given gene, the stronger the inference that the gene contributes to Mtb’s survival in at least one stage of sequential testing and likely in more than one stage. Smaller NOSDs for a given gene are more difficult to interpret. A NOSDs of 1 indicates non-reproducibility in the independently conducted screens, but could be of interest for hypothesis generation. A NOSDs of 2 or 3 at an early stage of testing could be meaningful if knockdown of the gene reduced the population of cells bearing those sgRNAs to such an extent that there were insufficient reads to allow statistical inference at later stages.

**Figure 4.**
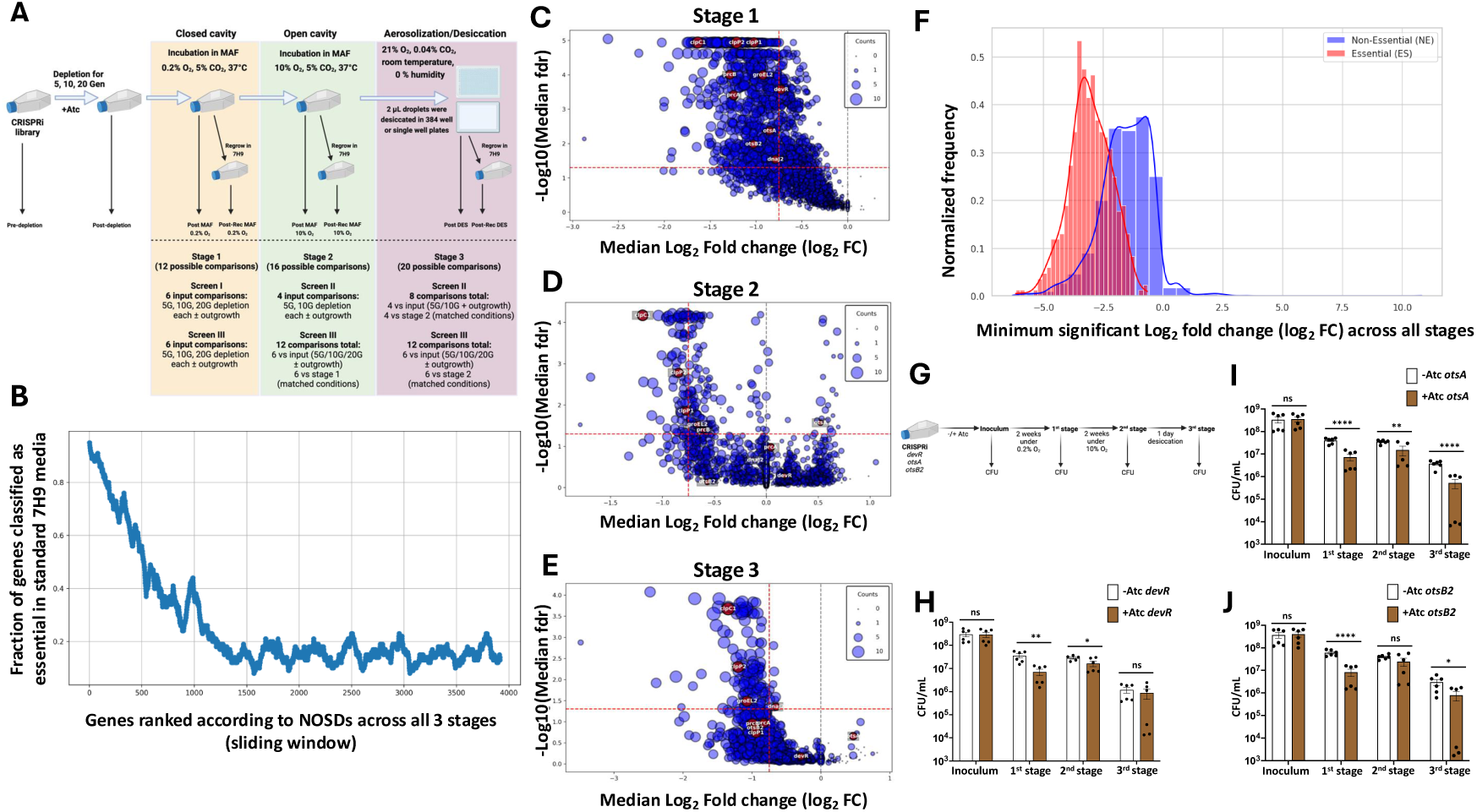
Genome-wide CRISPRi screen of Mtb during three modeled stages of transmission. **(A)** Schematic of the screen across transmission stages (created with BioRender.com) and summary of comparisons. Each stage allows for multiple possible comparisons where significant differences may be detected. Stage 1: 12 possible comparisons against the input library across two screens, each testing three pre-depletion times (5, 10, 20 generations) with and without outgrowth in 7H9 in 21% O2, 5% CO_2_ after exposure to the conditions of the stage. Stage 2: 16 possible comparisons, combining comparisons to the input library and stage-to-stage comparisons. Stage 3: 20 possible comparisons. ’Matched’ comparisons maintain the same pre-depletion time and outgrowth condition between stages (e.g., comparing Stage 2 vs Stage 1 at 5 generations, no outgrowth). **(B)** Variation in frequency of conventionally essential genes in MAF as a function of the NOSDs for each gene across all stages of the CRISPRi screen. Genes with the highest NOSDs are listed at the left and those with fewest NOSDs or none to the right. Within each set of 100 consecutive genes, the proportion of genes is indicated that were previously categorized as essential in standard culture conditions in 7H9 or 7H10 media. **(C-E)** Volcano plots for 1^st^, 2^nd^ and 3^rd^ stage, in which the median log_2_ FC for all significant differences in expression of a given gene at any stage compared to the starting libraries or the earlier stages is plotted against the median false discovery rate for those differences. Symbols for each gene are sized in proportion to the NOSDs (“Counts”). **(F)** Frequency distribution of the most negative log_2_ FC for each gene at any stage according to whether the gene has been characterized as essential (red bars) or nonessential (blue bars) under conventional conditions. For ease of visualization, the height of each histogram is normalized to equalize the areas under the curve for genes of the two classes. **(G)** Schematic of CFU survival assay described in Fig. 4H**-J** (created with BioRender.com). **(H-J)** Survival assay for CRISPRi knockdown strains of **(H)** *devR*, **(I)** *otsA* and (**J)** *otsB2*. Data are means ± SEM of two independent experiments. P values were determined by unpaired t test. *P ≤0.05; **P ≤0.01; ****P ≤0.0001; ns = non-significant.

The highest NOSDs achieved for an individual gene were 10 for the 1^st^ stage, 8 for the 2^nd^ stage, 13 for the 3^rd^ stage, and 28 for the three stages combined, as illustrated in **Fig. S4D** . The highest NOSDs in any stage were registered for genes of both known (*clpC1, ftsZ, rpoC, tatC, mtrA, secY, rnc, dnaB, rfe, clpP2, pknB, gyrB, murB*) and unknown (*rv1828*, *rv0430, rv1697, rv1480, rv2969c*) function. The numbers of genes with NOSDs ≥ 1 were 2508 in the 1^st^ stage, 1596 in the 2^nd^ stage, and 1949 in the 3^rd^ stage. Collectively, 69% of the 4014 genes had NOSD ≥ 1 and 45% had NOSD ≥ 3 in any of the stages (31%, 10% and 29% in 1^st^ to 3^rd^ stages, respectively). In sum, these results suggest that a significant proportion of Mtb’s genome may contribute to its survival in one or more of the modeled stages of transmission.

In Fig. 4B, we ranked genes in sliding windows of 100 from those with the highest to the lowest NOSDs. Within each window we noted the proportion that have been designated as conventionally essential or conventionally nonessential in 7H9 in 21% O_2_, 5% CO_260_ (**Fig. 4B**). Most genes with high NOSDs are conventionally essential. However, 64% (1873) of conventionally non-essential genes had NOSDs ≥ 1, along with 457 that are conventionally essential.

Next, we focused on stage-specific comparisons to identify some of the critical genetic pathways predicted to sustain Mtb during the stresses of transmission, aided by volcano plots of the median log_2_ FC for all stage-relevant comparisons against the negative log_10_ of the median FDR-adjusted p-value (Fig. 4C-E). Among the topmost hits were *clpC1*, *clpX, clpP1* and *clpP2*, which encode the Clp chaperonin-protease complex (Fig. 4C-E). Also prominent were other proteostasis pathway genes: proteasome genes *prcA* and *prcB*, chaperonin *groEL2* and chaperone *dnaJ2*.

The conservative presentation of *median* FC across multiple comparisons for individual stages obscures the extent of depletion (negative FC) imposed by any one of the sequential stresses in any one comparison. In contrast, Fig. 4F shows each gene’s maximally negative, statistically significant FC in any one condition for any one comparison according to whether the genes are designated as conventionally essential or nonessential. In general, there were more severe consequences for depletion of conventionally essential genes than for conventionally non-essential genes. Nonetheless, there were substantial survival penalties under the conditions tested for the individual knock-down of large numbers of conventionally non-essential genes.

To test the validity of the screens, we cross-referenced known regulators of Mtb’s response to hypoxia and desiccation. The regulator of hypoxia response *devR* was among the top hits in the 1^st^ stage (0.2% O_2_). During the 2^nd^ stage (10% O_2_), the median log_2_ FC for *devR* increased from -0.73 to +0.18 (Fig. 4C-D and respective source data files); that is, its knockdown ceased to appear consequential at the higher level of O_2_. During hypoxia, trehalose monomycolate and trehalose dimycolate are catabolized into activated pentose phosphate intermediates, which may prepare Mtb to resume peptidoglycan biosynthesis upon reaeration_61_. Trehalose biosynthesis genes *otsA* and *otsB2* were among the top hits during hypoxia (Fig. 4C). Their NOSDs decreased and median log_2_ FC values increased (that is, became less negative) upon the shift from 0.2% O_2_ to 10% O_2_ (Fig. S4*E* and source data files for Fig. 4C-D). Proteostasis pathway genes and many others followed the same trend (Fig. S4*E*), suggesting that transfer from an O_2_ concentration of 0.2% to adaptation in 10% O_2_ might allow the subsequent outgrowth in room air of cKD cells that would fail to replicate if taken directly from 0.2% O_2_ to 21% O_2_ in a medium that supports replication. We then created separate CRISPRi knockdown strains for *devR*, *otsA* and *otsB2* (table S4), pre-depleted the genes in the presence of Atc, transferred the cells to MAF and carried them sequentially through the 3 stages (Fig. 4G). Depletion of *devR*, *otsA* and *otsB2* in the closed cavity conditions each led to a *∼*0.5 log_10_ drop in CFU. For the *devR* knockdown strain, the survival defect was not increased upon shift to the open cavity environment (Fig. 4H). Individual knockdown of *otsA* and *otsB2* caused *∼*0.5 log_10_ decrease in CFU upon desiccation (Fig. 4I and 4J), perhaps reflecting the role of trehalose as an osmoprotectant.

### Dependence of transmission-related survival on proteostasis genes

Genes encoding *clpC1* and other members of the Clp complex were the most highly depleted genes during all stages, particularly during desiccation, along with other members of proteostasis pathway, namely *prcBA*, *dnaJ2* and *groEL2*. (Fig. 5A). Conditional knockdown strains for *clpP1P2*, *groEL2* and *dnaJ2* were pre-cultured in the absence or presence of Atc and tested along with a Δ*prcBA* knockout strain and the corresponding wild type Mtb. We sequentially incubated the strains in MAF in the 1^st^ stage for 2 weeks, the 2^nd^ stage for 2 weeks and the 3^rd^ stage (desiccation) for 1 day. Compared to the controls, knockdown of *clpP1P2* and knockout of *prcBA* led to ∼1 log_10_ drop in CFU during the 1^st^ stage (Fig. 5B, C) but no drop in the 2^nd^ stage. In fact, the CFU level rose in the 2^nd^ stage (Fig. 5B-C). Considering that Mtb does not replicate under those conditions, the reappearance of CFU suggests that *clpP1P2* knockdown and Δ*prcBA* Mtb may have formed “differentially detectable” (DD) Mtb^62^ in the 1^st^ stage. DD Mtb have been shown to form in response to oxidative damage^62^. CFU dropped again with desiccation of the *clpP1P2* knockdown and Δ*prcBA* strains in the 3^rd^ stage (Fig. 5B,C). Knockdown of the chaperonin *groEL2* and the chaperone *dnaJ2* led to a drop in survival of more than ∼0.5 log_10_ only during the 3^rd^ stage (Fig. 5D,E). We saw a similar phenotype with hypomorphs of *clpC1* and *clpP1P2* produced by CRISPRi, where the phenotype reversed in the 2^nd^ stage and reappeared in the 3^rd^ stage (FigS5*A*).

**Figure 5.**
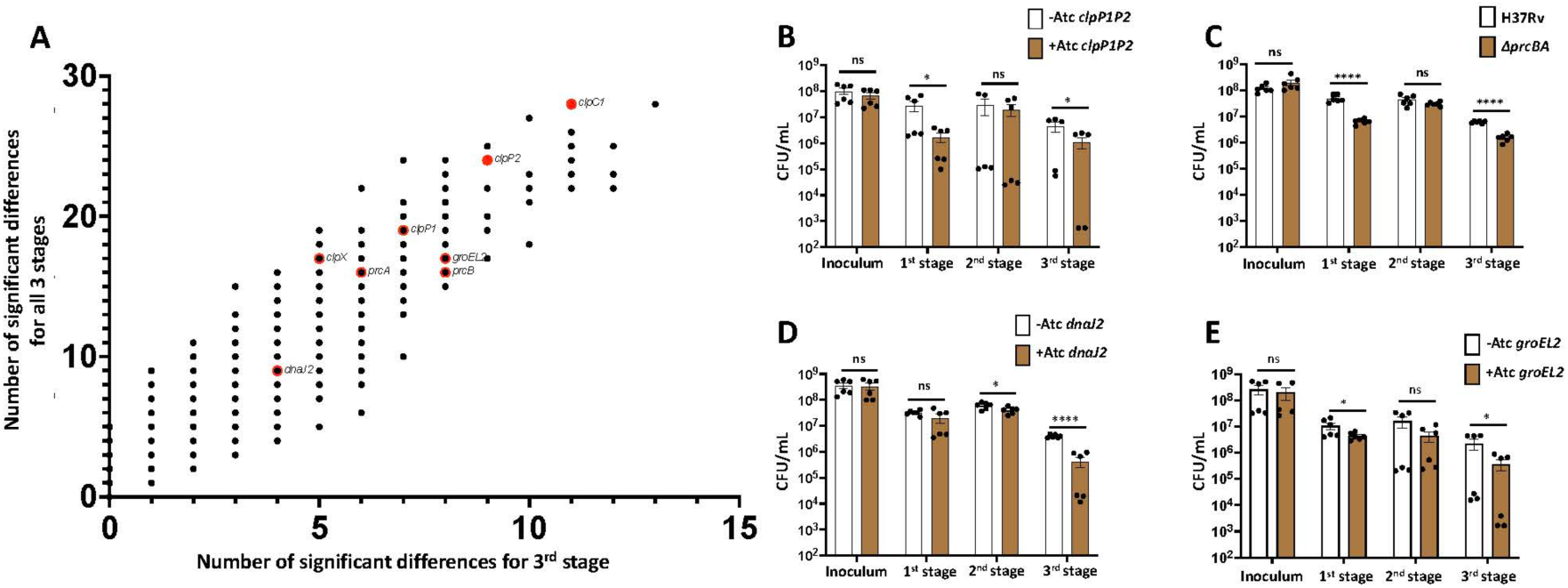
Prominence of proteostasis genes in Mtb’s survival during modeled transmission. **(A)** Comparison of NOSDs for all stages vs 3^rd^ stage. Each dot corresponds to a gene with NOSD ≥ 1 in the 3^rd^ stage. **(B-E)** Survival assay for Mtb knockdown or knockout strains of **(B)** *clpP1P2,* **(C)** *prcBA,* **(D)** *dnaJ2* and **(E)** *groEL2*. Data are means ± SEM of two independent experiments. P values were determined by unpaired t test. *P ≤0.05; ****P ≤0.0001; ns = non-significant.

There were 121 genes with NOSDs ≥ 1 *only* during the 3^rd^ stage (desiccation) (Fig. S5*B*), including *rv3662c* (NOSDs = 6), *rv0443* and *fadE5* (NOSDs = 5), *rv1404* (NOSDs = 4) and *rv1435c, rv3195, rv2661c, rv3716c* and *narG* (NOSDs = 3). Six of these 9 genes are bioinformatically linked to the *clp* system, and two of them, the conserved hypothetical proteins *rv3195* and *rv3716c*, are linked with each other as well (Fig. S5*C*) (https://string-db.org/network/83332.Rv2460c)^63^. The levels of mRNA of their homologs in *Corynebacterium glutamicum* (NCgl0748 and NCgl0240) are increased in that species’ *ΔclpC* mutant and restored upon deletion of *clgR*, a transcriptional regulator whose homolog in Mtb activates genes encoding ClpP1, ClpP2, two other proteases (PtrB and Rv1043c), ClpC, chaperones ClpB and Acr2 and chaperonin Acr2^64^. Moreover, Rv0443 was enriched in *clpC1*-and *clpP2*-depleted Mtb^65^, while Rv1404 and Rv2661c are predicted to interact with ClpC2 (http://galaganlab.bu.edu/tbdb_sysbio) and NarG with ClpP2 (https://www.hitpredict.org). In *M. smegmatis*, D’Andrea identified interactions of ClpC1 with the homologs of FadE5, Rv3195 and Rv3716c^66^.

### Candidacy of hydrophilin-like PE-PGRS proteins

In 2000, Garay-Arroyo et al. reported that the late embryogenesis abundant (LEA) proteins of plants increased in abundance in seeds at the onset of desiccation^67^. In their hydrophilicity and abundance of glycine, the LEA proteins resembled proteins the authors called “hydrophilins” in *Saccharomyces cerevisiae* and *Escherichia coli* that increased during hyperosmosis^67^. The hydrophilin proteins DtpA and DtpB in *Acinetobacter baumannii* are intrinsically disordered^68^. We used hydrophilicity index, glycine content, and the fraction of residues classified as belonging to intrinsically disordered regions to identify a cluster of 50 conventionally nonessential Mtb genes with high values for all 3 descriptors (Fig. 6A; source file for Fig. 6A; Fig. S6). Of these, 49 belong to the PE-PGRS family. Those with a higher NOSDs seem more likely to play a role in tolerance to desiccation, but even those with a NOSDs of 1 may warrant investigation, given the striking physical properties predicted for the encoded proteins and the possibility of redundancy among the 27 members of the family with NOSD ≥ 1 in the desiccation stage (Fig. 6B). The only protein in the cluster not belonging to the PE-PGRS family is Rv0378, a 73-amino acid protein predicted to be highly disordered (Fig. S6).

**Figure 6.**
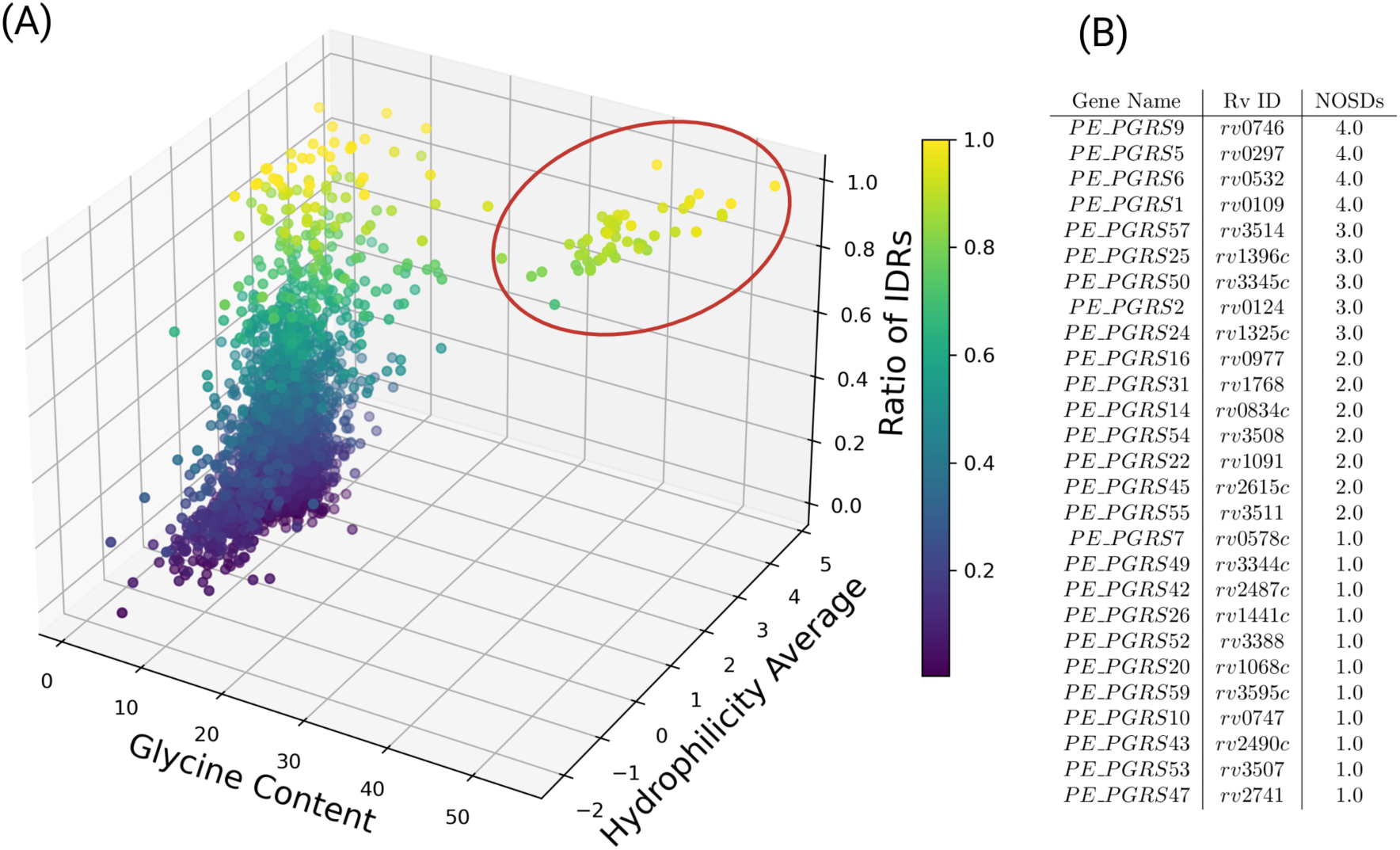
Identification of hydrophilin-like proteins as desiccation stage candidates in the transmission survival genome of Mtb. **(A)** Analysis of the full Mtb proteome in three descriptor dimensions: glycine content, hydrophilicity (averaged across all residues in each protein), and ratio of residues predicted to be in intrinsically disordered regions. Highlighted cluster contains 50 proteins. **(B)** Proteins in the highlighted cluster whose genes had NOSD ≥ 1 in the desiccation stage.

## Discussion

In 1998, Cole et al.^69^ reported the sequence of Mtb’s genome. The ensuing quarter century has witnessed a near-exponential increase in publications about Mtb genes along with multiple breakthroughs in relevant technologies. Yet the proportion of Mtb genes with defined physiologic functions has approached an asymptote, with about 27% of genes unassigned^70^ and a similar proportion bearing speculative annotations. Here we have addressed two possible reasons for this knowledge gap: use of unphysiologic culture media and lack of models of aerosol transmission amenable to the study of genotype-phenotype correlation. Many of the genes of unknown function are also conventionally non-essential, and many conventionally non-essential genes are members of the candidate transmission survival genome. That may help to reveal the function of some of them, as speculatively suggested by the findings of hydrophilin-like features of multiple PE-PGRS proteins in the candidate transmission survival genome (Fig 6).

In 1947, Dubos and Middlebrook introduced a medium that supported Mtb’s replication as fast as or faster than any other, noting that “Mixtures of sodium and potassium phosphates provide a satisfactory buffer system”^71^. Such media, including today’s Middlebrook 7H9, 7H10 and 7H11, bear little resemblance to the human environments in which Mtb spends all of its natural life except in transit to new hosts. Standard culture conditions for Mtb are hypotonic, hyperphosphatemic, alipidic, hypocapnic and hyperoxic relative to Mtb’s human environments. We compiled published and original observations on the composition of necrotic cavities from Mtb-infected people, non-human primates and rabbits to develop MAF. Rather than strive for the simplest mimic of cavity fluid that induces some pre-determined state in Mtb, we sought to include every known component of cavity fluid, reasoning that any component might influence the fluid dynamics^24^ and transmission biology in unanticipated ways. For example, to our knowledge, this is the first report of dihydrosphinganines in tuberculous caseum. Their impact is unknown, but sphingomyelin promotes Mtb’s growth in vitro^72^, certain dihydrosphingosines have potent antimycobacterial activity^73^, and sphingolipids can affect the fusion of lysosomes with mycobacteria-containing phagosomes^74^.

The drug tolerance and transcriptomic phenotypes of Mtb incubated in MAF mimicked those of Mtb in authentic caseum^30^ and in a caseum mimic made from the aged lysate of stearic acid-fed, macrophage-like THP1 leukemia cells^48^. Compared to water, 7H9, PBS, and the mammalian cell culture fluid MEM-*α*, MAF was most effective at protecting Mtb from death during desiccation in air. The reasons require further study, but may include three features of the fluid biophysics of desiccating droplets of MAF: (1) they appeared to retain more water than comparator fluids; (2) they formed residues that excluded mycobacteria from the air-exposed outer surface; and (3) they promoted aggregation of Mtb in the center of the droplet, reducing the proportion of mycobacterial surfaces exposed to the suspending medium. Such aggregation might favor retention of intrabacterial water, much as aggregation of *Streptococcus pneumoniae* protects them from desiccation^75^. It will be of interest to explore whether biomolecular condensation of proteins within Mtb in desiccating droplets liberates water from the proteins’ hydration shells to buffer the stress of hyperosmolarity and cooling during desiccation^76^.

Using MAF and a sequence of atmospheres, we sought to build a list of genes that Mtb may require to survive emergence from a hypoxic state in a necrotic, PMN-rich lesion into a euoxic pulmonary cavity and expulsion into the air in a micro-droplet of bicarbonate-buffered fluid small enough to stay suspended until it is inhaled, while cooling, concentrating, alkalinizing and becoming hyperoxic. These stresses constitute an evolutionary bottleneck for Mtb. The “transmission survival genome” sculpted by that bottleneck may encode potential targets for intervention to reduce transmission. Interest in finding ways to interrupt transmission^77^ is heightened by the finding that 80% of people with pulmonary symptoms reporting to a TB clinic in a high-incidence community who did not receive a diagnosis of TB were nonetheless found to be exhaling viable Mtb^78^. Given TB’s reproduction number (R_0_) of 5-15 (second only to measles’), preventing transmission is indispensable to end the TB pandemic.

The need to cope with oxidative stress and to refold damaged proteins or dispose of them emerged as a prominent role of the genes on which Mtb depended to survive modeled transmission. Most bacteria die upon desiccation^79^. Some, like *B. anthracis,* survive by forming spores. Bacteria that forego sporulation yet repair their DNA upon rehydration and survive, such as species of *Deinococcus*, *Chelatococcus*, *Methylobacterium* and *Bosea*, are those that succeed in protecting their proteins from oxidation^80^. Similarly, genes contributing to *Acinetobacter’s* desiccation tolerance encoded antioxidant, DNA repair and proteostatic functions, as well as hydrophilins.^68, 81^ *Acinetobacter’s* hydrophilins are speculated to act as chaperones protecting proteins from dehydration-induced denaturation^81^. We identified 50 conventionally nonessential Mtb genes that resemble hydrophilins in having high proportions of hydrophilic amino acids and glycines^67^ and being intrinsically disordered. Our screens indicated that 27 of these may warrant investigation as potential contributors to Mtb’s survival from desiccation. If further work confirms that role, it would add to the diverse functions ascribed to the PE-PGRS gene family^82^ related to their contribution to virulence^83^.

Proteins are expressed at levels close to their solubility limits in *C. elegans*, risking aggregation with loss of cell water^84^. When falling temperature increases cytosolic viscosity, yeast cells adapt by regulating their synthesis of glycogen and trehalose^85^. This may help explain why genes involved in trehalose metabolism were prominent in our screen.

There have been few pre-clinical models of aerosolization of Mtb and none to our knowledge that identified genes that contribute to Mtb’s survival. Loudon et al.^86^ observed ∼ 1 log_10_ reduction in Mtb CFU within 60 minutes after aerosolization in “Dubos medium”. Lever et al.^87^ observed a ∼2 log_10_ reduction in Mtb CFU within 60 minutes when bacteria taken directly from aerobic culture on plates containing Middlebrook 7H10 medium were mechanically aerosolized in artificial saliva containing a bovine salivary gland glycoprotein^88^. Pfrommer et al^39^ studied Mtb aerosols generated in PBS. To our knowledge, no earlier in vitro studies of mycobacterial aerosols have pre-conditioned the bacteria in lesion-like conditions. Collection of exhaled Mtb from patients with TB is an important research tool^9–13^ but does not allow for deliberate, selective or systematic genetic variation in the Mtb.

In sum, studies of MAF as a model for TB lesional fluid are consistent with evidence that Mtb induces the host to provide it with a medium that confers tolerance to some antibiotics^30^ and reveal that such a fluid can promote formation of aerosols in a size range consistent with substantial time in the air and access to pulmonary alveoli in new hosts, while partially protecting Mtb from death in rapidly shifting environmental conditions before and during its airborne journey. To the extent that our study of sessile droplets is relevant to aerosol transit, Mtb may depend not only on the host fluid in which it is carried but also on hundreds of its own genes. Study of Mtb’s transmission survival genome may also reveal essential functions for many of the 3008 genes that have been called non-essential^60^ based on experiments in laboratory conditions and animal models that do not include transmission.

### Limitations of the study

Our model of sequential stages of transmission of Mtb necessarily has many limitations. No one set of conditions can recapitulate the heterogeneity of TB lesions, and not all transmission of TB originates from cavities. MAF is a complex medium and inclusion of human serum and PMN lysate means that MAF is not fully defined. We did not model the oscillations in the pH of airway fluids that occur with tidal breathing. Such oscillations were bactericidal for *Pseudomonas aeruginosa*^89^. Our study was limited to the dispensation of relatively large, sessile droplets on an artificial surface. That slows the desiccation process by orders of magnitude compared to what occurs in smaller in-flight microdroplets surrounded by air. The resulting differences in rate of change, phase transition, and composition and structure of the intra-droplet environment could affect which genes Mtb depends on to survive and to what extent. This makes it risky to extrapolate from the relatively modest survival deficits we observed here for individual knock-out and knockdown strains to what may be seen during aerosol infection of mice. The survival deficits for individual gene-depleted strains should be assessed in the context that each of the separately tested genes appears to act nonredundantly and they and others would act collectively. The operational scale of genome-wide screening obliged us to choose single values for relative humidity and ambient temperature. Other than their size, we have not yet characterized physical features of droplet nuclei that could influence their efficiency in reaching distal airways. Finally, we have not proved that knockout or knockdown of any one gene in the candidate transmission survival genome of Mtb will impair Mtb’s infectivity upon inhalation of micro-aerosols by an experimental animal. Such studies require the design, construction, validation and installation of new kinds of equipment in BSL3 facilities. Those efforts are underway.

## Author Contributions

SM, PRS, XH, LLQ, RJ, XJ, and LG performed experiments. TH, KR, MZ and VD contributed key information. NS contributed to data analysis and modeling. AJ, MADJ, NS, RMJ and RAM provided computational analyses. CN, LB, SM, XH and AJ wrote the manuscript. LB designed and developed instrumentation and associated analysis methods. CN and LB designed and supervised the project. KR, CN, LB, and SM secured funding.

## Acknowledgments

We thank L. Via and C. Barry (NIAID, NIH) for providing specimens and the following people for generously sharing information about caseum, related materials and transcriptional responses of Mtb: H. Boshoff (NIAID, NIH), D. Russell (Cornell University), N. Walters (University of Colorado), D. Sherman (University of Washington); and J. Sarathy and B. Prideaux (Center for Discovery and Innovation, Hackensack Meridian Health). M. Wells (Cornell) provided statistical advice. Jeremy Rock (Rockefeller University) provided the Mtb CRISPRi library and *clpC1*/*clpP1P2* CRISPRi conditional knockdown strains. D. Schnappinger (Weill Cornell Medicine) advised on the construction of CRISPRi knockdown strains. F. Kaya (Center for Discovery and Innovation, Hackensack Meridian Health) helped identify deoxyphinganines. K. Saito and S. Ehrt (Weill Cornell Medicine) helped in diverse ways. J. Roberts, C. Suh, A. Lee, E. Kaplan and A. Singh (Weill Cornell Medicine) assisted in an experiment that required multiple personnel. Prof. T. Heldt and Prof. M. Gray (MIT) contributed to discussions and analysis. Prof. A. Singh (Indian Institute of Science) kindly shared the Mtb strain Mtb-MrxroGFP. K. Burns-Huang (Weill Cornell Medicine), Alison Fay (Sloan Kettering Institute) and Michael Glickman (Sloan Kettering Institute) provided additional Mtb strains. K. Burns-Huang and Ben Gold (Weill Cornell Medicine) provided administrative help. We thank J. Xiang of the Weill Cornell Genomics Resources Core Facility for extensive sequencing and analysis. This work was supported by NIH grant P01AI159402, the Abby and Howard P. Milstein Program in Chemical Biology and Translational Medicine and a grant to SM from the Potts Memorial Foundation. The Department of Microbiology & Immunology at Weill Cornell Medicine is supported by the William Randolph Hearst Foundation. The Fluid Dynamics of Disease Transmission laboratory (MIT) is also partially supported by CDC-NIOSH, INDITEX, NASA-TRISH and Analog Devices, Inc. All data needed to evaluate the conclusions in the paper are present in the paper, the supplementary materials and/or https://doi.org/10.7298/e4xs-y663.

