## Supplementary Materials for "Candidate transmission survival genome of *Mycobacterium tuberculosis*"

### Genes required by *Mycobacterium tuberculosis* to survive transmission

[1]Department of Microbiology and Immunology, Weill Cornell Medicine, New York, NY 10065; [2]The Fluid Dynamics of Disease Transmission Laboratory, Fluids and Health Network, Massachusetts Institute of Technology, Cambridge, MA 02139; [3]Skaggs School of Pharmacy & Pharmaceutical Sciences, University of California San Diego, San Diego CA 92093-0021; [4]Department of Chemistry and Biochemistry, University of California San Diego, San Diego CA 92093-0021; [5]Laboratory of Host-Pathogen Biology, Rockefeller University, New York, NY 10021; [6]Department of Medicine, Weill Cornell Medicine, New York, NY 10065; [7]Center for Discovery and Innovation, Hackensack Meridian Health, Nutley, NJ 07110; [8]Department of Microbiology, University of Washington, Seattle, WA 98195

*co-first authors

**Outline**

Materials, methods and theoretical analyses of droplet dynamics

Supplementary figures S1*A-D*, S2*A-C*, S3*A-M*, S4*A-D*, S5*A-B*, S6

Supplemental tables S1*A-C*, S2*A-E*, S3, S4

References S1-S54

#### Materials, methods and theoretical treatment

##### Microbial cultures

*Mycobacterium tuberculosis* (Mtb) H37Rv strains were cultured in Middlebrook 7H9 medium (BD Biosciences) with 10% (vol/vol) Middlebrook OADC (oleic acid, bovine albumin, dextrose, catalase) (BD Biosciences), 0.2% glycerol, and 0.02% tyloxapol (Sigma-Aldrich), hereafter referred to as complete 7H9, at 37°C under 5% $CO_2$ and ambient $O_2$, except for the CRISPRi library. The CRISPRi library was grown in 7H9 containing 0.2% glycerol, 0.05% Tween-80, OADC, and kanamycin 20 μg/mL, referred to as 7H9-Tween. Middlebrook 7H10 or 7H11 medium supplemented with 0.5% glycerol and 10% (vol/vol) OADC was used for assays of colony-forming units (CFU). Selected strains required antibiotics for selection (kanamycin [kan] 20 μg/mL; hygromycin [hyg] 50 μg/mL; Zeocin [zeo] 25 μg/mL) and were used as follows: *ΔprcBA*, hyg; *clpP1P2* knockdown, zeo, hyg, kan; *dnaJ2* knockdown, kan; *groEL2* knockdown, kan; *otsA* knockdown, kan; *otsB2* knockdown, kan; *devR* knockdown, kan; H37Rv-GFP, hyg; BCG-GFP, none.

##### HPLC-MS of deoxysphinganine

Deoxysphinganine lipid (DSGA) was quantified using laser capture microdissection on Mtb-infected rabbit (HN878) and marmoset lungs (H37Rv) followed by high performance liquid chromatography tandem mass spectrometry (LCM-LC-MS/MS) using published methods[S1]. Briefly, a 25 μm-thick section from an infected lung was sectioned on a Cryostat and subsequently a 3e6 $\mu m^2$ region was collected in caseous lung regions using a Leica LMD6500. A neat standard curve was created using the stable labeled analog DSGA-d3 (Avanti Polar Lipids). A DSGA-d3 surrogate was used in the standard curve due to the endogenous presence of DSGA. The neat standards were spiked into drug-free control caseum to create a standard curve. DSGA was extracted from the sections and spiked standards using methanol containing 20 ng/mL of the internal standard Verapamil. Extracts were vortexed for 5 minutes and centrifuged at 4000 RPM for 5 minutes. Supernatant was transferred for HPLC-MS/MS analysis. LC-MS/MS analysis was performed on a Sciex Applied Biosystems Qtrap 6500+ triple-quadrupole mass spectrometer coupled to a Shimadzu Nexera X2 UHPLC system. Chromatography was performed on an Agilent Zorbax SB-C8 column (2.1x30 mm; particle size, 3.5 μm) using a reverse phase gradient. Milli-Q deionized water with 0.1% formic acid was used for the aqueous mobile phase and 0.1% formic acid in acetonitrile for the organic mobile phase. We conducted multiple-reaction monitoring (MRM) of parent/daughter transitions in electrospray positive-ionization mode. The following mass transitions were monitored in MRM acquisition mode for DSGA (286/268), DSGA-d3 (289/271), and Verapamil (455.30/165.00). Sample analysis was accepted if the concentrations of the quality control samples were within 20% of the nominal concentration. Data processing was performed using Analyst software (version 1.6.2;

Applied Biosystems Sciex). Total ion chromatograms (TIC) and fragmentation patterns for lipid confirmation were acquired from caseum extracts and the synthetic DSGA standard (Avanti Polar Lipids) using a Thermo Q-Exactive high-resolution mass spectrometer (QE-HRMS) at 70000 mass resolution coupled to an Ultimate 3000 UHPLC.

**Metabolomics estimation**

Metabolomic analysis of caseous lesions was as described[S2]. For profiling, samples of human tissues from surgery subjects that contained caseous necrotic TB lesions were generously provided by Laura Via (Tuberculosis Research Section, Bethesda, MD). Caseous lung lesions were prepared by mechanical lysis in −20°C acetonitrile:methanol:H2O (40:40:20) using a Precellys tissue homogenizer under continuous cooling at 2°C. Biomass of each sample was determined by weight. Metabolite abundances were calculated using the method of standard addition with authentic chemical standards and normalized to residual protein concentration. Lesional metabolite concentrations were calculated assuming a lung tissue density of approximately 1 g/mL. Absent a validated internal standard to determine absolute recovery rates, the reported metabolite abundances and concentrations likely represent underestimations.

**PMN (polymorphonuclear leukocyte) lysate**

Anticoagulated blood from healthy adults was purchased from the New York Blood Center under Weill Cornell Medicine Institutional Review Board protocol 0806009836. 20 mL of blood was layered over 20 mL of Polymorphprep (density = 1.113 ± 0.001 g/mL) (Cosmo Bio USA, Inc.) in a

50 mL tube and centrifuged at 600 g for 30 min at room temperature. The rotor was allowed to decelerate without the brake and the band containing PMN (below plasma and mononuclear cells) was harvested using a Pasteur pipette. One volume of the PMN suspension was mixed with an equal volume of a 1:1 mixture of Hank's Balanced Salt Solution (--) (Gibco) and distilled water and centrifuged at 400 g for 10 min. The PMN pellet was resuspended in HBSS (--) before cell count and cells were lysed by 2 cycles of freezing at -20°C and thawing.

**Preparation of MAF**

Minimum Essential Medium (MEM)-α with nucleosides and no phenol red (Gibco|Thermo Fisher Scientific catalog no: 41061037) was used as a base. To this we added urea, sodium lactate, N,N-dimethylglycine hydrochloride and a 500X concentrate of the other metabolites listed in table S2c so as to give the final concentrations listed in the table. This mix was filter sterilized. For final preparation, the required volume was aliquoted in a glass tube or beaker to which we added the indicated lipids, phospholipids, serum, PMN lysate, deoxysphinganine, and guanine. This mix was sonicated at 30% amplitude using a probe sonicator for 10 seconds on and 20 seconds off for 30 minutes. Twenty minutes later, the cycle was repeated. We used two different versions of MAF in the study: the first was used for CRISPRi screens I and II, referred to as "$MAF_{screenI\&II}$," (table S1c). while the second was used for Screen III and all other experiments and is referred to only as MAF. $MAF_{screenI\&II}$ was used at the beginning of this work before we gained further knowledge of the composition of caseum.

**In vitro model of sequential stresses related to transmission**

Log phase bacterial cultures were grown to $OD_{580}$ = 0.4 - 0.8 in complete 7H9 and centrifuged in 50 mL Falcon (Corning) tubes (3098 x g, 8 minutes, RT). After discarding the supernatant, the pellet was suspended in 40 mL of phosphate-buffered saline (PBS) with 0.02% tyloxapol (PBS-tx), mixed and centrifuged (3098 x g, 8 minutes, RT). The supernatant was discarded and the pellet again suspended in 40 mL of PBS-tx. Cells were resuspended in MAF to an OD of 1.0, corresponding to 1 x $10^8$ cells/mL, and incubated under 0.2% $O_2$, 5% $CO_2$ at 37°C for two weeks, followed by 10% $O_2$ and 5% $CO_2$ at 37°C for two weeks. Cells were then desiccated in triplicate 2 μL droplets in wells of a 96-well polystyrene plate for the indicated durations in a closed plastic box whose floor was covered by Drierite under ambient atmospheric conditions. The cells were then rehydrated in 100 μL complete 7H9 for 20 minutes. Cells from each stage of the model were diluted in complete 7H9 to enumerate CFUs.

**Osmolality measurement**

Osmolality of media was measured using an Advanced Instruments model 3300 Micro-Osmometer.

**Drug tolerance assay**

Mtb cells resuspended in MAF at $OD_{580}$ = 0.8 were incubated under 0.2% or 21% $O_2$, 5% $CO_2$ at 37°C for 2 or 8 weeks and then treated with rifampicin (1 or 10 μM), moxifloxacin (1 or 10 μM), kanamycin (50 or 500 μM) and isoniazid (50 or 500 μM) with 1% DMSO as control for one week. Log phase replicating cells grown in complete 7H9 under 21% O2, 5% CO2 at 37°C were treated as controls. After one week, cells were serially diluted in complete 7H9, plated on 7H11 agar and incubated for 5 to 7 weeks at 37°C in ambient air before enumerating CFU.

**RNA sequencing (RNAseq)**

Log phase bacterial cultures were grown to $OD_{580}$ = 0.4 - 0.8 in complete 7H9, washed twice with PBS-tx, and resuspended in MAF at 1 x $10^8$ cells/mL. To model a closed cavity environment, cells resuspended in MAF were incubated under 0.2% $O_2$ and 5% $CO_2$ at 37°C for two weeks. As a control, we also incubated cells in MAF under 21% $O_2$ and 5% $CO_2$ at 37°C for 2 or 6 weeks. To model an open cavity environment, cells incubated under 0.2% $O_2$, 5% $CO_2$ at 37°C for two weeks were shifted to 10% $O_2$, 5% $CO_2$ at 37°C for two weeks. Additionally, we incubated cells in 7H9 at the same cell density under 0.2% or 10% $O_2$ and 5% $CO_2$ at 37°C for two weeks. Cells were then mixed with equal volumes of 5 M guanidinium isothiocyanate and centrifuged (3098 x g, 8 minutes, 4°C). After discarding the supernatant, cells were resuspended in 1 mL of Trizol (Invitrogen) and subjected to bead-beating in the presence of 0.5 mL silica beads using a Beadbug microtube homogenizer (Benchmark Scientific) at the 400 speed setting for 1 minute, followed by 2 minutes on ice, repeated five times. The tubes were centrifuged at (13,523 x g, 10 minutes, 4°C), and the supernatant was transferred to a fresh tube and mixed with 200 mL of chloroform before bringing samples out of the biosafety level 3 facility (BSL3). Samples were mixed by shaking, allowed to stand for 5 minutes at RT and centrifuged (13,523 x g, 15 minutes, 4°C). The aqueous layer was transferred to a fresh tube and mixed with one volume of ethanol before proceeding with the Direct-zol RNA miniprep protocol (Zymo Research) as per the manufacturer's protocol. Samples were treated with DNase using the TURBO DNA-free™ Kit (Catalog number:

AM1907) and RNA quantitated using NanoDrop. Sample libraries were prepared for RNA-seq by the WCM sequencing core using Illumina Stranded Total RNA Prep with Ribo-Zero Plus according to the manufacturer's protocol and subjected to Illumina sequencing. The sequencing libraries were sequenced with paired-end 100 bps on NovaSeq6000 sequencer. The raw sequencing reads in BCL format were processed through bcl2fastq 2.20 (Illumina) for FASTQ conversion and demultiplexing. After trimming the adaptors with cutadapt (version1.18) (https://cutadapt.readthedocs.io/en/v1.18/), RNA reads were aligned and mapped to the *Mycobacterium tuberculosis* genome (H37Rv) by Bowtie 2 version 2.2.8 (http://bowtie-bio.sourceforge.net/bowtie2/index.shtml). Raw read counts per gene were extracted using HTSeq-count v0.11.2. Gene expression profiles were constructed for differential expression, cluster, and principle component analyses with the DESeq2 package (https://bioconductor.org/packages/release/bioc/html/DESeq2.html). For differential expression analysis, pairwise comparisons were made between two or more groups using parametric tests where read-counts follow a negative binomial distribution with a gene-specific dispersion parameter. Corrected p-values were calculated based on the Benjamini-Hochberg method to adjust for multiple testing. The raw sequencing data files are submitted to NCBI under accession number PRJNA1140188.

**Fluorescence microscopy**

For imaging, desiccated droplets of Mtb or BCG strains expressing green fluorescent protein (GFP) resuspended in the indicated fluid were thoroughly mixed by pipetting ten times to break clumps. Then 2 μL aliquots were spotted in the wells of a 96-well plate. After desiccation for one day as described above, green fluorescent protein (GFP) visualized in the fluorescein isothiocyanate (FITC channel) and bright-field images were recorded with a Keyence microscope for media + Mtb-Mrx1roGFP2. Droplet residues of media and media + BCG-mEmeraldGFP were imaged using a Nikon Ti-U microscope.

**CRISPRi screens**

An Mtb CRISPRi library (RLC0012; containing ~99,000 sgRNAs) kindly shared by J. Rock, Rockefeller University was inoculated in 7H9-Tween containing 20 µg/mL kanamycin. The culture was grown at 37°C under 5% $CO_2$ and ambient $O_2$ to $OD_{580}$ = 1.0 and then diluted back to $OD_{580}$ = 0.05 in 7H9-Tween containing kanamycin 20 µg/mL and treated with 100 ng/mL of anhydrotetracycline (Atc). Cells were then grown to an $OD_{580}$ of ~ 1.5 to create cultures that had expanded for 5 generations under conditions that could select against knockdown of any given gene. Cells were then again diluted back to $OD_{580}$ = 0.05 and allowed to grow for another five generations in the presence of 100 ng/mL of Atc until $OD_{580}$ ~ 1.5 to create 10-generation-depleted cultures. Similarly, cells were grown in the presence of 100 ng/mL of Atc to create 15- and 20-generation-depleted cultures. Before starting the CRISPRi screen (input control) and after 5, 10 and 20 generations of depletion, 10 OD units of cells were harvested for next generation sequencing. We carried out three different CRISPRi screens as we gained experience, as follows:

***Screen I (for stage 1 only: closed cavity)***: For the first screen, cultures depleted for 5, 10 and 20 generations were spun down and washed twice with PBS-tx (3098 x g, 8 minutes, RT). Cells were resuspended in MAF at final $OD_{580}$ = 1.0 and incubated under 0.2 % $O_2$ and 5 % $CO_2$ at 37°C. After one week, one part of the culture was spun down and processed for DNA isolation and sequencing. The remainder was inoculated in 7H9-Tween containing 20 µg/mL of kanamycin and grown at 37°C under 5% $CO_2$ and ambient $O_2$ until OD = 1.0 before DNA isolation and sequencing.

***Screen II (progressed from stage 1 but analyzed only for stages 2 and 3: open cavity and desiccation)***: For the second screen, cultures depleted for 5 and 10 generations were spun down and washed twice with PBS-tx (3098 x g, 8 minutes, RT). Cells were resuspended in MAF at final $OD_{580}$ = 1.0 and incubated under 0.2 % $O_2$ and 5 % $CO_2$ at 37°C. After one week, cells were shifted to 10% $O_2$ and 5% $CO_2$ at 37°C and incubated for another week. The culture was then divided into three parts. The 1st part was processed for DNA isolation and sequencing as above. The 2nd part was inoculated in 7H9-Tween containing kanamycin 20 µg/mL and grown at 37°C until $OD_{580}$ = 1.0 before DNA isolation and sequencing. The 3rd part was used to set up desiccation by robotically dispensing 2 µL droplets in wells of 384 well plates (1 plate per replicate) followed by incubation above a layer of Drierite in air at RT. One and 3 days later, wells in different plates were filled with 7H9-Tween containing kanamycin 20 µg/mL and incubated at 37°C to allow growth to $OD_{580}$ = 0.4-0.8 to ensure a larger biomass. At each stage, cells were processed for DNA isolation and sequencing.

***Screen III (for stages 1, 2, 3: closed cavity, open cavity, and desiccation)***: For the third screen, cultures depleted for 5, 10 and 20 generations were spun down and washed twice with PBS-tx (3098 x g, 8 minutes, RT). Cells were resuspended in MAF at final $OD_{580}$ = 1.0 and 2 sets were incubated under 0.2 % $O_2$ and 5 % $CO_2$. After two weeks, one set was shifted to 10 % $O_2$ and 5 %

$CO_2$ and incubated for another two weeks before DNA isolation. Cells from the second set were divided into two parts. The 1st part was processed for DNA isolation. The 2nd part was inoculated in 7H9-Tween containing 20 µg/mL of kanamycin and grown till $OD_{580}$ = 1.0 and then harvested. The 3rd part was used to set up desiccation in 2 µL droplets in single-well plates (15 plates per replicate) and allowed to dry over Drierite. One day later, cells were rehydrated in MEM-α (15 mL per plate) for 20 minutes and scraped from the plates. Replicates were pooled and divided into two parts. The 1st part was processed for DNA isolation immediately. The 2nd part was grown in 7H9-Tween containing kanamycin 20 µg/mL to $OD_{580}$ = 0.4-0.8 before isolation of DNA.

**Genomic DNA extraction and sequencing**

For every replicate, 10 $OD_{580}$ units of cell pellets were collected and stored at -80°C until processing. Pellets were thawed, resuspended in 0.8 mL of Tris buffer (Qiagen) containing lysozyme (15 mg/ml and incubated at 37°C for 1 day for cells grown in 7H9-Tween or 2 days for samples coming out of MAF, because cells that had been incubated in MAF took longer to lyse. Each sample then received 70 µL of 10% SDS and 5 µL of Proteinase K (20mg/mL). After incubation at 65°C for 60 mins, samples received 100 µl of 5 M NaCl and 80 µl of 10% cetyltrimethylammonium bromide. After further incubation at 65°C for 60 mins, samples received 750 µL of chloroform, were shaken vigorously, left at RT for 15 mins and centrifuged at 13,523 x g, 5 min, RT. The upper phase was transferred to a 2 mL clean screw-capped tube and incubated with 25 mg RNase at 37°C for 60 mins. After an equal volume of phenol:chloroform: isoamyl alcohol was added, the tubes were mixed by inversion and centrifuged at 13,523 x g, 5 min, RT. Supernatant was transferred supernatant to a new tube and mixed with an equal volume of chloroform. After the contents were mixed and centrifuged as above, the supernatant was transferred to a fresh tube, and mixed with 0.6 volumes of isopropanol followed by 1/10 volume of 3 M sodium acetate, pH 5.5. DNA precipitated during overnight storage at -20 °C. Samples were centrifuged at 13,523 x g for 45 mins at 4°C, the supernatant was discarded, the pellet was washed with 70% ethanol and centrifuged again at 13,523 x g for 15 mins at 4 °C. The supernatant was discarded and the pellet allowed to dry in air. The DNA was dissolved DNA in water and quantitated using NanoDrop. To amplify the sgRNA-encoding region, 500-1000 ng genomic DNA was used as a template with 17 cycles of PCR using NEBNext Ultra II Q5 master mix (NEB M0544L). For Screen I, we used single indexing PCR where the reaction included a pool of forward primers (0.5 µM final concentration) and a unique indexed reverse primer (0.5 µM). For Screens II and III, dual indexing PCR was performed with a unique indexed forward primer (0.5 µM) and reverse primer (0.5 µM). PCR amplicons were purified using AMPure XP beads (Beckman–Coulter A63882) according to the manufacturer's protocol. Purified amplicons were quantitated using Qubit™ dsDNA HS Assay Kit (Catalog number: Q32851) and analyzed for amplicon size and purity on an Agilent 2100 bioanalyzer (high sensitivity chip; Agilent Technologies 5067–4626). Samples were pooled and sequenced on an Illumina sequencer according to the manufacturer's instructions. The raw sequencing data files for CRISPRi screens I, II and III are submitted to NCBI under accession numbers PRJNA1208037, PRJNA1208562 and PRJNA1209129.

**Gene essentiality analysis using MAGeCK, volcano plot visualization, and sliding window analysis**

The essentiality analysis of *Mycobacterium tuberculosis* genes was based on output from MAGeCK, a statistical framework used to analyze raw read counts from pooled CRISPRi screens. We extracted the $\log_2$ fold-change (LFC) values and Benjamini-Hochberg corrected p-values from the MAGeCK output files across multiple experimental conditions. During manual quality control, two sample sets (belonging to stage 3) were discarded due to missing values or zero LFC values, indicating experimental artifacts. Thresholds were applied for gene essentiality, with LFC set to $< -1$ and p-values $< 0.05$. For each condition, we calculated the number of essential genes under the condition tested by counting those passing the significance thresholds, and these counts were aggregated across all comparisons. The results were visualized using averaged volcano plots, where the bubble size reflects the number of times a gene was classified as essential across the multiple comparisons used to measure fitness effects in a given stage of the transmission model. A sliding window analysis further assessed the fraction of genes deemed essential in MAGeCK screens that had previously been identified as essential under standard in vitro conditions, providing additional insights into gene fitness under host-like conditions.

**CRISPRi knockdown strains**

To create individual CRISPRi constructs, we used plasmid pIRL58 (Addgene cat log no 166886) and followed the procedure outlined by[S3]. The plasmid was initially digested with BsmBI-v2 (NEB 0739L) and purified by gel electrophoresis. Single guide RNAs (sgRNAs) were designed to target the non-template strand of the gene of interest, considering the presence and importance of the PAM sequence for both essential and non-essential genes. For each sgRNA, two complementary oligonucleotides were synthesized, each containing compatible sticky ends (table S4). These oligonucleotides were annealed and then ligated into the BsmBI-digested plasmid backbone using T4 ligase (M2200L). The resulting constructs underwent Sanger sequencing to ensure the successful insertion of the sgRNA sequences. For transformation, we expanded the Mtb culture to an $OD_{580}$ of 0.8 to 1.0 and then pelleted the cells (3098 x g, 10 minutes, RT). The pellet underwent three washes in sterile 10% glycerol and bacilli were resuspended in 10% glycerol to a final volume equivalent to 5% of the original culture volume. For each transformation, 200 ng pIRL19 (Addgene plasmid cat log no 163634) was included along with 200 ng sgRNA-containing pIRL58 plasmid DNA, mixed with 400 µL of electrocompetent mycobacteria and transferred to a 2 mm electroporation cuvette (VWR 89047-208). The Gene Pulser X cell electroporation system (Bio-Rad 1652660) was used at 2500 V, 700 Ω, and 25 µF. Bacteria were then incubated in 7H9 complete for 24 hours and plated on 7H10 agar supplemented with the appropriate antibiotic to select transformants.

**Fluid densities and surface tensions**

Density and surface tension were measured with a tensiometer (Biolin Scientific Sigma 701) equipped with a spherical buoyancy probe and a du Noüy ring. A spherical buoyancy probe was suspended in the fluid of interest. The measured apparent weight $W_a$ of the suspended probe determines the fluid density $\rho = (W - W_a)/gV$, where $V$ is the probe volume, $W$ is the real probe weight and $g$ is the gravitational acceleration.

To measure surface tension, the tensiometer was equipped with a platinum du Noüy ring[S4] with radius $R = 9$ mm. The ring was suspended in a beaker with 50 mm radius that contained 8 mL of fluid. The ring was pulled until a fluid meniscus formed. The pulling force was measured until a maximum force $F_{max}$ was reached without breaking the meniscus. The surface tension $\gamma$ was calculated by the device's software based on its relation to the maximum pulling force $F_{max}$ as $\gamma = F_{\max}/(4\,\pi\,R)\,\psi$, where $\psi$ is a correction factor that depends on the ring geometry.[S5, S6] We repeated the pulling procedure 10 times per measurement.

**Shear viscosity**

We determined the shear viscosity of MAF, MEM–$\alpha$, 6 salts, MAF + BCG, and water as reference using a high-precision rotational rheometer (TA-instruments HR-1) with a cone and plate geometry, cone radius $R = 30$ mm, and cone contact angle $\theta = 1°$. The cone tip was cut, resulting in a truncation gap of $a = 33\ \mu m$. The cone was rotated with angular velocity $\Omega$ and the applied torque $M$ measured. For Newtonian fluids the viscosity can then be determined by $\eta = 3\theta/(2\pi R^3\,\Omega)\,M$ where we assume no edge effects and small cone angles.[S7] The plate was maintained at $20°C$ and covered to prevent evaporation during measurements. The measurements were performed for a range of shear rates $\dot{\gamma} = 10^1 - 10^3\ 1/s$. Fig. S1*D*(b) shows the measurement results and confidence region with <1% measurement method-induced error.

The confidence region is bounded: At high shear rates, secondary flow can spuriously increase the apparent fluid viscosity. For the geometry used here and measured torque, $M$, the secondary flow for Newtonian fluids is $\frac{M}{M_0} = 1 + \frac{3}{4900} Re^2$, where $M_0$ is the ideal torque and $Re = \frac{\rho \Omega L^2}{\eta}$ is the Reynolds number with characteristic length $L = \alpha R$,[8] $\rho$ is the density of the fluid, $\Omega$ is the angular velocity of plate rotation, and $\eta$ is the dynamic viscosity of the fluid. We chose a critical Reynolds number $Re_{crit} = 4$ below which errors due to secondary flow were less than

1%, i.e., $\frac{M}{M_0} < 1.01$; At low shear rate, the minimal measurable torque $M_{min} = 0.1\mu N \cdot m$ for our rheometer. The condition $M > M_{min}$ defines the low torque limit.[S8] For the cone plate considered here, $\eta = \frac{3M}{2\pi R^3 \dot{\gamma}}$. Thus, the low shear rate limit reads $\dot{\gamma} > \frac{3M_{min}}{2\pi R^3 \eta}$. However, the apparent low torque limit is larger than the instrument specification, typically due to surface tension effects.[S9] Therefore, we chose to limit the experimental window to 20 times the minimal torque limit specified by the manufacturer, consistent with prior studies.[S10-12]

The measurement results shown in Fig. S1*D*(b) indicate that the viscosity did not change significantly with shear rate for the fluids considered here, remaining in a Newtonian limit. Therefore, we report the mean value of shear viscosity of the fluids, within the confidence range. The viscosity measurement results show that MEM-$\alpha$ and the 6 salts fluids have similar viscosity to that of water, i.e., $\eta = 1mPa \cdot s$ at 20℃.[S13] The model aerosol fluid (MAF) has a slightly higher viscosity. Introducing BCG at $10^8$ CFU/mL did not alter the viscosity significantly. Table S3 lists the mean and sample standard deviation of the shear viscosity measurements as shown in Fig. S1*D*(b).

**Side imaging of evaporation**

We monitored evaporation of droplets by time-lapse imaging in an acrylic chamber containing Drierite (Fisher Scientific NC9979539) to keep the relative humidity ~ 5%. Relative humidity and temperature during evaporation were recorded by Sparkfun si7021 sensors. The droplets were illuminated by LED back-light and photographed from the front with a Nikon D3300 camera equipped with 2 2x Nikon TC-201 2x Manual Focus Teleconverters and a Nikon Telephoto AF Micro Nikkor 105mm f/2.8D lens. The cameras were connected to a computer and controlled by

DigiCamControl to take time-lapse photos. Analysis by MATLAB extracted geometrical parameters, including volume, surface area, contact radius, and contact angle.

**Preparation of polystyrene surfaces for biophysical studies**

To match the surfaces used in the biologic studies, we cut the bottom of polystyrene single well plates (Celltreat 229101) into 10 X 10 mm pieces with a laser cutter (GCC Spirit LS 12-100W CO2 Laser Engraver). To avoid deposition of vaporized polystyrene, we optimized the following parameters: 20% speed, 100% power, 3 repeats, resulting in partial penetration that allowed pieces to be detached by gentle manual pressure on the outer surface. Untouched inner surfaces were cleaned with an air stream to remove any dust before using.

**Profilometry**

We measured the height profiles of residues from evaporated droplets using an Optical Interferometric Profiler (Bruker ContourX 500). We took scans using 20x and 50x lenses with 0.55x magnification at scan speed 5x and analyzed the data with a MATLAB script.

**Scanning electron microscopy (SEM) and energy-dispersive X-ray spectroscopy (EDS)**

Samples for SEM were sputter coated with an 8 nm-thick layer of Au and imaged using the SE2 detector with EHT = 15 keV.  EDS spectra were collected at EHS = 15 keV with ab Oxford AZtec 100 EDS Detector. Samples without BCG were imaged using the MIT NANO Gemini 450 SEM. Samples with BCG were imaged at the MIT Nanotechnology Materials Core in the Koch Institute.

**Additional characterization of desiccating droplets**

*Composition of media*

The 36 water-soluble metabolites found in human caseum are listed in table S1*A*. MAF is made by mixing subgroups of media at specific volume fractions with supplements, whose summary can be found in table S2*A*, and the detailed composition can be found in tables S2*B*-S2*F*. The subgroups of media include the MEM-$\alpha$ aqueous base, lipid and phospholipid mix, serum, PMN lysate, and metabolite mix. For each ingredient in the subgroup, the volume fraction of the non-volatile solute can be calculated as $\phi = \frac{c_m}{\rho}$, where $\rho$ is the density of the ingredient in solid form and $c_m$ is the final mass concentration of the ingredient in MAF. One exception is MEM-$\alpha$ with additional metabolites (except guanine), for which $c_m$ for 100% MEM-$\alpha$ is listed in table S2*B* and *S2C*. In this case, the volume fraction of 93% was accounted for when calculating the final concentration in MAF. The volume fraction of all non-volatile components in MAF is the sum of the individual constituent volume fractions for all ingredients. Key subgroups of MAF are selected as control groups with simplified ingredients. The two subgroups are MEM-$\alpha$ and the 6-salts in MEM-$\alpha$. The composition of 6-salts is listed in table S2*G*. The volume fraction of non-volatile components is summarized and compared with MAF in table S2*H*.

*Evaporation of droplets made of MAF, MEM-$\alpha$, 6 salts, and MilliQ water on polystyrene surfaces with cell culture treatment*

Time-lapse imaging of the sessile droplets allowed us to determine the volume change of the droplet during the rapid evaporation process (Phase I). When the characteristic length of a droplet of volume $V$, for example estimated by its equivalent radius $R_{eq} = \left(\frac{3V}{4\pi}\right)^{\frac{1}{3}}$, becomes smaller than the capillary length, $\sqrt{\frac{\gamma}{\rho\, g}}$, the sessile droplet adopts a quasi-spherical cap shape (Fig. S3*C*). Here $\gamma$ is the liquid-air surface tension, $\rho$ is the density of liquid, and $g$ is the gravitational acceleration.

When considering a quasi-static diffusion-driven evaporation process of the sessile droplet, the vapor concentration field $c$ follows

$$\Delta c = 0 \qquad (s1)$$

The evaporative flux $\vec{j}$, defined at the water-air interface (fig. S3c), is:

$$\vec{j} = -D\nabla \mathrm{c} \cdot \vec{n} \qquad (s2)$$

where $D$ is the diffusion coefficient of water vapor in air.

The boundary conditions are

$$\left.\frac{dc}{dz}\right|_{z=0} = 0 \quad \text{at the solid} - \text{air interface.} \qquad (s3)$$

Moreover, $RH = c(\infty)/c_s$ is the air relative humidity, $c(\infty) = c_\infty$ is the mass concentration of water vapor in the environment far away from the surface, and $c_s$ is the saturated mass concentration of water vapor. Note that $c_s$ is a function of temperature $T$ through the ideal gas law

$$c_s = \frac{M_w p_s(T)}{R(273.15 + T)} \qquad (s4)$$

where $M_w = 18\, g/mol$ is the molar mass of water, $R = 8.314\, J/(mol \cdot K)$ is the ideal gas constant, $T$ is temperature measured in ˚C, and $p_s(T)$ is the partial pressure of saturated water vapor in air at $T$. An empirical formula for $p_s(T)$ is the Arden Buck relationship[S14]

$$p_s(T) \approx 0.61121 \exp\left[\left(17.513 - \frac{T}{234.5}\right)\frac{T + 273.15}{T + 530.29}\right] \qquad (s5)$$

The magnitude of the flux $j = |\vec{j}|$ (see fig. S3c) can be solved as[S15]

$$j(r) = j_0 (R_c - r)^{-\lambda}, \qquad j_0 = \frac{D(c_s - c_\infty)}{R_d}, \qquad \lambda = \frac{\pi - 2\theta}{2\pi - 2\theta}\,, \tag{s6}$$

where $R_c$ is the contact radius, $R_d$ is the radius of the droplet, $\theta$ is the contact angle of the sessile droplet, and $j_0$ is the evaporation flux at the apex, i.e., $j_0 = j(r = 0)$. The total evaporation rate can be found by integrating over the flux[S16]

$$J = -\frac{dm}{dt} = \int_0^{R_c} j(r)\sqrt{(1 + (\partial_r h)^2}\, 2\pi r\, dr \; = \pi R_c D(c_s - c_\infty) f(\theta), \tag{s7}$$

where $m$ is the mass of the liquid droplet and $f(\theta)$ is given by

$$f(\theta) = \frac{\sin\theta}{1 + \cos\theta} + 4\int_0^{\infty} \frac{1 + \cosh 2\theta\tau}{\sinh 2\pi\tau} \tanh\big((\pi - \theta)\tau\big) d\tau\,, \tag{s8}$$

with $m = \rho_L V$, where $\rho_L$ is the density of the solvent, i.e., water, $V$ is the volume of the liquid droplet, and the volumetric evaporation rate is

$$\frac{dV}{dt} = -\frac{\pi D(c_s - c_\infty)}{\rho_L} R_c f(\theta) \tag{s9}$$

When the droplet's contact line is pinned during the evaporation, i.e., $(dR_c)/dt = 0$, the mode of evaporation is called constant contact radius (CCR), and the dynamics embodied in (s9) can be recast leveraging the dependence of $V$ and on $\theta$ as

$$V(R_c, \theta) = \frac{\pi R_c^3}{3} b(\theta), \qquad b(\theta) = (1 - \cos\theta)^2\,(2 + \cos\theta)\sin^{-3}\theta, \tag{s10}$$

which leads to

$$\frac{d\theta}{dt} = -\frac{D(c_s - c_\infty)}{\rho_L R_c^2} (1 + \cos\theta)^2 f(\theta). \tag{s11}$$

Then, the evaporation time, i.e., the time at which $\theta = 0$ is reached, is

$$t_{CCR} = t_F I(\theta_0), \qquad t_F = \frac{\rho_L R_{c0}^2}{D(c_s - c_\infty)}\ , \tag{s12}$$

where $\theta_0$ is the droplet's initial contact angle, $R_{c0}$ is the initial contact radius, and $I(\theta_0)$ is the integral

$$I(\theta_0) = \int_0^{\theta_0} \frac{d\theta}{(1+\cos\theta)^2 f(\theta)}. \tag{s13}$$

For $\theta \ll 1\, rad\ (i.e., \approx 57°)$, the evaporation time can be approximated by[S15]

$$t_{CCR} \approx t_{CCRL} = \frac{\pi \rho_L R_{c0}^2 \theta_0}{16 D(c_s - c_\infty)} \tag{s14}$$

where $c_s = c_s(T_\infty)$, $c_\infty = RH c_s(T_\infty)$, and $T_\infty$ is the far-field ambient temperature. The evolution of the corresponding volume is a linear function of time:

$$\frac{V}{V_0} = 1 - \frac{t}{t_{CCRL}} \tag{s15}$$

where $V_0$ is the initial volume of the liquid droplet. Introducing $(V_0)^{\frac{1}{3}}$ as length scale and $t_{CCRL}$ as the time scale, a non-dimensional form of Eq. (s15) can be written as

$$V^* = 1 - t^* \tag{s16}$$

with $V^* = V/V_0$ and $t^* = t/t_{CCRL}$.

*Effects of evaporation cooling on the rate of evaporation*

We compared the evaporation model developed above against the experimental measurements of droplet evaporation on polystyrene surfaces treated for compatibility with cell culture. The evolution of dimensional and normalized $V$, $R_c$, and $\theta$ is shown as a function of (dimensional time) $t$ in Fig. S3*D*. We found that the experimental evaporation is slower than the predictions given by Eq. (S16). One reason is that the theory developed above neglects evaporative cooling effects. In fact, for water vapor to diffuse and evaporate from a droplet, the liquid water needs to first vaporize. This vaporization consumes heat $Q = h_L m$, where $h_L = 2441.7 \frac{kJ}{kg}$ is the latent heat of water at $T = 25^o C$ and $m$ is the mass of water being vaporized.[S17] Evaporation therefore cools down the droplet to a temperature $T_d$ lower than the environmental temperature $T_\infty$ (Fig. S1*A*). A reduced droplet temperature leads to a reduced vapor saturation concentration $c_s(T_d)$ according to Eq. (*s*4). Defining $\sigma$ to be the saturation coefficient of water vapor of temperature $T_d$ as follows,

$$\sigma = \frac{c_s(T_d)}{c_s(T_\infty)}, \qquad (s17)$$

and substituting (*s*17) into (*s*14) gives the following nondimensional evaporation time of the droplet $t^*_{CCRL,thermal}$ that accommodates a droplet surface temperature, $T_d$ that is different from the ambient temperature $T_\infty$

$$t_{CCRL,thermal} = \frac{1 - RH}{\sigma - RH} t_{CCRL} \equiv \alpha t_{CCRL}. \qquad (s18)$$

With $\alpha \geq 1$ and here $T_d < T_\infty$. Thus, analogous to (*s*16):

$$V^* = 1 - \frac{t}{t_{CCRL,thermal}} = 1 - t^*/\alpha. \qquad (s19)$$

In Fig. S1*A*, we show the droplet surface temperature of an evaporating water droplet on the polystyrene substrate measured using an infrared camera. We found a maximum difference between the droplet and substrate around 6 – 7 ˚C, i.e., $T_d = T_{IR} \approx T_\infty - 6.5\,^\circ C$. Here, $T_{IR}$ denotes the infrared camera reading.

Alternatively, for a suspended droplet that evaporates in air, its surface temperature can be approximated by the Wet-Bulb temperature $T_{WB}$ that gives the lowest temperature reached under a given ambient condition by the diffusion-driven evaporation process of water only. $T_{WB}$ can be directly measured using a wet-bulb thermometer, or estimated using the following empirical relationship[S18]

$$T_{WB} = T\tan^{-1}\left(0.151977\,(RH+8.313659)^{\frac{1}{2}}\right) + \tan^{-1}(T+RH) - \tan^{-1}(RH-1.67331) + 0.00391838(RH)^{\frac{3}{2}}\tan^{-1}(0.023101RH) - 4.686035 \quad (s20)$$

In Fig. S3*D*, we also compare the theoretical droplet evaporation curves [based on Eq. (*s*19)] obtained using $T_d = T_{IR}$ and $T_d = T_{WB}$, respectively. We found that the model using $T_d = T_{IR}$ agrees best with the experimental results, as shown in Fig. S3*D*. The agreement is valid for media including MAF, MEM-$\alpha$, 6 salts, and water, with and without BCG at $10^8$ CFU/mL concentration. Also, we observed from Fig. S3*D* that the droplet contact radius remained constant for most of the droplet's lifetime, and therefore our assumption of the CCR evaporation mode is valid.

**Microscopic images of evaporated residues**

Microscopic images of evaporated residues from droplets that are made of MAF, MEM-$\alpha$, and 6 salts under bright field, phase contrast, and fluorescent FITC channels are summarized in Fig. S3*G*-S3*I*. The residue of MEM-$\alpha$ showed a concentrated center and a larger circle surrounding it, indicating that the droplet had receded from the initial position during the evaporation process. In contrast, the residues of both MAF and 6 salts did not show recession, i.e., the droplets remained pinned.

When we introduced BCG at a concentration of $10^8$ CFU/mL into the media, microscopic images of the residues that evaporated from droplets of media+BCG are illustrated in Fig. S3*H*. Residues of all droplets remained pinned during the evaporation, including MEM-$\alpha$. The fluorescent FITC images show that BCG was concentrated at the center of MAF residues while more distributed on the edges in the residues of MEM-$\alpha$, 6 salts, and water. Such distribution of BCG in residues qualitatively agrees with the distribution of Mtb as shown in Fig. S3*I*.

**Height profile of residues**

Axially averaged height profiles of residues evaporated from MAF, MEM-$\alpha$, and 6 salts are summarized in Fig. S3*E*. The residues of both MAF, MEM-$\alpha$, and 6 salts were higher at the edges than at the center due to coffee ring effect.[S19] The 6 salts' outer radial layer of the residues is detached from the substrate with regular – along the outer contour – patterns of buckling/wrinkling. These patterns reflect that an outer surface skin layer formed in the region of the foot of the evaporating droplet and then collapsed and deformed onto the surface as the drops fully evaporate also forming a gap with the substrate (Fig. S3*F*). The gap volume was quantified with CT imaging with corresponding profilometry height measurements and used to correct the volume of the residue.

**SEM images of evaporated residues**

In order to directly visualize the morphology of evaporated BCG and its location in the residue, detailed SEM imaging analysis was performed for residues evaporated from media + $10^8$ CFU/mL BCG, including MAF, MEM-$\alpha$, 6 salts, and water (Figs. S3*K*-S3*M*).

In residues of MAF + $10^8$ CFU/mL BCG, we did not find any BCG at the surface of the residues, indicating that all BCG cells were imbedded inside the residues (Fig. S3*M*). In the residues of MEM-$\alpha$ + $10^8$ CFU/mL BCG, both clusters of exposed BCG and flattened BCG are observed at various locations on the surface of the residue (Fig. S3*L*). In the residues of 6 salts and water + $10^8$ CFU/mL BCG, all BCG were exposed (Fig. S3*K*).

**Atomic chemical composition of residues using EDS analysis**

EDS analysis of residues evaporated from media only, including MAF, MEM-$\alpha$, and 6 salts are summarized in Fig. S3*J*. The chemical element maps show that the crystals in the residues were mostly sodium chloride, and oxygen appeared to extend widely through the residues of MAF.

**Computing hydrophilicity, glycine content and intrinsically disordered regions across the Mtb proteome.**

To search for hydrophillin-like proteins in the Mtb proteome, we employed a method[S20] that uses a two-descriptor approach: the average amino acid hydrophilicity index and the glycine content (normalized as a percentage of protein size). We combined these two with a third predictor, the ratio of residues predicted to form part of an intrinsically disordered region (IDR). The hydrophilicity averages of all proteins in Mtb were calculated employing the Parker algorithm[S21] with a window size of 2 as implemented in the Protein Analysis module in Biopython[S22]. The Parker scale consists of a set of experimental parameters obtained from retention times in high-performance liquid chromatography. In this scale, which ranges from -10 to 10, a positive value represents a hydrophilic residue, whereas more negative values represent more hydrophobic residues. The same tool and module were used to calculate the glycine content of each Mtb protein. To calculate the ratio of IDRs we used the SETH[S23] model, where each residue is associated with being in an IDR or not based on the value of CheZOD score, with a threshold of CheZOD<=8 as the definition of resides in an IDR.  All Python code for this and other analyses can be found in the following github repository: https://github.com/richiam16/MAF_Nathan

**Supplementary figures**

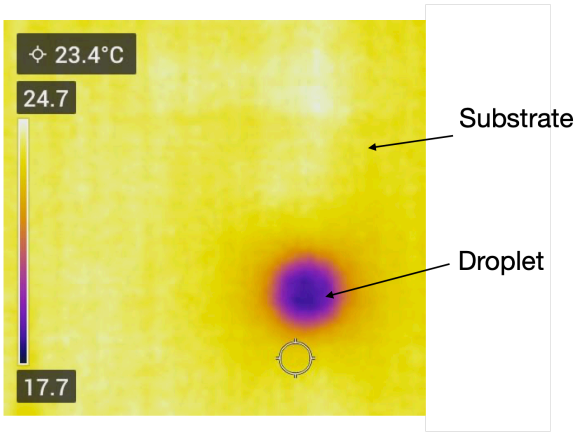


**Fig. S1*A*. Infrared image of an evaporating droplet on a polystyrene surface.** The temperature of the evaporating water droplet is about 6 – 7 ($\pm$ 3) °C cooler than the polystyrene surface.

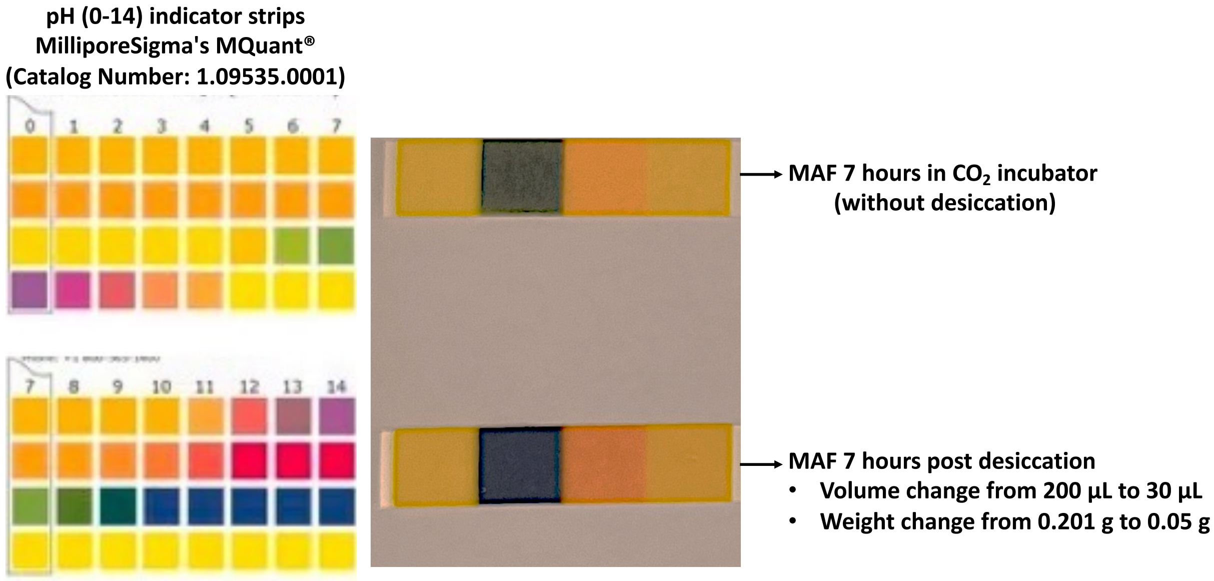


**Fig S1*B*. MAF alkalinizes as it desiccates in air.** Color of pH paper changes from greenish before desiccation to blueish (2$^{nd}$ square from left).

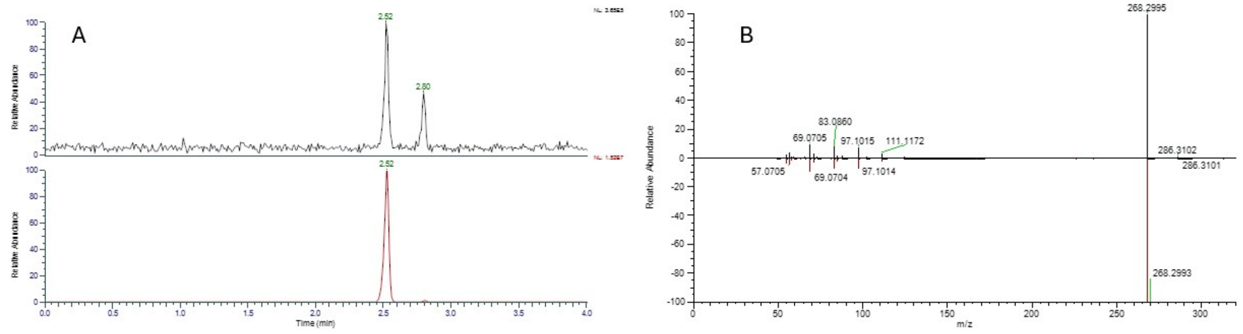


**Fig S1*C*. HPLC-MS confirmation of deoxysphinganine (DSGA) in caseum.** (A) The top panel illustrates the 5 ppm extracted ion chromatogram (XIC) of protonated DSGA in rabbit caseum and the bottom panel illustrates the XIC of the protonated DSGA mass in the synthetic DSGA standard dissolved in methanol. (B) The fragmentation mass spectrum of DSGA in rabbit caseum extract is illustrated above the x-axis and the fragmentation spectrum of the DSGA standard is illustrated below the x-axis.

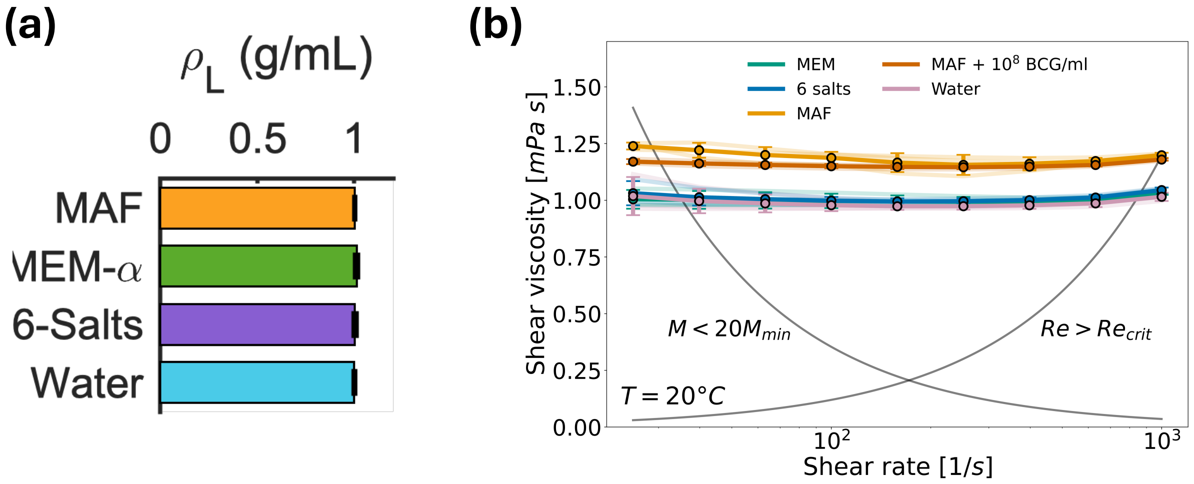


**Fig. S1*D*.** **(a)** Density, $\rho_L$, of the various fluids compared, showing that they are similar, consistent with their solute mass concentration of < 2%. **(b)** Shear viscosity as a function of shear rate for MAF (with and without $10^8$ CFU/mL of BCG), MEM-$\alpha$, 6 salts, and water at 20$^o$C. The shear rate dependent limits of the measurement window are shown. Within the measurement window, all fluids showed constant viscosity. The individual repeats are shown as translucent curves; the means are shown as darker lines. For both (a) and (b), the error bars indicate the sample standard deviation over three repeats.

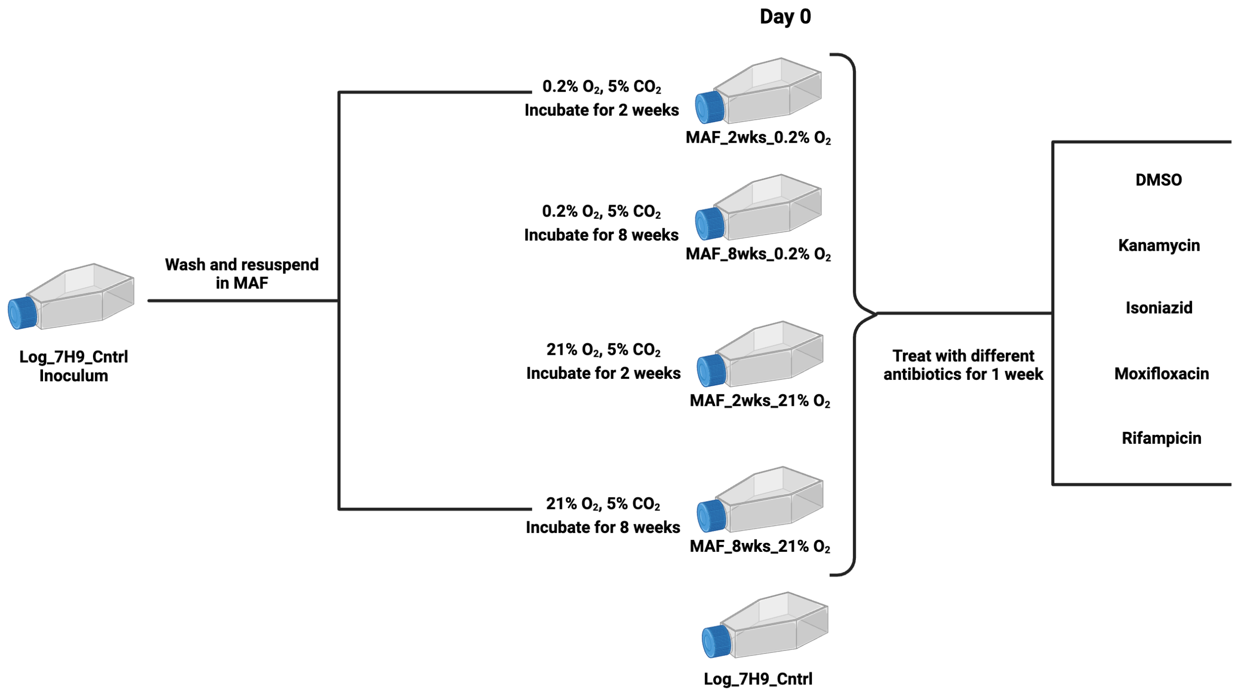


**Fig. S2*A*. Schematic of *Mtb*'s survival in MAF and drug tolerance assay under both 0.2% $O_2$, 5% $CO_2$ and 21% $O_2$, 5% $CO_2$** (Created with BioRender.com).

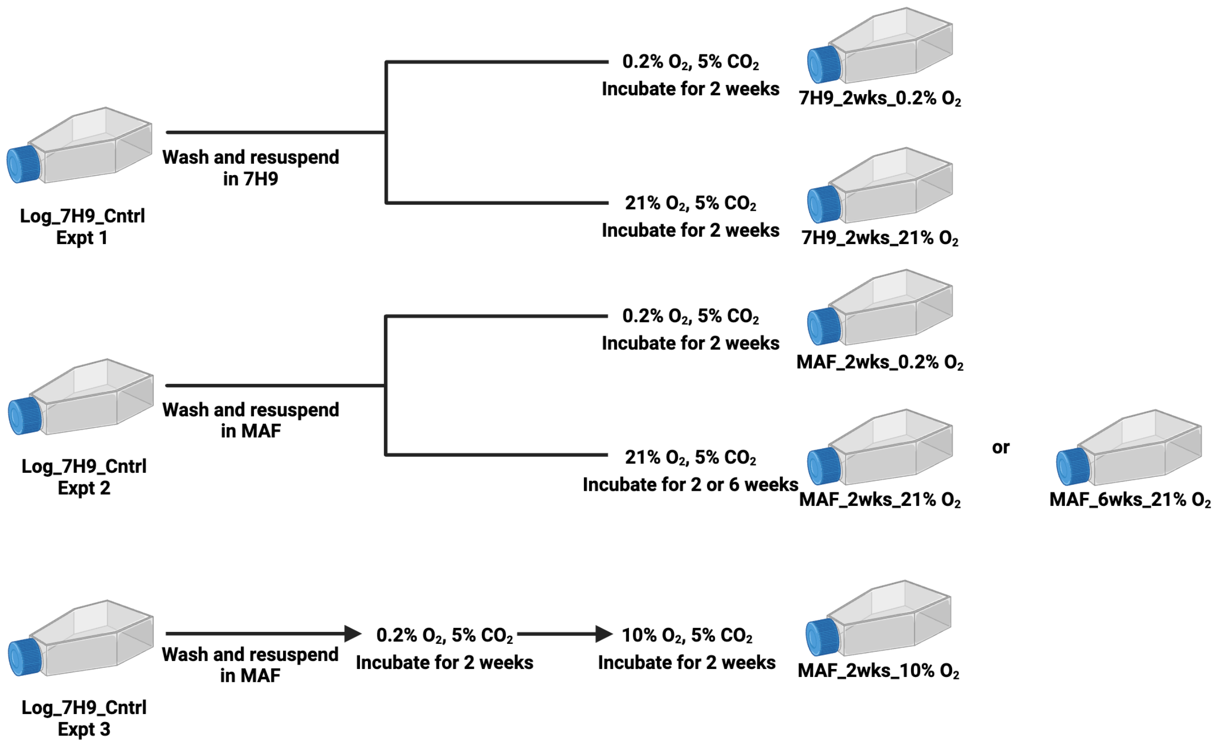


**Fig. *S2B*. Schematic of transcriptomic comparison of Mtb in MAF vs 7H9 under 0.2% $O_2$ or 21% $O_2$ (**Created with BioRender.com).

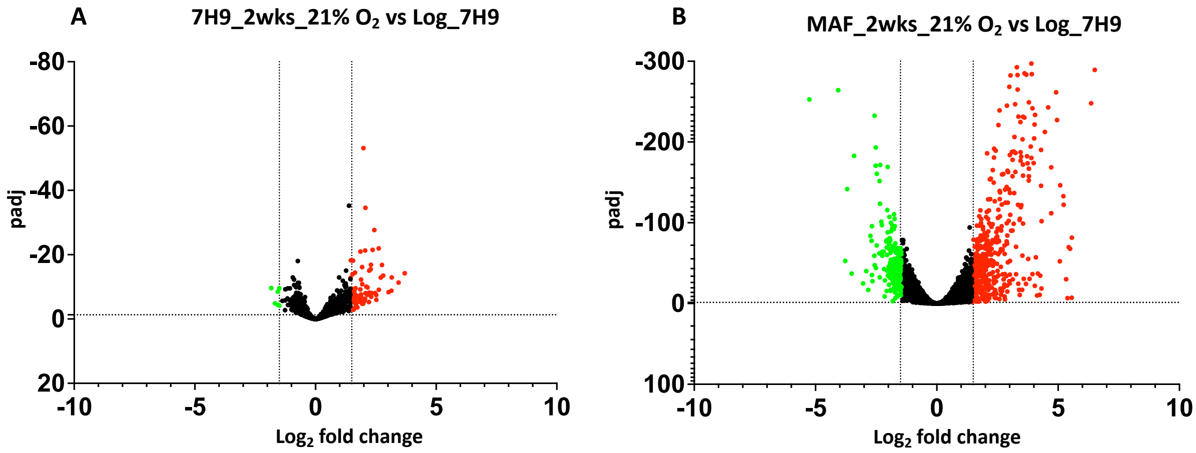

**Fig. S2*C*. Volcano plot of differentially expressed genes from Mtb incubated under 21 % $O_2$, 5 % $CO_2$ in 7H9 or MAF for 2 weeks versus a log phase replicating culture in 7H9 under 21 % $O_2$, 5 % $CO_2$.**

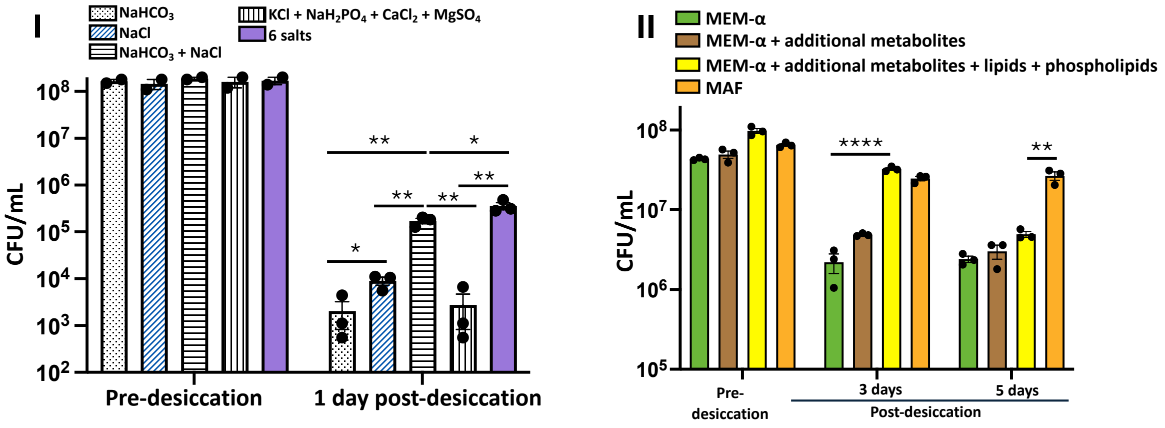


**Fig. S3*A*. Contribution of different components of MAF to MAF's promotion of Mtb's survival during desiccation.** (I) Contribution of different salts. (II) Contribution of additional metabolites, lipids, phospholipids, serum and PMN lysate. Data are means ± SEM of duplicates/triplicates in one experiment. P values were determined by unpaired t test. *P ≤0.05; **P ≤0.01; ****P ≤0.0001.

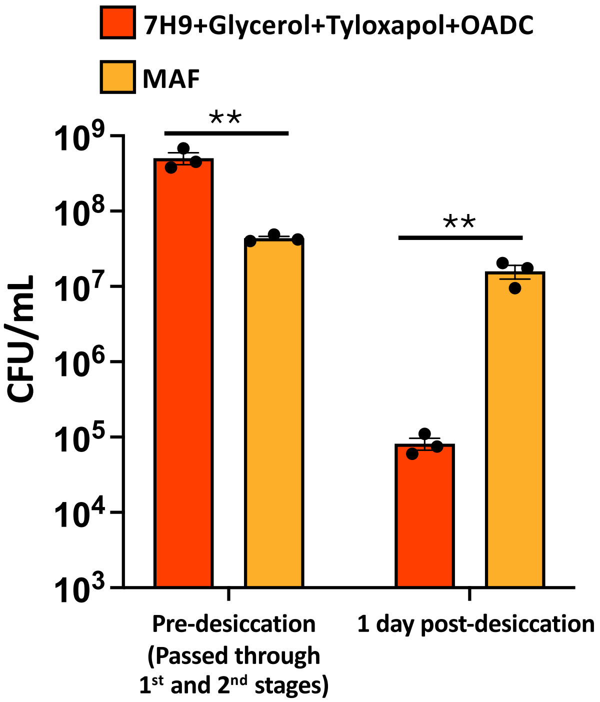


**Fig. S3*B*. Contribution of different components of MAF to MAF's promotion of Mtb's survival during desiccation after passage of Mtb through the first two stages of the screen.** Data are means ± SEM of triplicates in one experiment. P values were determined by unpaired t test. **P ≤0.01.

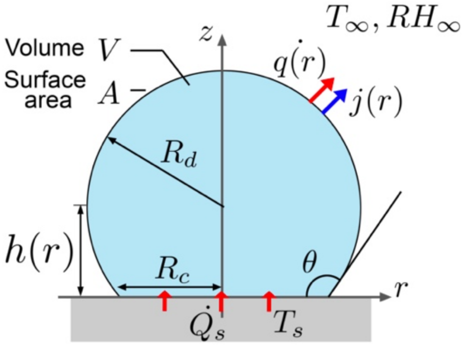

**Fig. S3*C*.** Schematic of an evaporating sessile droplet (blue) of volume V, area A, and wetting angle $\theta$, on a solid surface (grey). $R_c$ is the contact radius, and $R_d$ is the radius of the droplet surrounded by ambient with temperature, $T_\infty$, and ambient relative humidity, $RH_\infty$. Other geometric and flux variables are defined at the air-liquid and solid-liquid interfaces.

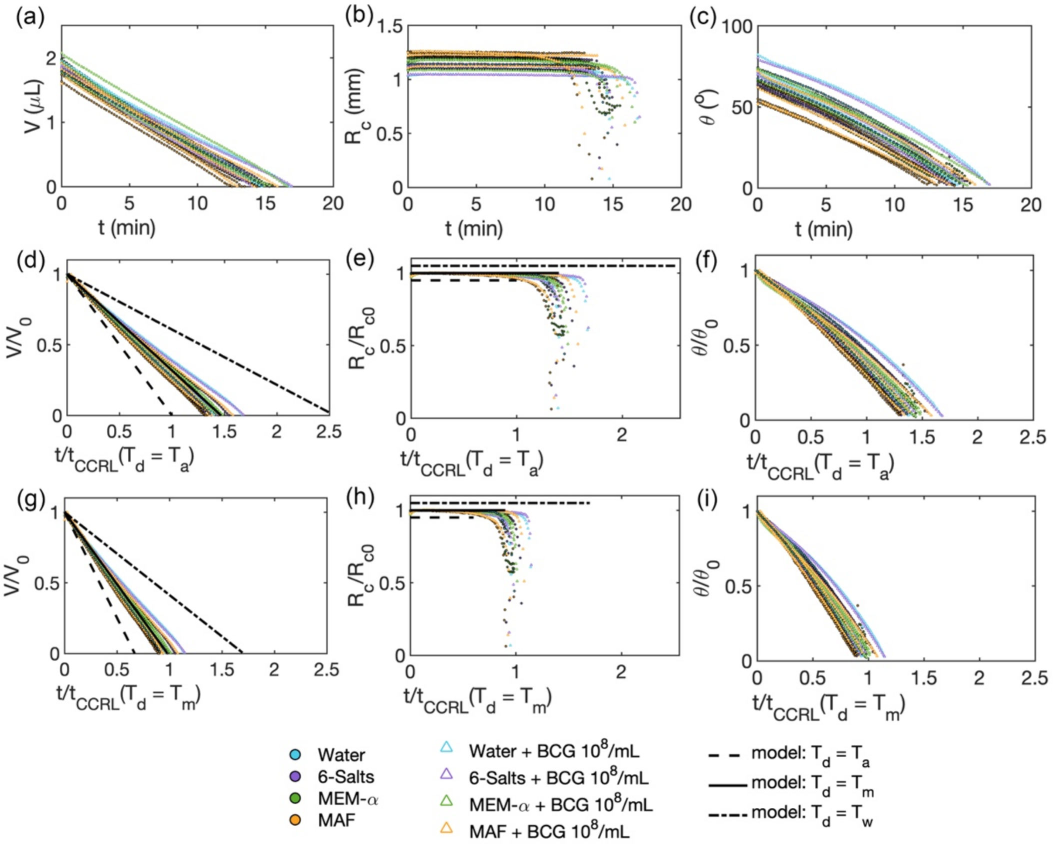


**Fig. S3*D*. Evaporation dynamics of sessile droplets made of various media, with and without BCG.** Evolution of experimentally measured $V$, $R_c$, and $\theta$ as a function of time $t$ (a - c). Note that $\frac{R_c}{R_{c0}} = 1$ is shifted for different models for visibility in panels (e) and (h). The evolution of the normalized quantities $V/V_0$, $R/R_{c0}$, and $\theta/\theta_0$ as a function of $t/t_{CCRL}$ when the droplet temperature $T_d$ is approximated by ambient temperature (d – f) and by experimentally measured

(IR imaging; Fig. S1*A*) temperature $T_m$ (g – i). The solid lines in panels (d) and (g) show the model prediction when the droplet temperature is taken to be the cooler than ambient, measured (see fig. S1a**)** $T_m$. The dashed and dash-dotted lines show the model predictions when the droplet temperature is taken to be ambient air temperature, $T_a$, and wet bulb temperature (see eq. $(s20)$), $T_w$, respectively.

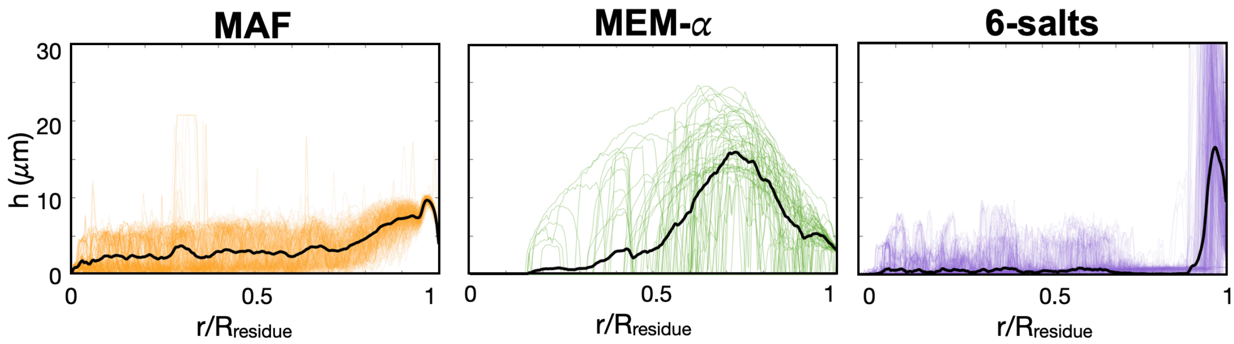


**Fig. S3*E*. Average height profile for residues made of MAF, MEM-$\alpha$, and 6 salts.** Thin lines are radial profiles measured from the center to the (normalized) edge of the residue along 100 evenly spaced angular locations. Thick lines are the averages of the thin lines. For 6-salts the edges show folds (Fig. S3*F*) that were accounted for when estimating the residue height.

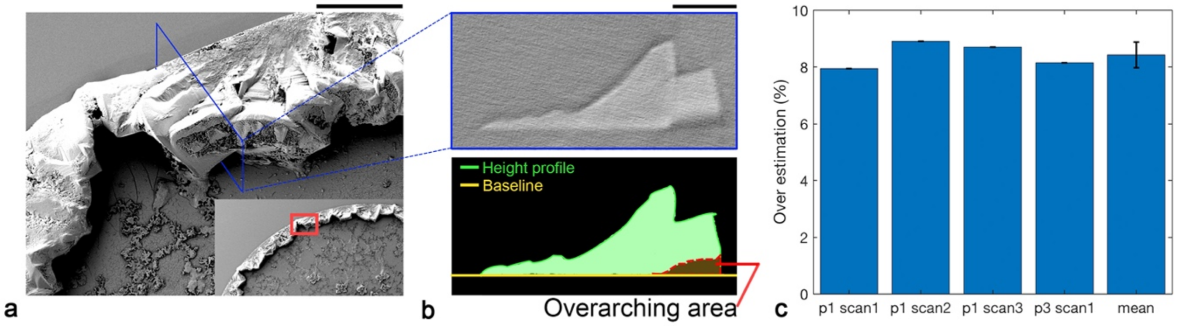


**Fig. S3*F*. Quantification of gap volume below the 6 salts residues.** (a) SEM images of crystals at the elevated edge of the 6 salts+$10^8$/mL BCG residue showing the collapsed buckled layer from an overhang skirt, with a gap underneath. Length scale: 50 $\mu$m. (b) Example of reconstructed cross-section view from micro-CT scan of a 6 salts residue showing the gap below the crystal overhang skirt (top). The height profile overestimates the volume of the crystal by including the gap volume (bottom). Length scale: 25 $\mu$m. (c) The bar plot shows the measurement of such over-estimation following the methods shown in (a) and (b) from different residue images and locations in such images. The average overestimation is around 8% (error bar shows standard deviation).

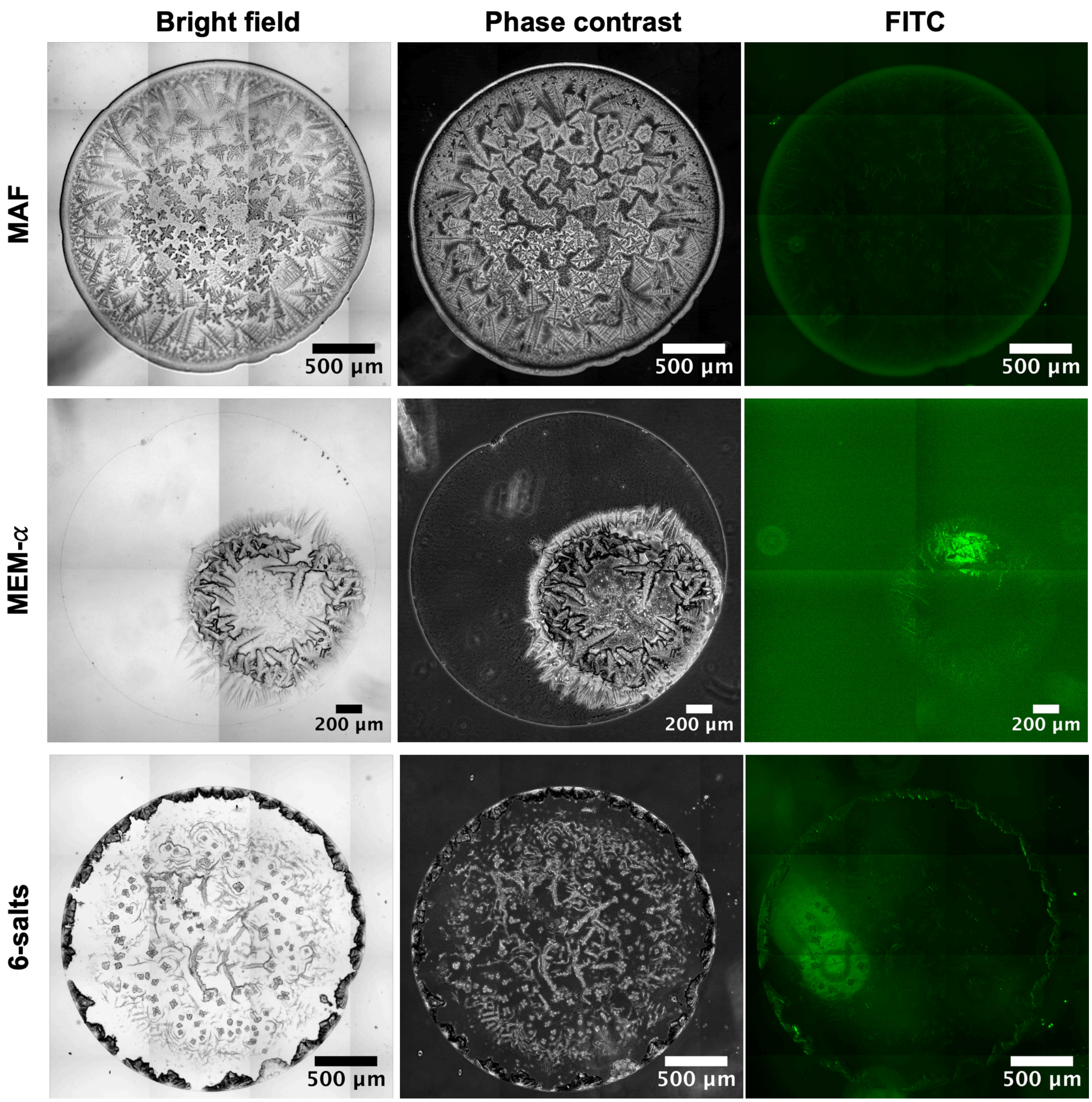


**Fig. S3*G*. Microscopic images of residues without BCG, with bright field (left-most column), phase contrast, (middle) and FITC (right-most).** Here the FITC is only shown as a reference for the fluorescence shown in subsequent figures that contain florescent organisms.

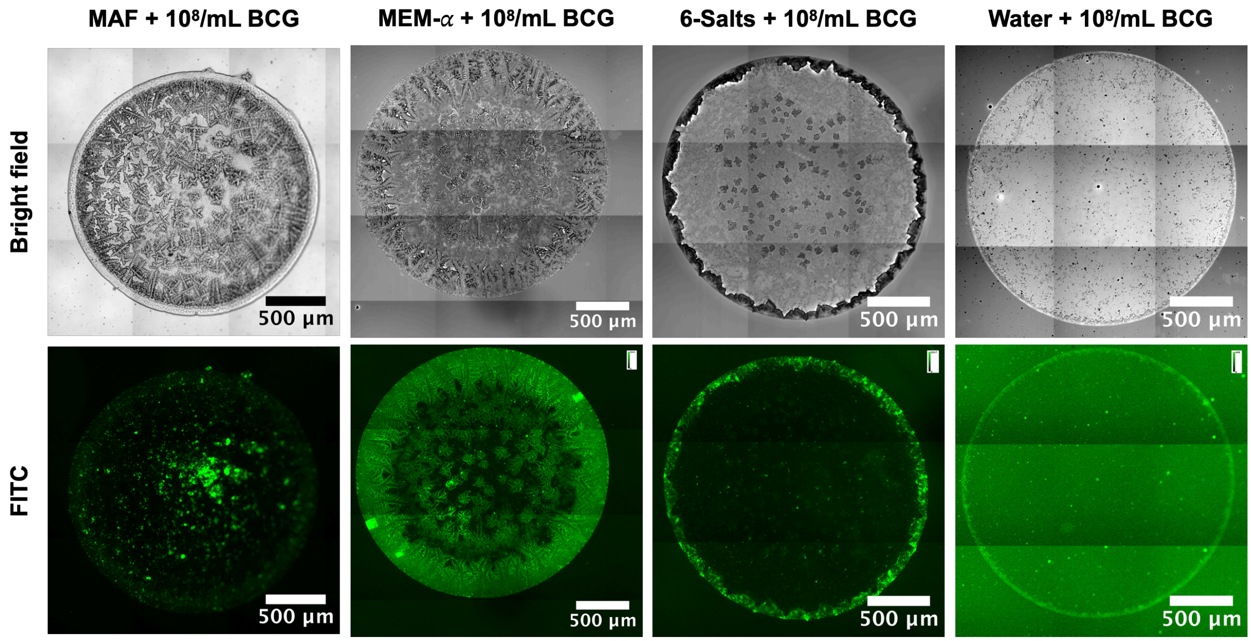


**Fig. *S3H*. Microscopic images of residues with $10^8$ CFU/mL BCG.** All residues indicate that droplets remained pinned during the evaporation, including MEM-$\alpha$. The fluorescent FITC images show that BCG is concentrated at the center of MAF residues while more distributed on the edges in the residues of MEM-$\alpha$, 6 salts, and water.

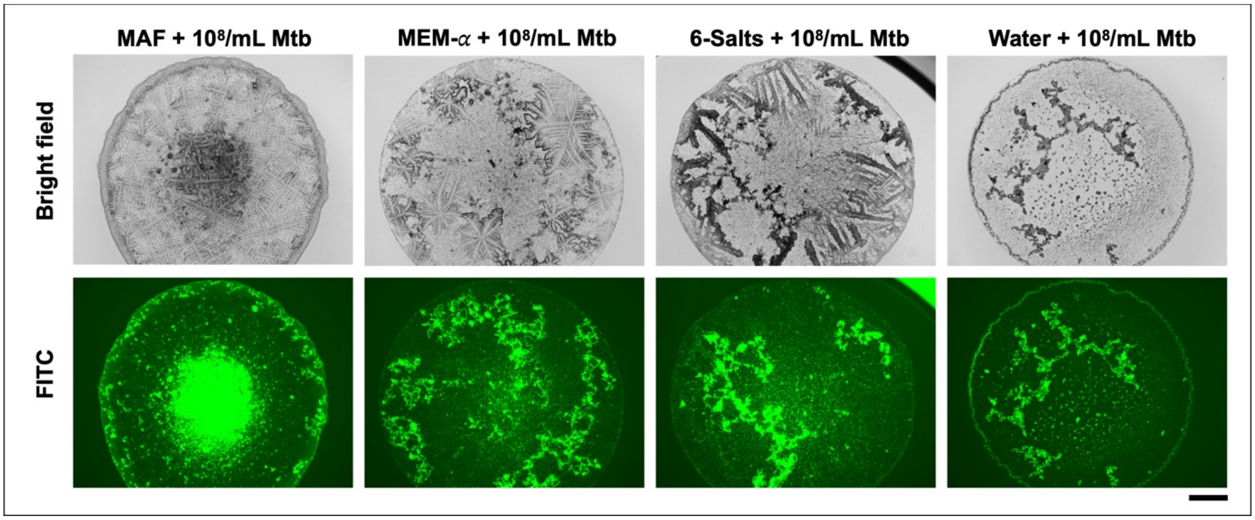


**Fig. S3*I*. Microscopic images of residues with $10^8$ CFU/mL Mtb.** The fluorescent FITC images show that Mtb is concentrated at the center of MAF residues while more distributed in the residues of MEM-$\alpha$, 6 salts, and water and Mtb appears to coat or colocalize with some of the larger-scale dendritic crystals in MEM-$\alpha$ and 6 salts. Scale bar: 500 $\mu$m.

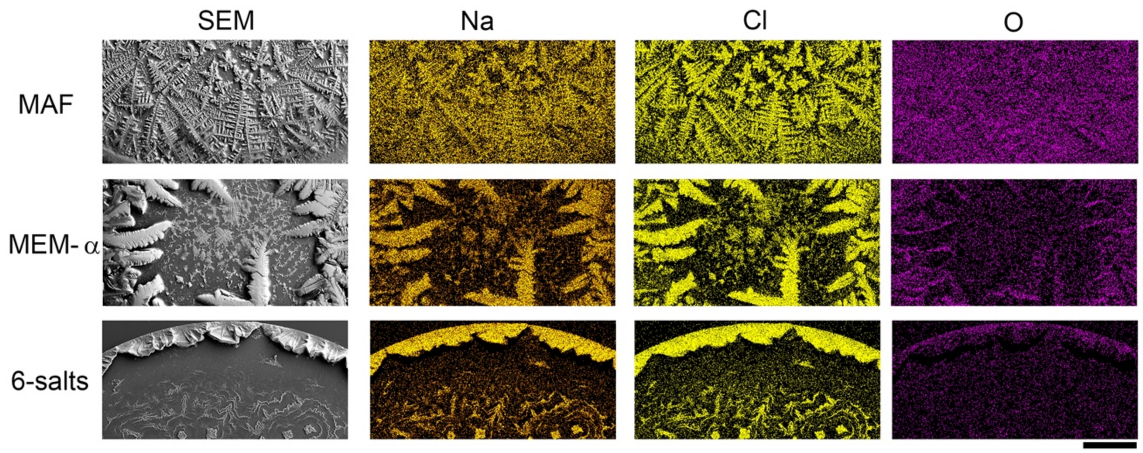


**Fig. S3*J*. EDS analysis of the chemical composition of residues.** Crystals are mainly composed of sodium chloride. In MAF, oxygen appears evenly distributed, whereas in MEM-$\alpha$ and 6 salts, the oxygen appears enriched at the borders of the sharpest crystals. Scale bar: 250 *m*m.

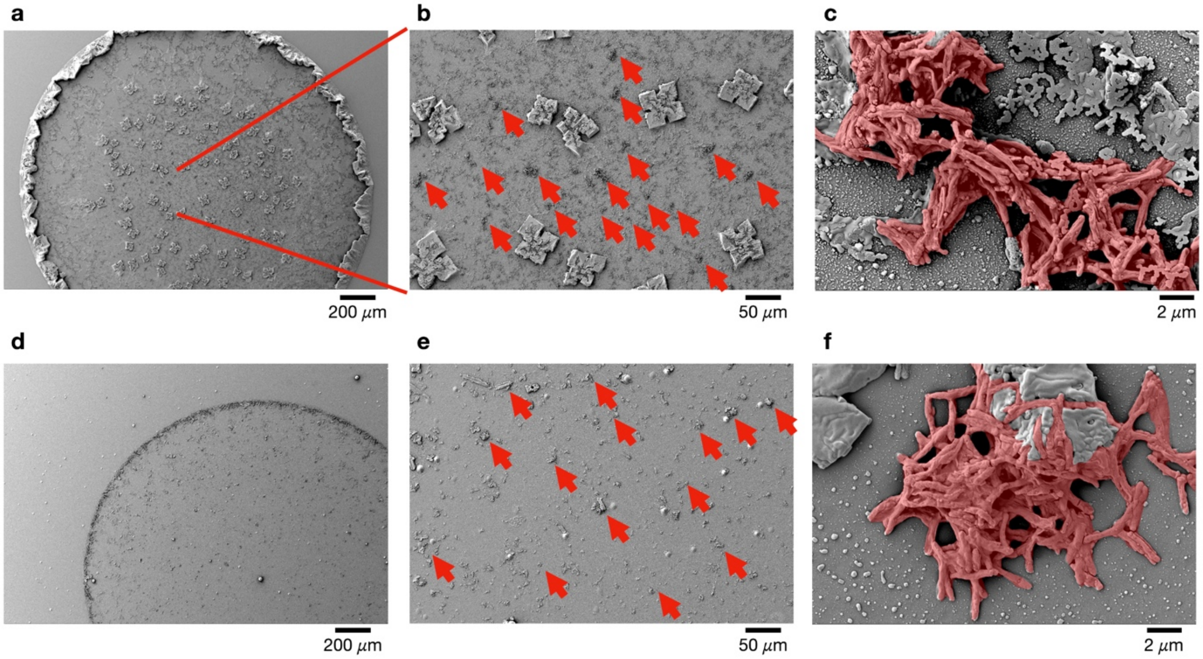


**Fig. S3*K*. SEM images of 6 salts and water with BCG.** SEM images of clustering BCG in residues evaporated from (a - c) 6 salts + $10^8$ CFU/mL BCG and (d - f) water + $10^8$ CFU/mL BCG. Crumpled BCG cells in residues are identified with red arrows.

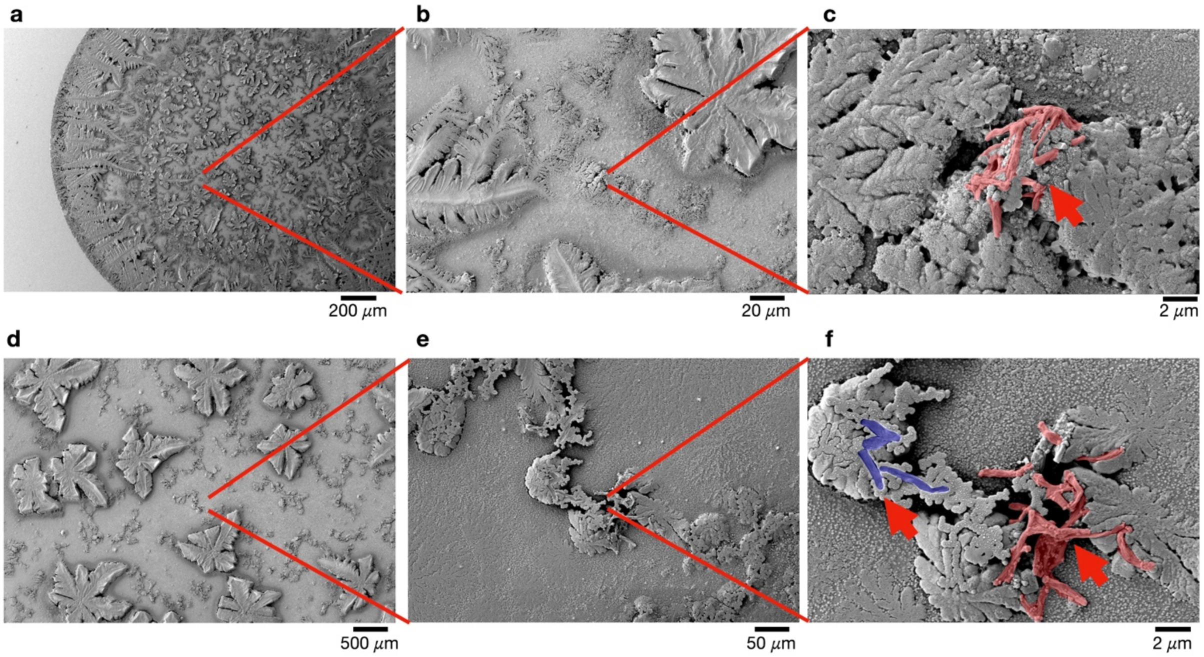


**Fig. S3*L*. SEM images of flattened and crumpled BCG in MEM-$\alpha$ residue identified with red arrows. In subpanel (f), one of the red arrows points to BCG with normal shape, which are false-colored blue.**

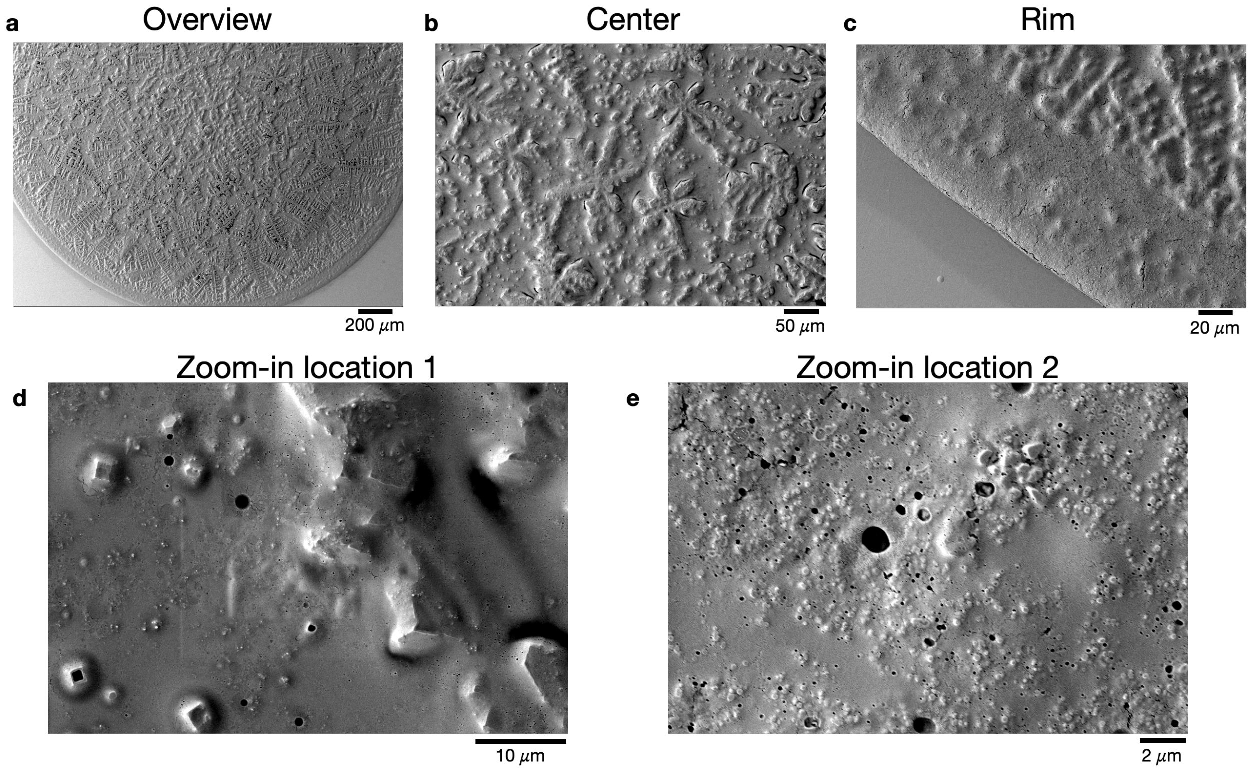


**Fig. S3*M*. SEM images of MAF residue with BCG.** Images show overview and focusing with various magnification levels at the center, rim, and surface a porous surface structure with no visible BCG on the surface, contrary to the BCG in simpler solutions such as 6 salts and MEM-$\alpha$ where the bacilli can be observed on the surface, as seen in Figs. S3*K* and S3*L***.**

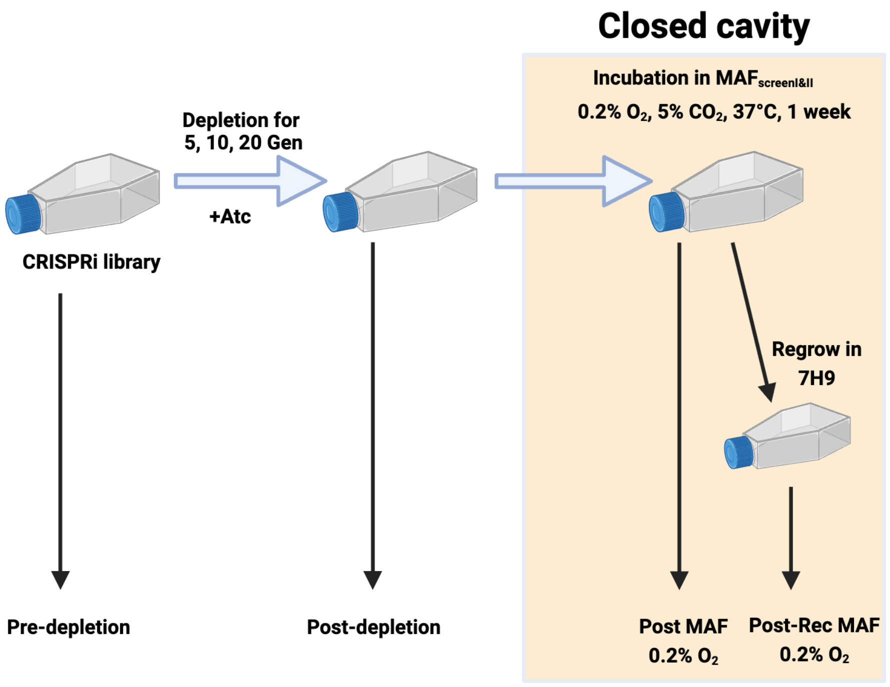


**Fig S4*A*. Schematic of CRISPRi screen I (Created with BioRender.com).**

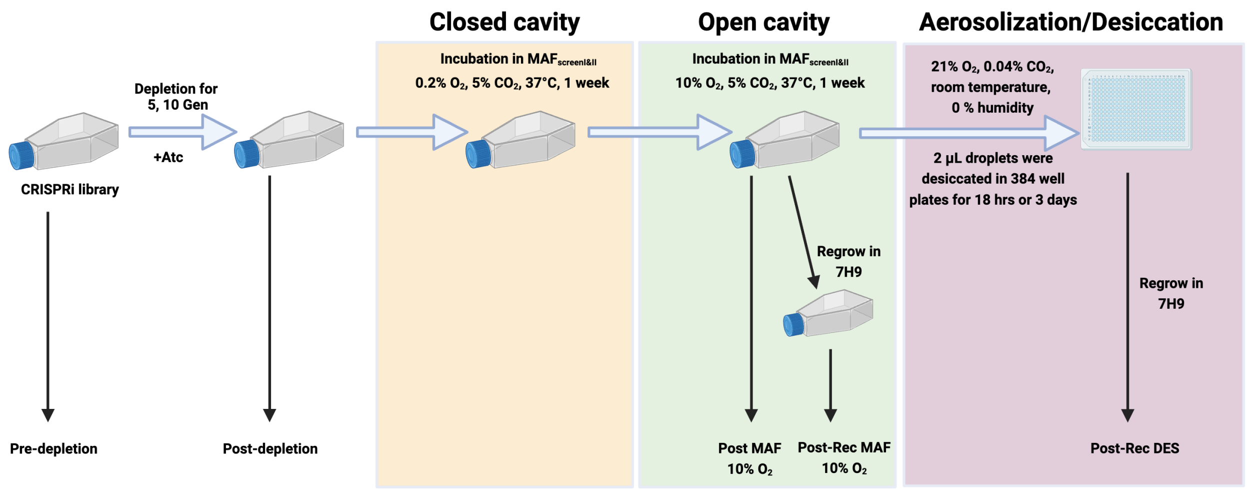


**Fig S4*B*. Schematic of CRISPRi screen II (**Created with BioRender.com).

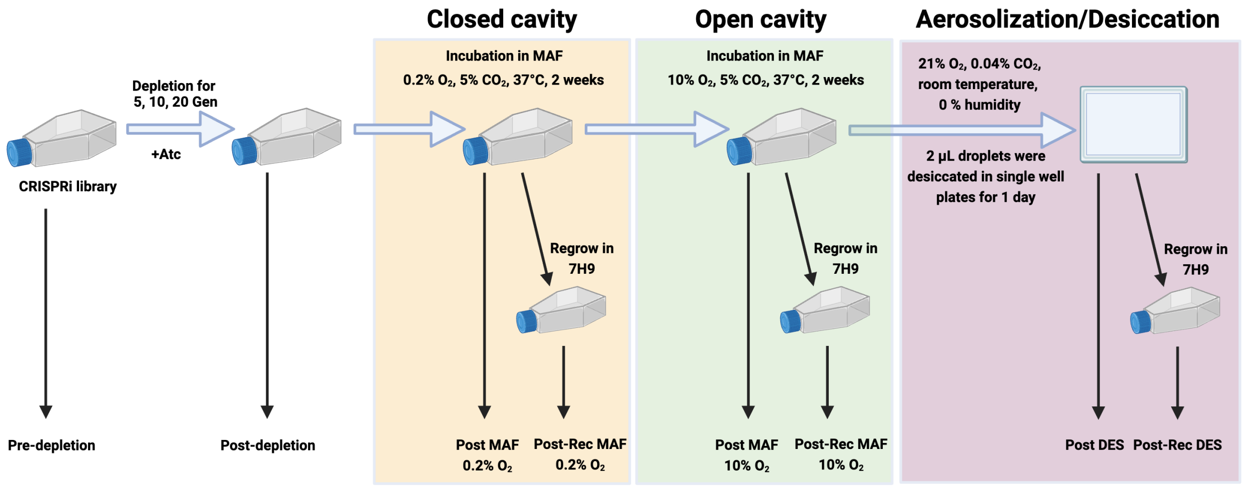


**Fig S4*C*. Schematic of CRISPRi screen III** (Created with BioRender.com).

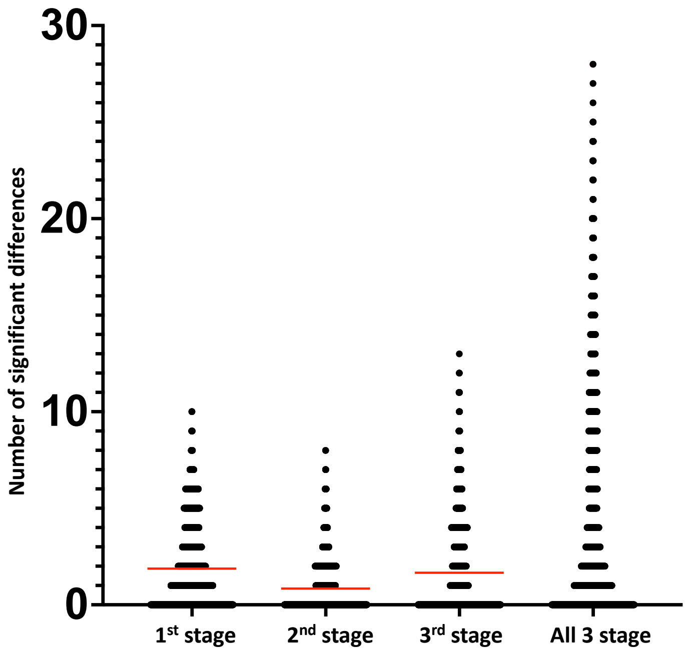


**Fig S4*D*. NOSDs for each gene at each stage of the transmission model and for all three stages.** Each dot represents one of the 4014 genes queried. Horizontal red lines indicate mean values for the stage.

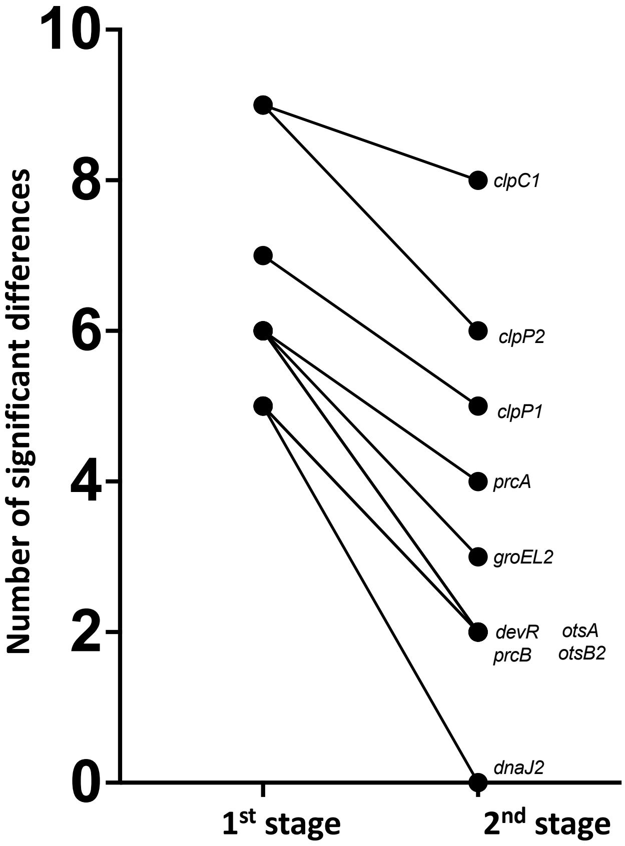


**Fig S4*E*. Changes in NOSDs with re-aeration after hypoxia.**

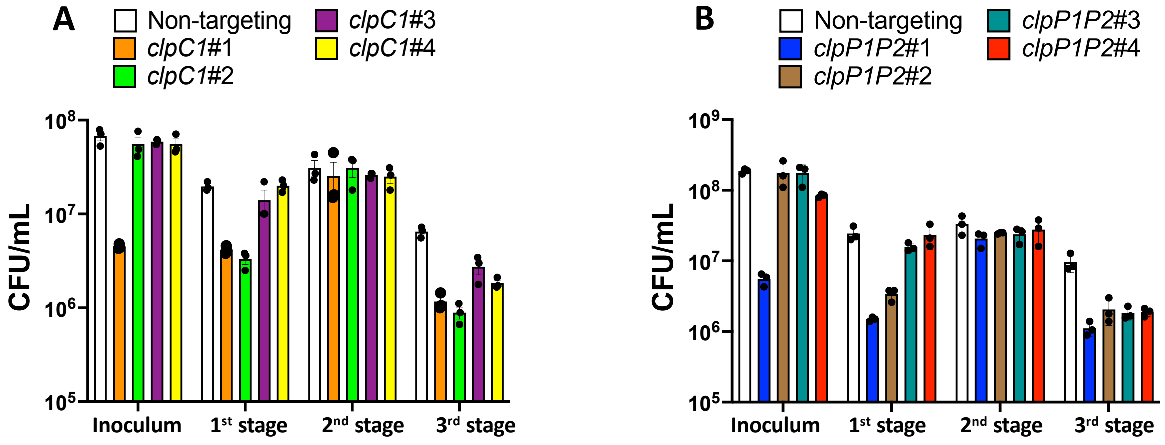

**Fig S5*A*.** Survival assay for Mtb CRISPRi knockdown strains of **(A)** *clpC1* and **(B)** *clpP1P2*. A non-targeting strain (that is, a strain containing an sgRNA directed at nothing in the Mtb genome) was used as control while strains #1 to #4 are hypomorphs with PAM sequences producing a decreasing level of knockdown from #1 to #4.

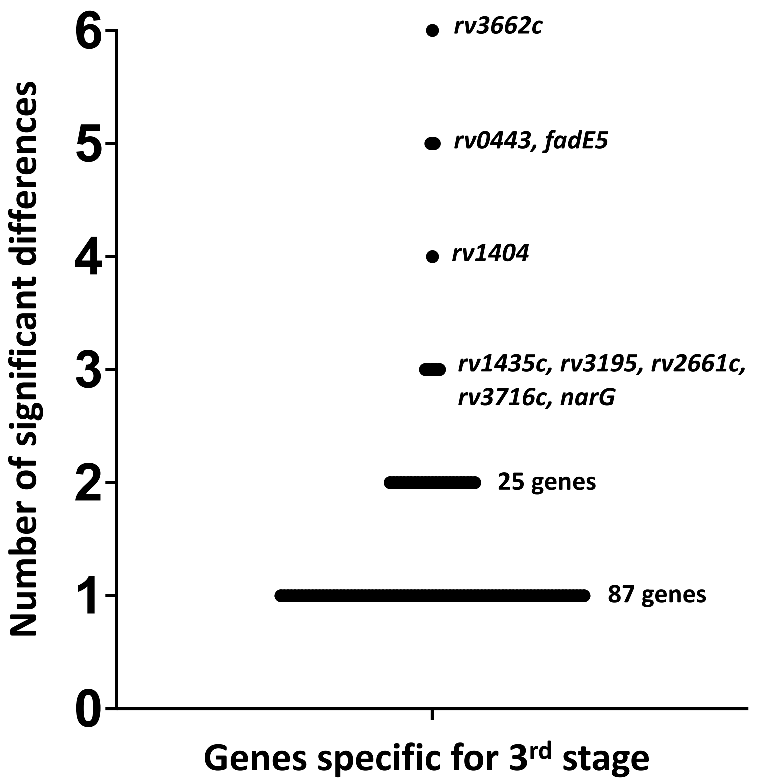


**Fig S5*B*. Genes with NOSDs $\geq$ 1 *only* during the 3rd stage (desiccation).** Each dot represents a gene.

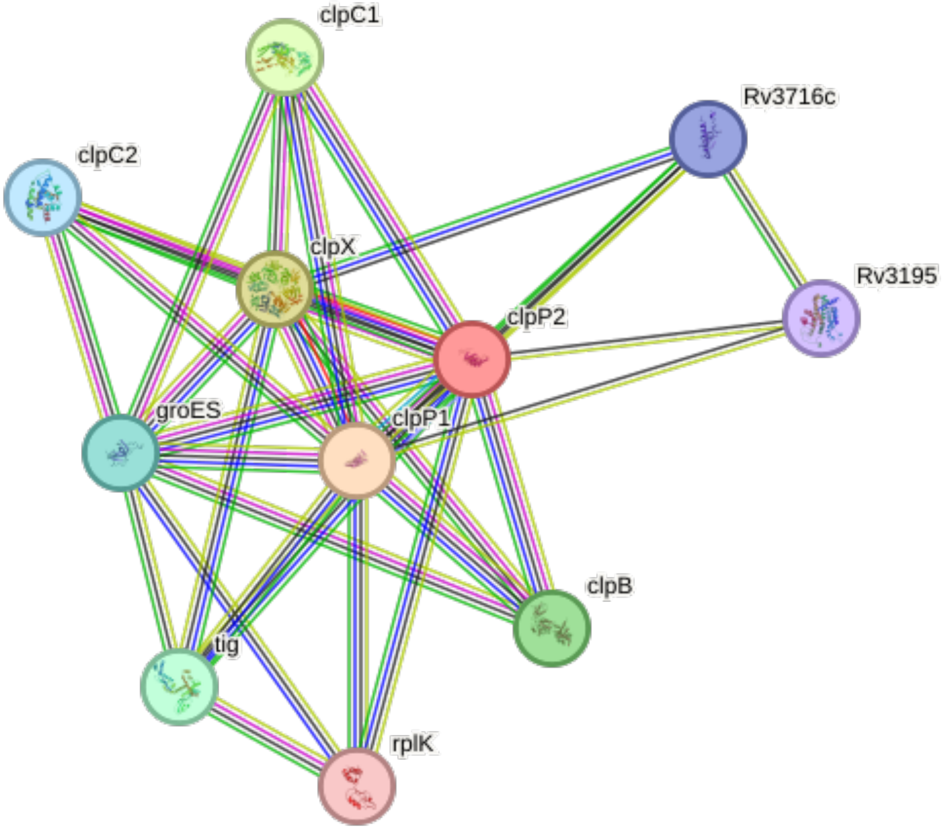


**Fig S5*C*. Bioinformatic links between Clp system, Rv3716c and Rv3195 (https://string-db.org/network/83332.Rv2460c).**

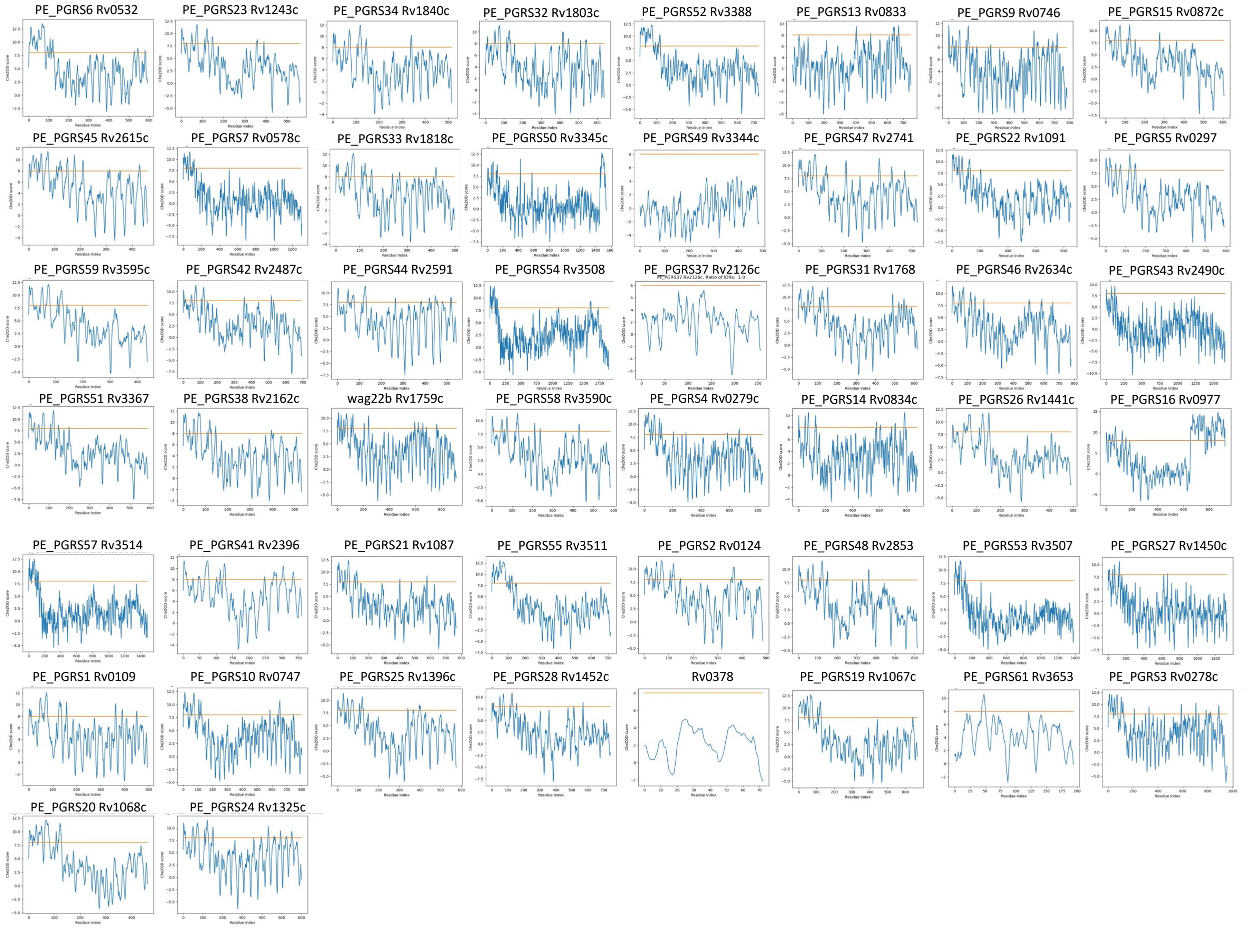


**Fig S6. Intrinsically disordered regions in genes with NOSDs ≥ 1 during 3rd stage (desiccation) as described in ref. S23.** X-Axis: Residue Index, the sequential numbering of amino acid residues within the studied protein, ranging from 0 to the total number of residues. Y-Axis: CheZOD Score, a measure\t of residue disorder within a protein based on NMR data [32696364]. This score is calculated using the SETH model [36304335], which is trained to reproduce experimental features from protein language embeddings derived from the protein's sequence. According to studies in [36304335], a CheZOD Score below 8 (represented by the horizontal orange line) indicates a disordered residue.

**Supplementary tables**

**table S1*A*. Average concentration of metabolites identified in caseum.**

| **Components** | **Molar conc.** | **Components** | **Molar conc.** |
|---|---|---|---|
| | mM | | mM |
| Dimethylgycine | 2.394 | Histidine | 0.200 |
| Adenine | 0.003 | Homoserine | 0.131 |
| (Iso)citric acid | 0.143 | Lactate | 20.060 |
| Citrulline | 0.067 | (Iso)leucine | 0.400 |
| Creatine | 0.266 | Lysine | 0.399 |
| Hexose (Glucose) | 5.556 | Methionine | 0.101 |
| Ethanolamine | 1.647 | Phenylalanine | 0.194 |
| Fumaric acid | 0.079 | Proline | 0.348 |
| Glycine | 0.667 | Serine | 0.238 |
| Guanine | 0.347 | Threonine | 0.403 |
| GABA | 0.040 | Tryptophan | 0.049 |
| Alanine | 0.281 | Valine | 0.393 |
| Arginine | 0.498 | Methionine sulfoxide | 0.004 |
| Asparagine | 0.333 | Ornithine | 0.085 |
| Aspartate | 0.226 | Pyruvic acid | 1.000 |
| Cysteine | 0.568 | Succinic acid | 0.288 |
| Glutamate | 0.510 | Uracil | 0.038 |
| Glutamine | 2.000 | Urea | 8.608 |

**table S1*B*. Concentrations of metabolites identified in 8 caseous necrotic TB lesions.**

| Metabolites. (Red: present in MEM-α; Green: added to MEM-α) | Concentrations in caseum/nodule from 8 human TB lesions (mM) | | | | | | | | Average concentration (mM) |
|---|---|---|---|---|---|---|---|---|---|
| Dimethylgycine | 2.44636 | 0.0011 | 0.0011 | 0.0011 | 4.790713 | 9.551637 | 2.360256 | 0.0011 | 2.394 |
| Adenine | 0.0011 | 0.008372 | 0.007151 | 0.0011 | 0.0011 | 0.0011 | 0.0011 | 0.0011 | 0.003 |
| (Iso)citric acid | 0.119806 | 0.017614 | 0.006111 | 0.072007 | 0.253189 | 0.264696 | 0.336832 | 0.076408 | 0.143 |
| Citrulline | 0.0011 | 0.01303 | 0.059774 | 0.056601 | 0.163523 | 0.130575 | 0.097505 | 0.013704 | 0.067 |
| Creatine | 0.581086 | 0.240248 | 0.160877 | 0.071155 | 0.144096 | 0.422527 | 0.395206 | 0.114969 | 0.266 |
| Hexose (Glucose) | 4.905952 | 1.966355 | 0.622116 | 0.368965 | 0.763197 | 3.289337 | 1.867553 | 0.880926 | 5.556 |
| Ethanolamine | 2.519237 | 5.378674 | 0.0011 | 0.0011 | 0.391447 | 0.156966 | 0.397555 | 4.329536 | 1.647 |
| Fumaric acid | 0.031722 | 0.091692 | 0.011711 | 0.019289 | 0.018588 | 0.185235 | 0.224545 | 0.050369 | 0.079 |
| Glycine | 1.268718 | 0.539828 | 0.613291 | 0.645056 | 1.538489 | 2.773081 | 1.909503 | 0.527252 | 0.667 |
| Guanine | 0.246784 | 0.246694 | 0.260154 | 0.217576 | 0.387153 | 0.500995 | 0.511808 | 0.406834 | 0.347 |
| GABA | 0.052897 | 0.030039 | 0.060408 | 0.020857 | 0.021474 | 0.065527 | 0.046205 | 0.020595 | 0.040 |
| Alanine | 1.948556 | 1.008586 | 2.072184 | 1.274689 | 2.866751 | 3.919854 | 3.198456 | 0.769938 | 0.281 |
| Arginine | 0.075456 | 0.050296 | 0.214644 | 0.097045 | 0.236896 | 0.278266 | 0.365245 | 0.077169 | 0.498 |
| Asparagine | 0.371065 | 0.238097 | 0.35312 | 0.220373 | 0.689721 | 0.749916 | 0.907746 | 0.302951 | 0.333 |
| Aspartate | 0.481133 | 0.132365 | 0.153612 | 0.235398 | 0.600942 | 1.177926 | 0.397526 | 0.211044 | 0.226 |
| Cysteine | 0.061045 | 0.0011 | 0.0011 | 0.0011 | 0.0011 | 0.162841 | 0.117734 | 0.0011 | 0.568 |
| Glutamate | 0.508413 | 0.194273 | 0.150746 | 0.283901 | 0.843178 | 1.652884 | 0.478009 | 0.164019 | 0.510 |
| Glutamine | 1.744896 | 0.792994 | 0.763114 | 0.082842 | 0.105881 | 0.685901 | 1.412436 | 0.770622 | 2.000 |
| Histidine | 0.094149 | 0.062346 | 0.19614 | 0.116401 | 0.284911 | 0.274839 | 0.320991 | 0.078761 | 0.200 |
| Homoserine | 0.043709 | 0.013255 | 0.064454 | 0.082795 | 0.263648 | 0.349887 | 0.204495 | 0.02669 | 0.131 |
| Lactate | 20.88617 | 11.07947 | 38.42265 | 13.5225 | 23.11153 | 21.89736 | 28.07979 | 3.477581 | 20.060 |
| (Iso)leucine | 0.701579 | 0.349128 | 0.683037 | 0.670002 | 1.626739 | 0.962272 | 0.885223 | 0.384819 | 0.400 |
| Lysine | 0.107438 | 0.075992 | 0.336512 | 0.383298 | 0.890639 | 1.011339 | 0.627775 | 0.217756 | 0.399 |
| Methionine | 0.083644 | 0.0011 | 0.098898 | 0.06222 | 0.231872 | 0.343552 | 0.31296 | 0.040619 | 0.101 |
| Phenylalanine | 0.064248 | 0.051117 | 0.111999 | 0.056096 | 0.207786 | 0.247942 | 0.344613 | 0.077365 | 0.194 |
| Proline | 0.419873 | 0.204058 | 0.465492 | 0.300821 | 0.689554 | 1.107512 | 0.919626 | 0.143911 | 0.348 |
| Serine | 0.262764 | 0.225806 | 0.297026 | 0.437725 | 0.947524 | 1.196121 | 0.815621 | 0.288702 | 0.238 |
| Threonine | 0.252362 | 0.178062 | 0.329424 | 0.436532 | 1.112955 | 1.367035 | 0.914194 | 0.253389 | 0.403 |
| Tryptophan | 0.006305 | 0.008123 | 0.016909 | 0.014848 | 0.039292 | 0.054201 | 0.068823 | 0.016064 | 0.049 |
| Valine | 0.417654 | 0.28286 | 0.750929 | 0.570107 | 1.20866 | 1.324725 | 1.444898 | 0.279219 | 0.393 |
| Methionine sulfoxide | 0.0011 | 0.0011 | 0.0011 | 0.0011 | 0.016006 | 0.0011 | 0.00842 | 0.0011 | 0.004 |
| Ornithine | 0.0011 | 0.0011 | 0.071348 | 0.074604 | 0.257551 | 0.122469 | 0.154045 | 0.0011 | 0.085 |
| Pyruvic acid | 0.0011 | 0.031297 | 0.300708 | 0.029525 | 0.036489 | 0.651577 | 0.38742 | 0.027994 | 1.000 |
| Succinic acid | 0.246565 | 0.158383 | 0.215386 | 0.205875 | 0.41212 | 0.32478 | 0.611193 | 0.126781 | 0.288 |
| Uracil | 0.008831 | 0.006026 | 0.006881 | 0.0011 | 0.0011 | 0.276825 | 0.0011 | 0.0011 | 0.038 |
| Urea | 4.956289 | 2.374756 | 11.64219 | 9.038622 | 12.84214 | 12.06038 | 13.67338 | 2.275759 | 8.608 |

**table S1*C*: Comparison of MAFscreen$_{I\&II}$ and MAF compositions.**

| Recipe for preparation of MAFscreen$_{I\&II}$ and MAF | | |
|---|---|---|
| | **MAFscreen$_{I\&II}$** | **MAF** |
| Lipid and phospholipid mix (prepared in 100% Ethanol) | 3% v/v | 3% v/v |
| Serum | 3% v/v | 3% v/v |
| PMN lysate | 0.17 µg/mL DNA (≃1% v/v) | 0.17 µg/µL DNA (≃1% v/v) |
| Deoxysphinganine | 70 ng/mL | 70 ng/mL |
| MEM-α + additional metabolites | 93% v/v | 93% v/v |
| **Additional metabolites added to MEM-α** | | |
| | **For MAFscreen$_{I\&II}$** | **For MAF** |
| Urea | 8.60 mM | 8.60 mM |
| Dimethylglycine | 2.39 mM | 2.39 mM |
| Lactate | 20.06 mM | 20.06 mM |
| Iso-citric acid | 143 µM | 143 µM |
| Citrulline | 67 µM | 67 µM |
| Creatine | 266 µM | 266 µM |
| Ethanolamine | 1.65 mM | 1.65 mM |
| Fumaric acid | 79 µM | 79 µM |
| GABA | 40 µM | 40 µM |
| Homoserine | 131 µM | 131 µM |
| Methionine sulfoxide | 4 µM | 4 µM |
| Ornithine | 85 µM | 85 µM |
| Succinic acid | 288 µM | 288 µM |
| Adenine | Not added | 3 µM |
| Uracil | Not added | 38 µM |
| Guanine | Not added | 347 µM |
| **Lipids and phospholipids added to lipid and phospholipid mix** | | |
| | **For MAFscreen$_{I\&II}$** | **For MAF** |
| DPPG | 0.05 mg/mL | 0.05 mg/mL |
| DPPC | 0.5 mg/mL | 0.5 mg/mL |
| Sphingomyelin | 0.01 mg/mL | 0.03 mg/mL |
| Glyceryl trioleate | 0.02 mg/mL | 0.15 mg/mL |
| 1,3-dioleoyl-2-palmitoylglycerol | 0.02 mg/mL | 0.15 mg/mL |
| Cholesterol | 0.007 mg/mL | 0.075 mg/mL |
| Cholesteryl oleate | 0.005 mg/mL | 0.018 mg/mL |
| Cholesteryl linoleate | Not added | 0.018 mg/mL |
| C24:1 Lactosyl(ß) Ceramide (d18:1/24:1) | Not added | 0.005 mg/mL |
| C16 Lactosylceramide (d18:1/16:0) | Not added | 0.005 mg/mL |

**table S2*A*. Composition of MAF.**

| No. | Subgroups of media | Amount | Volume fraction of non-volatile components $\phi$ |
|---|---|---|---|
| | | | ppm |
| 1 | MEM-α[a] + additional metabolites excluding guanine[b] | 93% (v/v) | 5858.85 + 2405.97 = 8264.82 |
| 2 | Guanine[b] | Added powder directly to MAF | 35.70 |
| 3 | Serum[c] | 3% (v/v) | 2126.71 |
| 4 | Lipid and phospholipid mix including deoxysphinganine (prepared in 100% Ethanol)[d] | 3% (v/v) | 992.96 |
| 5 | PMN lysate[e] (0.17 μg DNA/μL MAF) | 1% (v/v) | 176.84 |
| | **Sum** | | 11,597.02 |

[a]: See table S2b for the detailed composition of MEM-*a*
[b]: See table S2c for the detailed composition of additional metabolites added to MEM-*a*
[c]: See table S2d for the detailed composition of serum.
[d]: See table S2e for the detailed composition of the lipid and phospholipid mix.
[e]: See table S2f for the detailed composition of PMN lysate.

**table S2*B*. Composition of MEM-*a* aqueous base.**

| Components | Molecular Weight | Molar conc. | Mass conc. $c_m$ | Density $\rho$ | Volume fraction $\phi$ |
|---|---|---|---|---|---|
| | g/mol | mM | mg/L | g/cm³ | ppm |
| ***Amino Acid*** | | | | | |
| Glycine | 75.07 | 0.67 | 50 | 1.16 [S13] | 43.10 |
| L-Alanine | 89.09 | 0.28 | 25 | 1.43[S13] | 17.48 |
| L-Arginine hydrochloride | 210.66 | 0.50 | 105 | 1.42 [S24] | 73.94 |
| L-Asparagine-H2O | 150.13 | 0.33 | 50 | 1.54 [S13] | 32.47 |
| L-Aspartic acid | 133.10 | 0.23 | 30 | 1.66 [S13] | 18.07 |
| L-Cysteine hydrochloride-H2O | 175.60 | 0.57 | 100 | 1.54 [S25] | 64.94 |

| | | | | | |
|---|---|---|---|---|---|
| L-Cystine 2HCl | 313.22 | 0.099 | 31 | 1.52 [S26] | 20.39 |
| L-Glutamic Acid | 147.13 | 0.51 | 75 | 1.54 [S13] | 48.70 |
| L-Glutamine | 146.14 | 2.00 | 292 | 1.18 [S27] | 247.46 |
| L-Histidine | 155.15 | 0.20 | 31 | 1.24 [S27] | 25.00 |
| L-Isoleucine | 131.17 | 0.40 | 52.4 | 1.00 [S27] | 52.40 |
| L-Leucine | 131.17 | 0.40 | 52 | 1.29 [S13] | 40.31 |
| L-Lysine hydrochloride | 182.65 | 0.40 | 73 | 1.28 [S28] | 57.03 |
| L-Methionine | 149.21 | 0.10 | 15 | 1.11 [S27] | 13.51 |
| L-Phenylalanine | 165.19 | 0.20 | 32 | 1.28 [S27] | 25.00 |
| L-Proline | 115.13 | 0.35 | 40 | 1.06 [S27] | 37.74 |
| L-Serine | 105.09 | 0.24 | 25 | 1.60 [S27] | 15.63 |
| L-Threonine | 119.12 | 0.40 | 48 | 1.08 [S27] | 44.44 |
| L-Tryptophan | 204.23 | 0.050 | 10 | 1.19 [S27] | 8.40 |
| L-Tyrosine disodium salt | 225.15 | 0.23 | 52 | 1.40 [S29] | 37.14 |
| L-Valine | 117.15 | 0.39 | 46 | 0.93 [S27] | 49.46 |
| ***Vitamins*** | | | | | |
| Ascorbic Acid | 176.12 | 0.28 | 50 | 1.65 [S27] | 30.30 |
| Biotin | 244.31 | 4.10E-04 | 0.1 | 1.20 [S27] | 0.08 |
| Choline chloride | 139.62 | 0.007 | 1 | 0.97 [S27] | 1.03 |
| D-Calcium pantothenate | 476.53 | 0.0021 | 1 | 1.27 [S30] | 0.79 |
| Folic Acid | 441.40 | 0.0023 | 1 | 1.44 [S27] | 0.69 |
| Niacinamide | 122.12 | 0.0082 | 1 | 1.40 [S13] | 0.71 |
| Pyridoxal hydrochloride | 203.62 | 0.0049 | 1 | 1.28 [S31] | 0.78 |
| Riboflavin | 376.36 | 2.66E-04 | 0.1 | 1.33 [S27] | 0.08 |
| Thiamine hydrochloride | 337.26 | 0.003 | 1 | 1.30 [S27] | 0.77 |
| Vitamin B12 | 1355.38 | 0.001 | 1.36 | 0.95 [S32] | 1.43 |
| i-Inositol | 180.16 | 0.01 | 2 | 1.75 [S27] | 1.14 |
| ***Inorganic salts and bicarbonate*** | | | | | |
| Calcium Chloride (CaCl2) (anhyd.) | 110.98 | 1.80 | 200 | 2.15 [S13] | 93.02 |
| Magnesium Sulfate (MgSO4) (anhyd.) | 120.37 | 0.81 | 97.67 | 2.66 [S13] | 36.72 |
| Potassium Chloride (KCl) | 74.55 | 5.33 | 400 | 1.99 [S13] | 201.01 |
| Sodium Bicarbonate (NaHCO3) | 84.01 | 26.19 | 2200 | 2.20 [S13] | 1000.00 |

| | | | | | |
|---|---|---|---|---|---|
| Sodium Chloride (NaCl) | 58.44 | 117.24 | 6800 | 2.17 [S13] | 3133.64 |
| Sodium Phosphate monobasic (NaH2PO4-H2O) | 137.99 | 1.01 | 140 | 2.04 [S33] | 68.63 |
| ***Carbon source*** | | | | | |
| D-Glucose (Dextrose) | 180.16 | 5.56 | 1000 | 1.56 [S27] | 641.03 |
| ***Other organics*** | | | | | |
| Adenosine | 267.24 | 0.037 | 10 | 1.39 [S27] | 7.19 |
| Cytidine | 243.22 | 0.041 | 10 | 1.33 [S27] | 7.52 |
| Guanosine | 283.24 | 0.035 | 10 | 1.43 [S27] | 6.99 |
| Uridine | 244.20 | 0.041 | 10 | 1.36 [S27] | 7.35 |
| 2'-Deoxyadenosine | 251.24 | 0.040 | 10 | 1.90 [S34] | 5.26 |
| 2'-Deoxycytidine HCl | 263.68 | 0.042 | 11 | 1.70 [S35] | 6.47 |
| 2'-Deoxyguanosine | 267.24 | 0.037 | 10 | 2.10 [S36] | 4.76 |
| Thymidine | 242.23 | 0.041 | 10 | 1.27 [S27] | 7.87 |
| Lipoic Acid | 180.16 | 9.7E-04 | 0.2 | 1.20 [S37] | 0.17 |
| Sodium Pyruvate | 110.04 | 1 | 110 | 1.78 [S38] | 61.80 |
| **Sum** | | | | | **6299.84** |
| **Final volume fraction in MAF** | | | | | **6299.84*93% = 5858.85** |

**table S2*C*. Composition of the additional metabolites found in caseum.**

| Components | Molecular Weight $m_w$ | Molar conc. $c_n$ (Added in MEM) | Mass conc. $c_m$ (Added in MEM) | Density $\rho$ | Volume fraction $\phi$ |
|---|---|---|---|---|---|
| | g/mol | mM | mg/L | $g/cm^3$ | ppm |
| N,N-Dimethylglycine hydrochloride | 139.58 | 2.394 | 334.18 | 1.33 [S39] | 251.26 |
| Adenine | 135.13 | 0.003 | 0.37 | 1.39 [S27] | 0.27 |
| DL-Isocitric acid trisodium salt hydrate | 258.07 | 0.143 | 36.99 | 1.75 [S40] | 21.14 |
| Citrulline | 175.19 | 0.067 | 11.734 | 1.18 [S27] | 9.94 |
| Creatine | 131.13 | 0.266 | 34.916 | 1.33 [S27] | 26.25 |
| Ethanolamine hydrochloride | 97.54 | 1.647 | 160.64 | 1.12 [S41] | 143.43 |
| Fumaric acid | 116.07 | 0.079 | 9.19 | 1.64 [S27] | 5.60 |
| GABA | 103.12 | 0.040 | 4.10 | 1.10 [S42] | 3.73 |
| Homoserine | 119.12 | 0.131 | 15.62 | 1.08 [S27] | 14.46 |
| Sodium L-lactate | 112.06 | 20.060 | 2247.88 | 1.33 [S43] | 1690.14 |
| L-Methionine sulfoxide | 165.21 | 0.004 | 0.64 | 1.18 [S27] | 0.54 |
| L-Ornithine monohydrochloride | 168.62 | 0.085 | 14.40 | 1.43 [S44] | 10.07 |
| Succinic acid | 118.09 | 0.288 | 33.97 | 1.57 [S27] | 21.63 |
| Uracil | 112.09 | 0.038 | 4.24 | 1.34 [S27] | 3.17 |
| Urea | 60.00 | 8.608 | 516.476 | 1.34 [S27] | 385.43 |
| **Sum** | | | | | **2587.06** |
| **Final density in MAF** | | | | | **2587.06*93%<br>2405.97** |
| **Components** | **Molecular Weight $m_w$** | **Molar conc. $c_n$ (Added in MAF)** | **Mass conc. $c_m$ (Added in MAF)** | **Density $\rho$** | **Volume fraction $\phi$** |
| Guanine | 151.13 | 0.347 | 52.480 | 1.47 [S27] | 35.70 |

**table S2*D*. Composition of serum and its final concentration in MAF.**

| Components | Mass conc. $c_m$ | Density $\rho$ | Volume fraction $\phi$ |
|---|---|---|---|
| | mg/L | g/cm³ | ppm |
| Proteins | 2169.24 | 1.02 [S45,S46] | 2126.71 |

**table S2E. Composition of the lipid and phospholipid mix and their final concentration in MAF.**

| Components | Molecular Weight $m_w$ | Molar conc. $c_n$ | Mass conc. $c_m$ | Density $\rho$ | Volume fraction $\phi$ |
|---|---|---|---|---|---|
| | g/mol | mM | mg/L | g/cm³ | ppm |
| DPPG (1,2-Dipalmitoyl-sn-glycero-3-phospho-rac-(1-glycerol) ammonium salt) | 740.00 | 0.069 | 50 | 1 | 50.00 |
| DPPC | 734.09 | 0.681 | 500 | 1.08 [S47] | 462.96 |
| sphingomyelin | 813.20 | 0.037 | 30 | 1 | 30.00 |
| glyceryl trioleate | 885.43 | 0.169 | 150 | 0.9 [S48] | 166.67 |
| 1,3-dioleoyl-2-palmitoylglycerol | 859.39 | 0.175 | 150 | 0.9 [S49] | 166.67 |
| cholesterol | 386.65 | 0.194 | 75 | 1.07 [S27] | 70.09 |
| cholesteryl oleate | 651.10 | 0.028 | 18 | 0.91 [S27] | 19.78 |
| cholesteryl linoleate | 649.08 | 0.028 | 18 | 1 [S50] | 18.00 |
| C24:1 Lactosyl(ß) Ceramide (d18:1/24:1) | 972.38 | 0.0051 | 5 | 1.1 [S51] | 4.55 |
| C16 Lactosylceramide (d18:1/16:0) | 862.20 | 0.0058 | 5 | 1.2 [S52] | 4.17 |
| Deoxysphinganine | 285.51 | 0.00025 | 0.07 | 0.9 [S53] | 0.078 |
| **Sum** | | | | | 992.96 |

The densities of DPPG (1,2-Dipalmitoyl-sn-glycero-3-phospho-rac-(1-glycerol) ammonium salt) and sphingomyelin were assumed to be 1.

**table S2*F*. Composition of PMN lysate and their final concentration in MAF.**

| Components | Mass conc. $c_m$ | Density $\rho$ | Volume fraction $\phi$ |
|---|---|---|---|
| | mg/L | g/cm$^3$ | ppm |
| Protein | 78.97 | 1.02 [S45,S46] | 77.42 |
| DNA | 170 | 1.71 [S54] | 99.42 |
| **Sum** | | | **176.84** |

**table S2G. Composition of 6-salts in MEM-*α*.**

| Components | Molecular Weight | Molar conc. | Mass conc. $c_m$ | Density $\rho$ | Volume fraction $\phi$ |
|---|---|---|---|---|---|
| | g/mol | mM | mg/L | g/cm$^3$ | ppm |
| Calcium Chloride (CaCl2) (anhyd.) | 110.98 | 1.80 | 200 | 2.15 [S13] | 93.02 |
| Magnesium Sulfate (MgSO4) (anhyd.) | 120.37 | 0.81 | 97.67 | 2.66 [S13] | 36.72 |
| Potassium Chloride (KCl) | 74.55 | 5.33 | 400 | 1.99 [S13] | 201.01 |
| Sodium Bicarbonate (NaHCO3) | 84.01 | 26.19 | 2200 | 2.20 [S13] | 1000.00 |
| Sodium Chloride (NaCl) | 58.44 | 117.24 | 6800 | 2.17 [S13] | 3133.64 |
| Sodium Phosphate monobasic (NaH2PO4-H2O) | 137.99 | 1.01 | 140 | 2.04 [S33] | 68.63 |
| **Sum** | | | | | **4533.02** |

**table S2*H*. Non-volatile component volume fraction of media.**

| Media | Non-volatile component volume fraction |
|---|---|
| | ppm |
| MAF | 11,597.02 |
| Aqueous base (MEM-$\alpha$) | 6299.84 |
| 6-salts | **4533.02** |

**table S3. Mean shear viscosity within the confident region for the investigated fluids.**

| Fluid | Mean viscosity $(mPa \cdot s)$ | Standard deviation $(mPa \cdot s)$ |
|---|---|---|
| Water | 0.98 | 0.02 |
| 6-salts | 1.00 | 0.02 |
| MEM-$\alpha$ | 1.00 | 0.02 |
| MAF | 1.18 | 0.04 |
| MAF + $10^8$/mL BCG | 1.15 | 0.01 |

**table S4. Primers used for construction of CRISPRi knockdown strains.**

| Primer name | Primer sequence: 5'-3' |
|---|---|
| dnaJ2CRISPRifor | GGGAGCCCCCACCGAAACCCCCGCC |
| dnaJ2CRISPRirev | AAACGGCGGGGGTTTCGGTGGGGGC |
| groEL2CRISPRifor | GGGATTCGCTGATCACCTGACCACCGG |
| groEL2CRISPRirev | AAACCCGGTGGTCAGGTGATCAGCGAA |
| devRCRISPRifor | GGGAGCACCACCTCGTGGTCATCGAC |
| devRCRISPRirev | AAACGTCGATGACCACGAGGTGGTGC |
| otsACRISPRifor | GGGAACAGCGGCCACAGTGTGGCGTT |
| otsACRISPRirev | AAACAACGCCACACTGTGGCCGCTGT |
| otsB2CRISPRifor | GGGAGTCGCCGAGGTAGATCGGCACC |
| otsB2CRISPRirev | AAACGGTGCCGATCTACCTCGGCGAC |

S2. Park Y, Pacitto A, Bayliss T, Cleghorn LA, Wang Z, Hartman T, Arora K, Ioerger TR, Sacchettini J, Rizzi M, Donini S, Blundell TL, Ascher DB, Rhee K, Breda A, Zhou N, Dartois V, Jonnala SR, Via LE, Mizrahi V, Epemolu O, Stojanovski L, Simeons F, Osuna-Cabello M, Ellis L, MacKenzie CJ, Smith AR, Davis SH, Murugesan D, Buchanan KI, Turner PA, Huggett M, Zuccotto F,

Rebollo-Lopez MJ, Lafuente-Monasterio MJ, Sanz O, Diaz GS, Lelievre J, Ballell L, Selenski C, Axtman M, Ghidelli-Disse S, Pflaumer H, Bosche M, Drewes G, Freiberg GM, Kurnick MD, Srikumaran M, Kempf DJ, Green SR, Ray PC, Read K, Wyatt P, Barry CE, 3rd, Boshoff HI. Essential but Not Vulnerable: Indazole Sulfonamides Targeting Inosine Monophosphate Dehydrogenase as Potential Leads against *Mycobacterium tuberculosis*. ACS Infect Dis. 2017;3(1):18-33. Epub 20161017. doi: 10.1021/acsinfecdis.6b00103. PubMed PMID: 27704782; PMCID: PMC5972394.

S3. Wong AI, Rock JM. CRISPR Interference (CRISPRi) for Targeted Gene Silencing in Mycobacteria. Methods Mol Biol. 2021;2314:343-64. doi: 10.1007/978-1-0716-1460-0_16. PubMed PMID: 34235662.

S4. du Noüy PL. A new apparatus for measuring surface tension. J Gen Physiol 1919;1:521–4.

S5. Lee B-B, Chan E-S, Ravindra P, Khan TA. Surface tension of viscous biopolymer solutions measured using the du Nouy ring method and the drop weight methods. Polym Bull. 2012;69:471–89.

S6. Macy R. Surface tension by the ring method. J Chem Educ 1935;12:573.

S7. Walters K, Jones WM. Measurement of Viscosity. In: Boyes W, editor. Instrumentation Reference Book. 4th Edition ed: Butterworth-Heinemann; 2009.

S8. Ewoldt RH, Johnston MT, Caretta LM. Experimental Challenges of Shear Rheology: How to Avoid Bad Data. In: Spagnolie SE, editor. Complex Fluids in Biological Systems: Experiment, Theory, and Computation. New York, NY: Springer New York; 2015. p. 207-41.

S9. Johnston MT, Ewoldt RH. Precision rheometry: Surface tension effects on low-torque measurements in rotational rheometers. Journal of Rheology. 2013;57(6):1515-32. doi: 10.1122/1.4819914.

S10. Rodd LE, Scott TP, Boger DV, Cooper-White JJ, McKinley GH. The inertio-elastic planar entry flow of low-viscosity elastic fluids in micro-fabricated geometries. Journal of Non-Newtonian Fluid Mechanics. 2005;129(1):1-22. doi: https://doi.org/10.1016/j.jnnfm.2005.04.006.

S11. Soulages J, Oliveira MSN, Sousa PC, Alves MA, McKinley GH. Investigating the stability of viscoelastic stagnation flows in T-shaped microchannels. Journal of Non-Newtonian Fluid Mechanics. 2009;163(1):9-24. doi: https://doi.org/10.1016/j.jnnfm.2009.06.002.

S12. Oliveira MSN, Yeh R, McKinley GH. Iterated stretching, extensional rheology and formation of beads-on-a-string structures in polymer solutions. Journal of Non-Newtonian Fluid Mechanics. 2006;137(1):137-48. doi: https://doi.org/10.1016/j.jnnfm.2006.01.014.

S13. CRC Handbook of Chemistry and Physics, 85th Edition. Lide DR, editor: Taylor & Francis; 2004.

S14. Buck AL. New Equations for Computing Vapor Pressure and Enhancement Factor. Journal of Applied Meteorology and Climatology. 1981;20(12):1527-32. doi: https://doi.org/10.1175/1520-0450(1981)020<1527:NEFCVP>2.0.CO;2.

S15. Sobac B, Brutin D. Chapter 8 - Pure Diffusion. In: Brutin D, editor. Droplet Wetting and Evaporation. Oxford: Academic Press; 2015. p. 103-14.

S16. Popov YO. Evaporative deposition patterns: Spatial dimensions of the deposit. Physical Review E. 2005;71(3):036313. doi: 10.1103/PhysRevE.71.036313.

S17. Henderson-Sellers B. A new formula for latent heat of vaporization of water as a function of temperature. Quarterly Journal of the Royal Meteorological Society. 1984;110(466):1186-90. doi: https://doi.org/10.1002/qj.49711046626.

S18. Stull R. Wet-Bulb Temperature from Relative Humidity and Air Temperature. Journal of Applied Meteorology and Climatology. 2011;50(11):2267-9. doi: https://doi.org/10.1175/JAMC-D-11-0143.1.

S19. Deegan RD, Bakajin O, Dupont TF, Huber G, Nagel SR, Witten TA. Capillary flow as the cause of ring stains from dried liquid drops. Nature. 1997;389(6653):827-9. doi: 10.1038/39827.

S20. Garay-Arroyo A, Colmenero-Flores JM, Garciarrubio A, Covarrubias AA. Highly hydrophilic proteins in prokaryotes and eukaryotes are common during conditions of water deficit. J Biol Chem. 2000;275(8):5668-74. doi: 10.1074/jbc.275.8.5668. PubMed PMID: 10681550.

S21. Parker JM, Guo D, Hodges RS. New hydrophilicity scale derived from high-performance liquid chromatography peptide retention data: correlation of predicted surface residues with antigenicity and X-ray-derived accessible sites. Biochemistry. 1986;25(19):5425-32. doi: 10.1021/bi00367a013. PubMed PMID: 2430611.

S22. Cock PJ, Antao T, Chang JT, Chapman BA, Cox CJ, Dalke A, Friedberg I, Hamelryck T, Kauff F, Wilczynski B, de Hoon MJ. Biopython: freely available Python tools for computational molecular biology and bioinformatics. Bioinformatics. 2009;25(11):1422-3. Epub 20090320. doi: 10.1093/bioinformatics/btp163. PubMed PMID: 19304878; PMCID: PMC2682512.

S23. Ilzhofer D, Heinzinger M, Rost B. SETH predicts nuances of residue disorder from protein embeddings. Front Bioinform. 2022;2:1019597. Epub 20221010. doi: 10.3389/fbinf.2022.1019597. PubMed PMID: 36304335; PMCID: PMC9580958.

S24. SigmaSDS L-Arginine monohydrochloride for biochemistry [2024-07-23]. Available from: https://www.sigmaaldrich.com/US/en/sds/mm/1.01543?userType=anonymous.

S25. ChemicalBook L-Cysteine hydrochloride monohydrate [2024-07-09]. Available from: https://www.chemicalbook.com/ChemicalProductProperty_EN_CB9694981.htm.

S26. ChemicalBook L-CYSTINE DIHYDROCHLORIDE [2024-07-09]. Available from: https://www.chemicalbook.com/ChemicalProductProperty_EN_CB9147199.htm.

S27. Yaws CL. Chapter 5 - Density of Solid – Organic Compounds. In: Yaws CL, editor. Thermophysical Properties of Chemicals and Hydrocarbons (Second Edition). Oxford: Gulf Publishing Company; 2014. p. 264-352.

S28. SigmaSDS L-Lysine monohydrochloride [2023-07-23]. Available from: https://www.sigmaaldrich.com/US/en/sds/sial/l5626?userType=anonymous.

S29. ChemicalBook L-TYROSINE DISODIUM SALT [2024-07-09]. Available from: https://www.chemicalbook.com/CASEN_69847-45-6.htm.

S30. BOCScience Calcium D-Pantothenate [2024-07-09]. Available from: https://www.bocsci.com/product/calcium-d-pantothenate-cas-137-08-6-464900.html.

S31. ChemicalBook Pyridoxine hydrochloride [2024-07-09]. Available from: https://www.chemicalbook.com/ChemicalProductProperty_EN_CB8853882.htm.

S32. GuideChem Vitamin B12 [2024-07-09]. Available from: https://www.guidechem.com/dictionary/en/68-19-9.html.

S33. PubChem Monosodium phosphate [2024-07-09]. Available from: https://pubchem.ncbi.nlm.nih.gov/compound/Monosodium-phosphate.

S34. ChemSpider 2'-Deoxyadenosine [2024-07-09]. Available from: https://www.chemspider.com/Chemical-Structure.13135.html.

S35. ChemSpider 2'-Deoxycytidine [2024-07-09]. Available from: https://www.chemspider.com/Chemical-Structure.13117.html.

S36. ChemSpider 2'-Deoxyguanosine [2024-07-09]. Available from: https://www.chemspider.com/Chemical-Structure.163230.html.

S37. ChemSpider lipoic acid [2024-07-09]. Available from: https://www.chemspider.com/Chemical-Structure.841.html.

S38. MilliporeSigma Sodium pyruvate [2024-07-09]. Available from: https://www.sigmaaldrich.com/US/en/product/sigma/p5280.

39. ChemicalBook N,N-Dimethylglycine hydrochloride [2024-07-09]. Available from: https://www.chemicalbook.com/ChemicalProductProperty_EN_CB8360829.htm.

S40. eChemi DL-Isocitric acid, trisodium salt hydrate [2024-07-09]. Available from: https://www.echemi.com/produce/pr220729208495-dl-isocitric-acid-trisodium-salt-hydrate.html.

S41. MiliPoreSigma Ethanolamine hydrochloride [2024-07-09]. Available from: https://www.sigmaaldrich.com/US/en/product/sial/e6133.

S42. ChemSpider gamma-Aminobutyric acid [2024-07-09]. Available from: http://www.chemspider.com/Chemical-Structure.116.html.

S43. Lactic Acid, Sodium Lactate, and Potassium Lactate [2024-07-09]. Available from: https://www.ams.usda.gov/sites/default/files/media/Lactic%20Acid%20TR%202015.pdf.

S44. CarlRoth L-Ornithine monohydrochloride [2024-07-09]. Available from https://www.carlroth.com/com/en/ornithine/l-ornithine-monohydrochloride/p/t204.1.

S45. Bronzino JD, Peterson DR. Biomedical engineering fundamentals: CRC press; 2014.

S46. Oikonen V. Bloody PET [2024-07-25]. Available from: http://www.turkupetcentre.net/petanalysis/blood.html.

S47. Nagle JF, Wilkinson DA. Lecithin bilayers. Density measurement and molecular interactions. Biophysical Journal. 1978;23(2):159-75. doi: https://doi.org/10.1016/S0006-3495(78)85441-1.

S48. ChemSpider Triolein [2024-07-09]. Available from: https://beta.chemspider.com/Chemical-Structure.4593733.html.

S49. ChemSpider 1,3-Dioleoyl-2-palmitoylglycerol [2024-07-09]. Available from: https://beta.chemspider.com/Chemical-Structure.4720768.html.

S50. ChemSpider cholesteryl linoleate [2024-07-09]. Available from: https://beta.chemspider.com/Chemical-Structure.5021035.html.

S51. BOSScience C24 Lactosyl(beta) Ceramide (d18:1/24:0) [2024-07-09]. Available from: https://liposomes.bocsci.com/product/c24-lactosyl-beta-ceramide-d18-1-24-0-cas-105087-85-2-363136.html?nid=6648.

S52. ChemSpider C16 Lactosylceramide (d18:1/16:0) [2024-07-09]. Available from: https://beta.chemspider.com/Chemical-Structure.24765743.html.

S53. ChemSpider Deoxysphinganine [2024-07-09]. Available from: https://www.chemspider.com/Chemical-Structure.8101521.html.

S54. Schildkraut CL, Marmur J, Doty P. Determination of the base composition of deoxyribonucleic acid from its buoyant density in CsCl. Journal of Molecular Biology. 1962;4(6):430-43. doi: https://doi.org/10.1016/S0022-2836(62)80100-4.
